# Aperiodic and Periodic EEG Component Lifespan Trajectories: Monotonic Decrease versus Growth-then-Decline

**DOI:** 10.1101/2025.08.26.672407

**Authors:** Min Li, Ying Wang, Yaqi Chen, Adrien E. E. Dubois, Gangyong Jia, Qing Wu, Maria L. Bringas-Vega, Guillaume Dumas, Pedro A. Valdes-Sosab

## Abstract

1.1

Unraveling the lifespan trajectories of human brain development is critical for understanding brain health and disease. Recent research demonstrates that electroencephalography signals are composed of periodic and aperiodic components reflecting distinct physiological substrates. This dissociation raises the possibility that they follow different developmental tendencies. Here, we delineate the lifespan trajectories of aperiodic and periodic neural oscillations using a large international cohort (N=1,563, ages 5–95, resting state, eyes closed). We reveal two fundamental developmental patterns: a Monotonic decrease in aperiodic activity and a Growth-and-Decline pattern for periodic activity. Both components have inflections around age 20 and transition to a stable senescent phase around age 40. Spatially, anterior regions mainly exhibit aperiodic activity, while periodic activity concentrate on posterior regions and these patterns remain stable throughout life. Crucially, multimodal analysis shows these trajectories map onto distinct biological substrates. The periodic component’s Growth and Decline trajectory aligns with GABAergic function and myelination. In contrast, the monotonically decreasing trajectory of aperiodic activity mirrors fundamental biomarkers of biological aging, such as DNA methylation and telomere length. Transforming age to a logarithmic scale simplifies these nonlinear trajectories into a linear decreasing and a piecewise concave linear model for aperiodic and periodic components. This form provides a robust and parsimonious framework for quantifying maturation and identifying neurological deviations.

**Highlights:**

1. We delineate distinct lifespan trajectories of aperiodic and periodic neural activity in a large-scale international cohort (N=1,563, ages 5–95). Aperiodic activity undergoes a Monotonic Decrease with age. In contrast, periodic activity follows a Growth-then-Decline trajectory, peaking in early adulthood.
2. Both trajectories feature a critical transition around age 20 and stabilize into a protracted senescent phase from approximately 40 onward.
3. These neural trajectories map onto distinct biological substrates: periodic activity tracks integrative functions (myelination, GABAergic, and aperiodic decline mirrors fundamental aging processes (DNA methylation).
4. A stable pattern observed throughout the lifespan is the spatial segregation of neural activity, where aperiodic signals are dominant in anterior regions and periodic signals are concentrated in posterior ones.
5. Logarithmically transforming age linearized the developmental trajectories, yielding a monotonic decline for the aperiodic component and a concave piecewise for the periodic one. This process establishes robust linear norms for the personalized assessment of brain dysfunction.

## 1.2 Introduction

We investigate the lifespan trajectories of the human electroencephalographic power spectral density (EEG-PSD) components, addressing the need for temporally resolved, scalable neuroimaging markers in developmental research. EEG offers complementary advantages to modalities such as MRI, including enhanced temporal resolution and lower implementation cost.

Emerging normative studies reveal that several brain measures, such as total white matter volume (WMV), follow a “Growth-then-Decline” curve—rapid expansion from mid-gestation to early adulthood, peaking at 28.7 years, followed by a steady decline over the next five decades (R. A. I. Bethlehem et al., 2022). Similar patterns are observed in cognitive development (McArdle et al., 2002), hormonal changes, and neurodegenerative processes (Cole et al., 2018). While MRI-based approaches dominate, their scalability and accessibility limitations underscore the value of EEG for population-level studies (Thompson et al., 2020).

EEG-PSD studies frequently report analogous growth and decline patterns. For instance, the alpha (8–13 Hz) central frequency increases from childhood to adulthood and diminishes in later life (Szava et al., 1994; Li et al., 2022a). Studies across infancy (Wilkinson et al., 2024), adolescence, adulthood (Rempe et al., 2023; Tröndle et al., 2022), and aging (Voytek et al., 2015) confirm this bidirectional maturation. Notably, not all biological indicators follow this trajectory; DNA methylation and telomere length demonstrate monotonic age-related changes (Hannum et al., 2013; Horvath, 2013; Jones et al., 2015), forming the basis for biological clocks and research into aging and neurodegeneration(Forero et al., 2016; Jylhävä et al., 2017; Fani et al., 2020; Seale et al., 2024). These results suggests brain development may follow at least two distinct temporal profiles.

Our hypothesis, which drives the stated purpose of our study, is that aggregate EEG-PSD analyses may conceal the coexistence of these developmental trajectories. Focusing on two separable PSD components—aperiodic (xi process) and periodic (alpha process) (Pascual et al., 1988)—we note that the broadband xi process resembles pink noise, while the alpha process manifests as a spectral peak, corresponding to harmonic oscillations. Contemporary models (e.g., FOOOF) support these decompositions. Evidence indicates that these processes reflect distinct physiological mechanisms (Valdés et al., 1992; He, 2014; Pascual-Marqui et al., 2022; Kramer and Chu, 2023; Ouyang et al., 2020; Ronaldo, Wang), and their developmental patterns differ in brain disorders and cognitive impairments (Klimesch, 2012; Gerster et al., 2022; Wilson et al., 2022; Albertson et al., 2024).

Prior work has addressed age-related shifts in these components. Amador et al. (Amador et al., 1989) observed linear changes in the amplitude of xi and alpha processes from ages 5–12, with the former decreasing and the latter increasing. Recent findings show older adults exhibit flatter EEG-PSD compared to middle-aged individuals (Voytek et al., 2015), and accounting for aperiodic contributions enhances the age-correlation of periodic (alpha) activity (Tröndle et al., 2022). However, most studies are constrained by narrow age ranges or focus on specific frequency bands.

Leroy et al. (Leroy et al., 2025) provide the first broad age-range analysis (8–92 years, n=532) of EEG periodic and aperiodic activity, reporting only linear age associations in alpha and gamma bands, and no significant changes in aperiodic activity. These outcomes may reflect sample heterogeneity and limitations in spectral parameterization.

Therefore, our central questions are: What are the lifespan trajectories of periodic and aperiodic EEG components? How can these be mathematically described and spatially mapped? How do EEG trends compare with trajectories from other biological modalities?

To address these, we generated nonlinear normative curves for each EEG component using 1,563 EEG recordings (781 males, 782 females, resting state, eyes closed) from 14 studies across 9 countries (HarMNqEEG/Global Brain Consortium, globalbrainconsortium.org) (Figure 1a). We integrated multimodal age metrics for comparative analysis (Figure 1b)—including MRI, myelination, cognitive scores, DNA methylation, telomere length, metabolism, and water turnover (see Data Available). Through preprocessing (check in method 1.1.3-1.1.4), batch harmonization (method 1.1.5) and component analysis (method 1.1.6), the spatial characteristics of trajectories were explored, and resulting in a simplified linear model for classifying populations with multiple mental disorders finally (Figure 1-c). This endeavor constitutes a foundational step toward systematic and quantitative EEG analysis. It not only provides a basis for the physiological and biophysical interpretation of each component but also offers a crucial reference for accurately distinguishing between normal and abnormal patterns of brain development, maturation, and aging.

**Figure 1:**
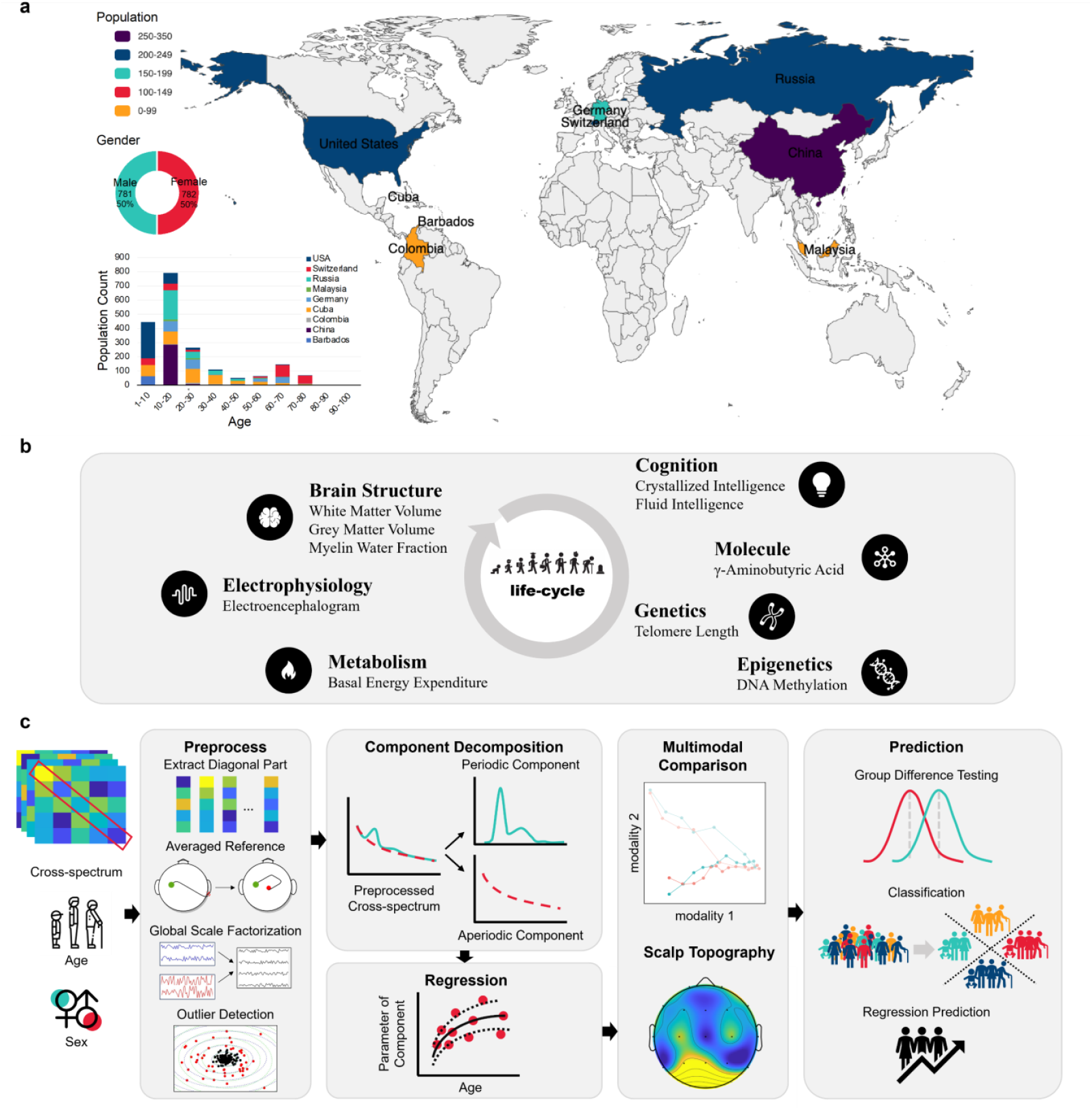
Dataset, Multimodal Comparison, and Analytical Workflow. a: Lifespan EEG cross-spectra were obtained from the HarMNqEEG project (https://globalbrainconsortium.org/project-eegNorms/), comprising 1,563 healthy participants (781 M, 783 F) recorded during eyes-closed rest. The data were aggregated by the Global Brain Consortium (GBC, https://globalbrainconsortium.org/about/) from 14 studies across 9 countries and 12 devices via a decentralized, code-based protocol that avoided raw data collecting. Centralized preprocessing involved interpolation to a standardized 19-channel 10-20 montage (Dong et al., 2021), average referencing, global scale factorization (GSF), outlier detection, batch harmonization (see details of preprocessing in SI 1.1.1-1.1.5 (Li et al., 2022)). b: EEG-derived developmental trajectories were benchmarked against normative data from other modalities, including brain structure and cognitive ability. c: The analytical pipeline involved three primary stages: (i) periodic and aperiodic components were extracted using a robust in-house toolbox (see Methods 1.1.6); (ii) age-related trajectories (mean and standard deviation) were modeled using locally weighted kernel regression (LOWESS, statsmodels library, https://www.statsmodels.org/stable/index.html)); and (iii) these normative models were used to develop a simplified linear classifier for distinguishing individuals with various mental disorders.

## 1.3 Results

### 1.3.1 Nonlinear Lifespan Trajectories of Aperiodic and Periodic Components

Grounded in biophysical models (Kramer and Chu, 2023; Brake et al., 2024) and the physiological properties of cortical signal generation (Mendoza-Halliday et al., 2022), aperiodic and periodic signals are conceptualized as independent. We, therefore, constructed an additive model of these components on a natural scale (Brillinger, 2001). We employed our in-house spectral parameterization toolbox (see Method 1.16) to decompose the EEG-PSD. The aperiodic component was modeled with a Lorentzian function, and periodic peaks were modeled with Gaussian kernels. This model yielded quantitative parameters for the aperiodic component-Offset and Exponent and the periodic component-Center Frequency [CF], Power [PW], and Bandwidth [BW]) (Figure 2a). Guided by the age-energy distribution pattern observed in the log-spectral developmental surface (see SI-2.1, Figure S1), and considering that children tend to exhibit lower-frequency alpha peaks while adults have higher-frequency alpha components, we adjusted the frequency bands as follows: 1–4 Hz (delta), 4–6 Hz (theta), 6–14 Hz (alpha), and 14–20 Hz (beta). Please note that the term ‘periodic activity’ throughout results in this study refers specifically to the aperiodic-corrected periodic activity.

**Figure 2:**
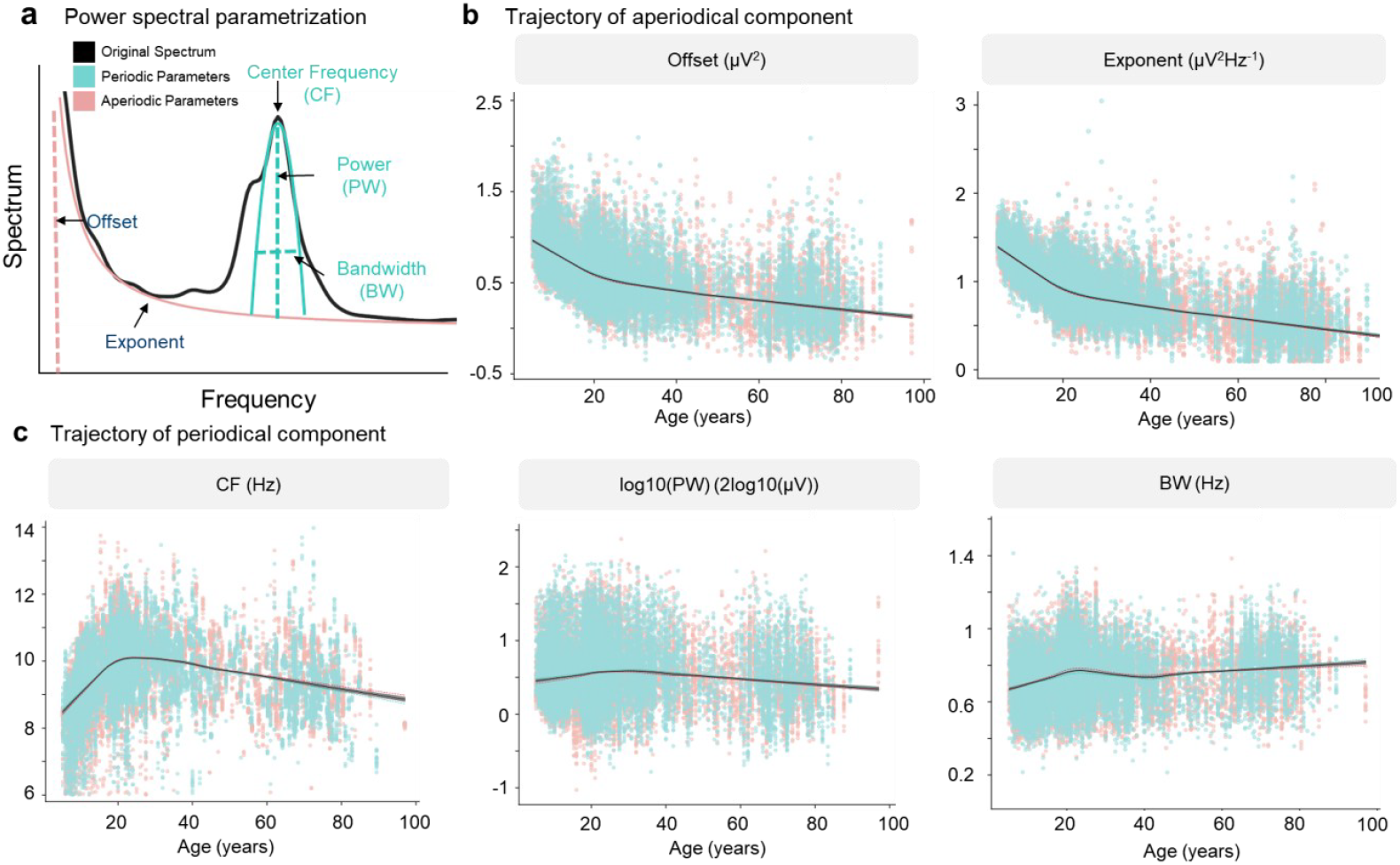
The developmental trajectory of aperiodic and periodic components. a: The EEG spectrum can be parameterized into aperiodic and periodic components using the Lorentzian function and Gaussian kernel, with parameters including Exponent and Offset for the aperiodic component, and CF, PW, and BW for the periodic component; b: The developmental trend of aperiodic component parameters with age. c: The developmental trend of periodic component parameters with age. Pink represents females, blue represents males, black represents the entire group, and the gray shading represents the confidence interval (CI) (Davison and Hinkley, 1997) of the overall regression. The regression and CI calculation details can be checked in Method 1.1.7. The changes in EEG component parameters with age are not linear but follow two distinct developmental trends: a growth-then-decrease pattern for alpha and a continuous decline for xi.

As shown in Figure 2, nonlinear kernel regression (LOWESS) across the whole lifespan (ages 5–95) and all brain regions revealed two predominant and fundamentally distinct developmental trends. This nonlinearity was evident across parameters R^2^ (e.g., for alpha-band CF, nonlinear R^2^ = 0.22 versus linear R^2^ = 0; Table S2). The periodic component, particularly in the alpha band, traced a canonical Growth-then-Decline trajectory (Fig. 2c). Both CF and PW increased from childhood (ages 5–20) to a peak in the early twenties, subsequently declining throughout adulthood, mirroring the pattern reported in other neuroimaging studies (Yeatman et al., 2014; Hill et al., 2022). Bandwidth (BW) follows a similar developmental trajectory to CF and PW before age 40; however, after 40 years of age, BW shows a slight increase rather than a decline.

In low-frequency bands (1–4 Hz and 4–6 Hz), adults exhibit fewer detectable periodic components compared to children, as indicated by sparser CF estimates. Moreover, PW in these bands shows an inverse developmental trend relative to the alpha band: it decreases during childhood and after the inflection point at about 20 years old it increase (Figure. S6 a–f). This suggests that children have lower-frequency rhythmic activity than adults. In contrast, adults show stronger periodic components at the high-frequency band (14–20 Hz) than children, with all parameters—CF, PW, and BW—increasing with age which is consistent with the result reported in paper (W. He et al., 2019) that the Beta activities is increase with age (Figure S6 g–i).

In stark contrast, the aperiodic component parameters—Offset and Exponent—show a rapid decline from childhood to early adulthood (ages 5–20), followed by a more gradual decrease beyond age 20 (Figure. 2b). The observed trend indicates that elderly individuals exhibit a flatter 1/f slope compared to children and younger adults, a pattern consistent with previously reported age-related changes in aperiodic activity.(Voytek et al., 2015; W. He et al., 2019; Donoghue et al., 2020; Pathania et al., 2022; Hill et al., 2022; Bender et al., 2023; Favaro et al., 2023).

To investigate potential sex differences in the developmental trajectories, we performed separate model fittings for male and female participants, comparing both mean values and standard deviations. The age-related changes in both periodic and aperiodic EEG components showed no significant differences between sexes (Offset: *p* = 8.16 × 10^−17; Exponent: *p* = 1.12 × 10^−14; CF_alpha: *p* = 9.22 × 10^−28; PW_alpha: *p* = 0; BW_alpha: *p* = 0 at α = 0.05), consistent with previous findings from Li et al., 2022 (Li et al., 2022a), who also reported no sex differences in the developmental patterns of EEG-PSD using cross-validated modeling approaches. Nevertheless, However, the conclusion of no sex differences is not consistent with previous findings (Cave and Barry, 2021; McSweeney et al., 2021), which does not preclude variations in results due to differences in sample geographic distribution, ethnic composition, and other factors. Beyond the question of sex differences in developmental trajectories, we also assessed the degree of sample deviation in these trajectories. Through age-related modeling of parameter variance, we observed no substantial change across development except for a modest increase around age 20. (Figure S7).

To verify that these two patterns represent fundamental organizing principles of neurodevelopment, we performed a Principal Component Analysis (PCA) on the full electrode-by-frequency data matrix. The first three components, accounting for 65% of total variance (Figure S8 d), mapped cleanly onto our hypothesized trajectories. PC1 closely tracked the Monotonic Decrease of the aperiodic component (Figure S8 a). At the same time, PC2 mirrored the Growth-then-Decline of the periodic alpha-band component (Figure. S8 b). At the same time, PC3 reflects a composite pattern of the remaining periodic developmental trajectories (Figure S8 c). Examination of the component loadings revealed that specific frequency bins—not individual channels—dominate the contributions to each principal component (Figure S8 e-f). This data-driven analysis provides robust, independent evidence for the biological salience of these two dissociable developmental modes.

Since EEG activity under resting-state, eyes-closed conditions is predominantly concentrated in the alpha band, the following characterization of developmental trajectories will focus on the aperiodic parameters and the periodic parameters within the alpha band.

### 1.3.2 Ages 20 and 40 Demarcate Critical Developmental Transitions

To delineate the temporal dynamics of neurodevelopment, we computed the first (velocity) and second (acceleration) derivatives of the fitted aperiodic and periodic trajectories. As shown in Figure 3 and Figure. S9, both Exponent and Offset exhibit consistently negative rates of change across the lifespan (first derivative < 0, solid black line). However, the decline decelerates after age 20, which serves as an inflection point coinciding with the second derivative’s peak—the maximum deceleration rate (solid blue line). By approximately age 44, the second derivative approaches zero, indicating the onset of a sustained aging phase characterized by a constant, linear decline (Figure 3a, Figure S8a).

**Figure 3:**
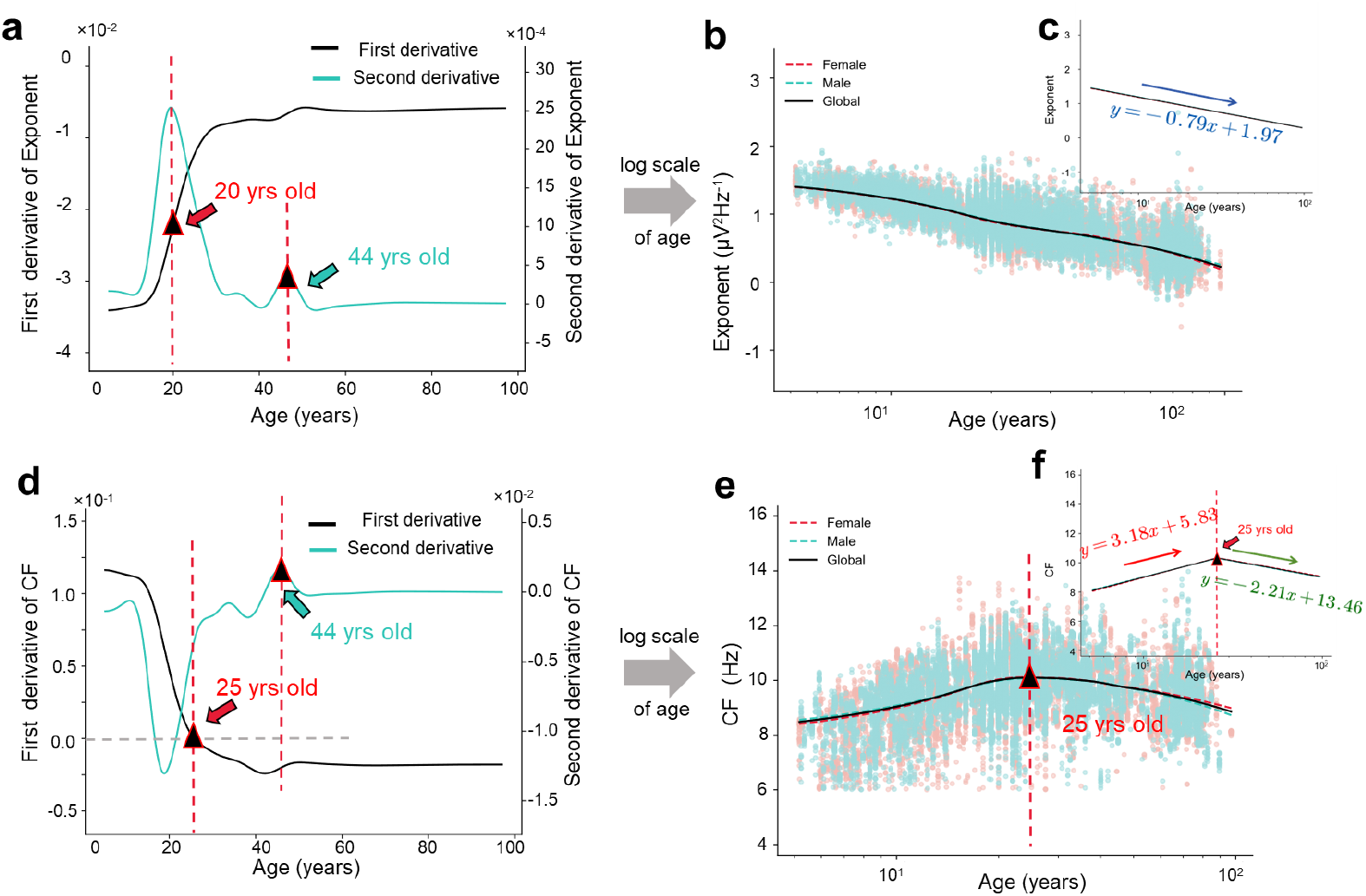
Key Developmental Inflection Points and Log-Scaled Trajectories. a: For the aperiodic Exponent, age 20 represents a critical inflection point in the rate of change, first coinciding with the peak of the second derivative. After this point, the developmental speed gradually declines until age 44, when the second derivative reaches zero, indicating the onset of a stable, adult-like state.; b: On the log(age) scale, the Exponent shows a Monotonic Decrease across the lifespan, (c) which is well captured by a simple linear model with 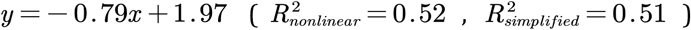. d: Age 25 marks the CF development, where the first derivative equals zero. By age 44, CF enters a plateau phase, as indicated by a second derivative approaching zero; e: When plotted on a log(age) scale, the CF trajectory exhibits an approximately symmetric, inverted-U shape centered around age 25. This nonlinear pattern can be approximated by a piecewise linear model (f), defined by two distinct developmental phases (*y* = 3.18*x* + 5.83 and 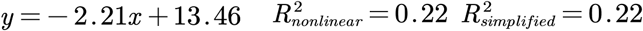). The first derivation is colored with a solid black line, and the second is colored with a solid blue line. Check Figure S9 for Offset and PW.

For periodic components, the first derivative (solid black line) of CF and PW gradually declines from age 5 through early adulthood (ages 20–25), remaining positive until approximately age 25 (30 for PW). Around this point, the rate crosses zero, marking the peak of development. Thereafter, the rate becomes negative and stabilizes around age 40—indicating a transition to a phase of steady aging with no further acceleration (second derivative ≈ 0, solid blue line) (Figure 3d and Figure S9d)

These findings identify the early twenties and early forties as key developmental milestones—marking the transition from rapid, nonlinear maturation to a slower, more stable aging phase. This temporal pattern aligns with observations from other neuroimaging modalities. It supports a broader model of brain development governed by distinct phases of dynamic change and stabilization (Liu et al., 2024; Shen et al., 2024).

Based on the logarithmic development progression, where 23 years is the approximate median of the 5–95 year range, we replotted parameter trajectories on a base-10 logarithmic age scale. Figure 3 shows a near-linear Monotonic Decrease in the aperiodic Exponent (Figure 3b) and an approximately symmetric Growth-then-Decline path centered on the inflection point for the periodic parameter (Figure 3e). The monotonic and symmetric patterns in the log age scale of these parameters enable the nonlinear developmental trajectories to be effectively approximated by a linear and piecewise linear model (Figure 5c,f; Figure S9c, f). Since the rate of change is variable before the age around 40, such, to achieve the greatest possible simplification of the non-linear model, here only consider the first inflection point to generate the linear and piecewise linear model. The Exponent could be modeled with a linear function 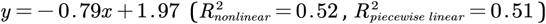 (Figure 3c). CF could be described with a piecewise linear function and *y* = 3.18*x* + 5.83 and 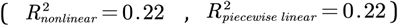 (Figure 3f) (for other parameters check in SI 3.5,). Compared to non-linear models, the log(age) piecewise linear functions provide a new type of developmental norms for neuro oscillations, yielding more robust parameter estimates than non-linear models without sacrificing accuracy (check R^2^ in SI Table 2). The piecewise linear models render brain age prediction tractable (which can do brain age prediction by dividing into two age geoups), and the linear trends of aperiodic parameters underscore their potential as reliable biomarkers of biological age. Their stability across development and wide dynamic range place them on par with established molecular markers such as telomere length and DNA methylation, thereby providing a functional electrophysiological correlation of aging.

### 1.3.3 Topographical Organization is Distinct but Spatially Stable over the lifespan

Research has suggested that aperiodic components reflect background neural activity associated with short-range connectivity (Ibarra Chaoul and Siegel, 2021; Cross et al., 2024), whereas periodic components are thought to index widespread, long-range neuronal synchronization, including cross-frequency coupling (Pascual-marqui et al., 1988; Valdés et al., 1992; Mendoza-Halliday et al., 2022). However, age-related changes in the spatial distribution of these components across the scalp remain uncharacterized.

This study investigated the spatial distribution of periodic and aperiodic component parameters and their developmental trajectories for different electrode on the scalp. During resting-state eyes-closed conditions, at key developmental inflection points, we observed prominent alpha-band periodic activity of CF and PW in the occipital lobe relative to other brain regions in alpha band. BW showed elevated values in both the occipital and sensorimotor regions (Figure 4a; see Figure S10 for other periodic bands). In contrast, the aperiodic exponent was most pronounced in the frontal lobe. At the same time, the offset exhibited higher values in both frontal and occipital areas. The topographical results of the aperiodic parameters in our study differ from those reported by Hill et al. (2022)(Hill et al., 2022), who found that the exponent was concentrated in the parietal and temporal lobes, while the offset was concentrated in the occipital lobe with sample of children aged 4-12. However, their finding is consistent with the results from the 5-7 year-old group in this study. This discrepancy may be caused by age-related biases resulting from factors such as geographical sampling differences.

**Figure 4:**
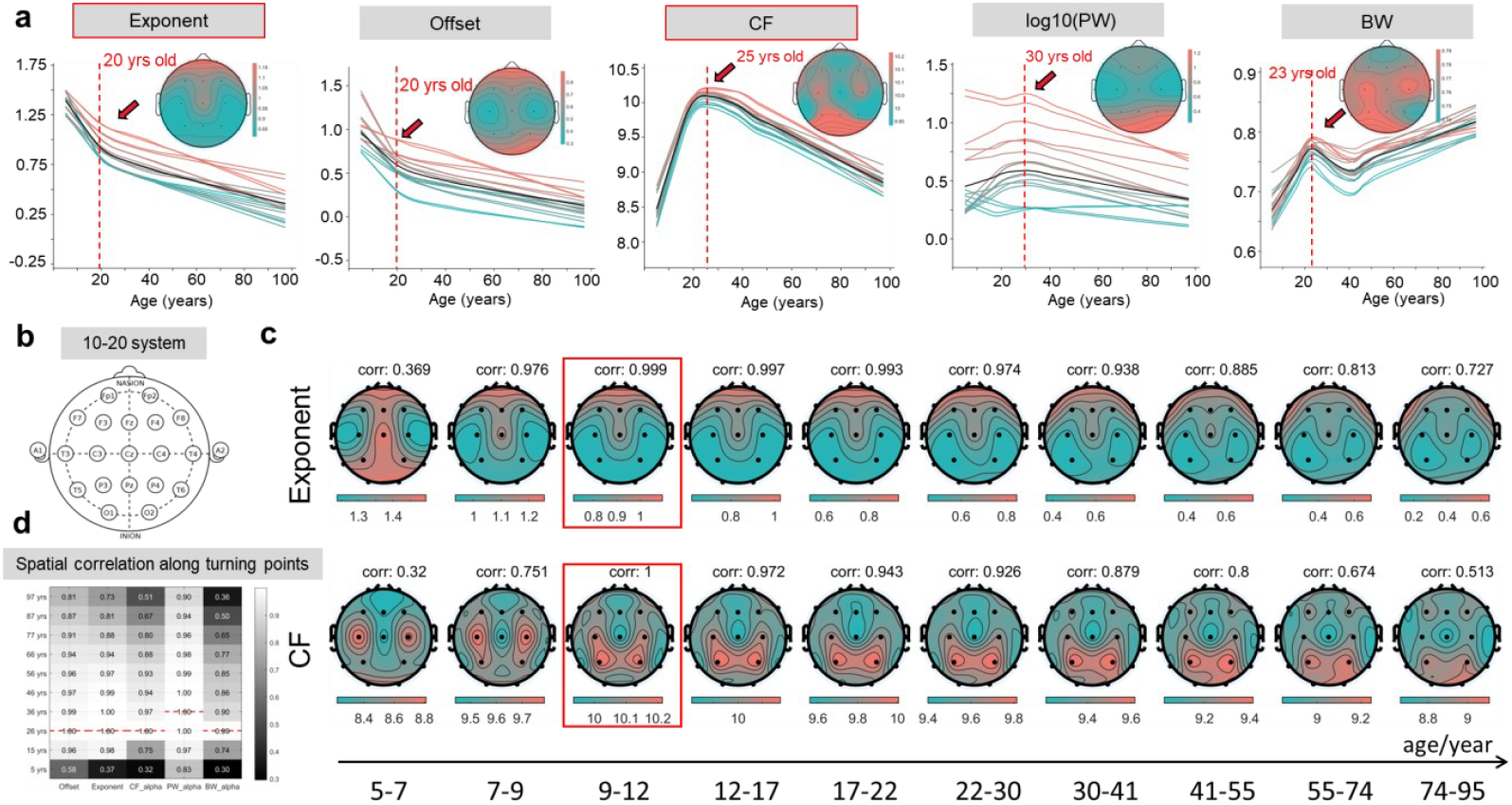
The spatial distribution of the scalp EEG spectral parameters. a: The spatial distribution of aperiodic and periodic parameters at developmental inflection points (topography plot) and its developmental tendency of all 18 channels (solid line, colored according to the topography plot). b: EEG signals were recorded according to the 19 electrodes international 10–20 electrode placement system; c: The developmental tendency of spectral parameters’ spatial distribution. The topographies show mean values at sampled ages. Correlation values measure the spatial consistency of topographies at different ages with the distribution at inflection points (red circle). D: The spatial distribution correlation values of aperiodic and periodic of the alpha band. The dashed line inside the matrix means the correlation baseline, corresponding to the inflection points. The most significant correlation values are labeled with a dashed red line.

**Figure 5:**
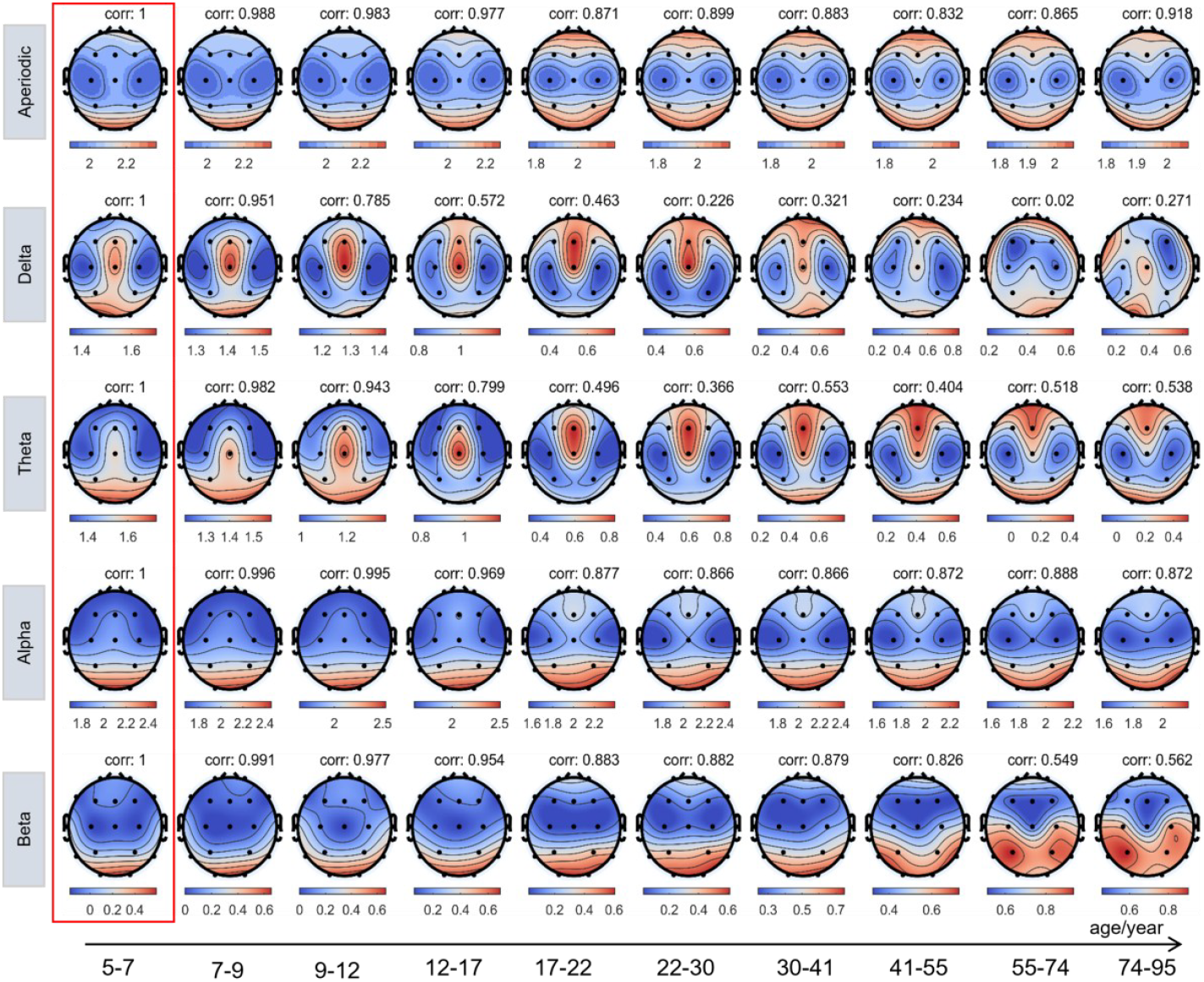
EEG energy spatial distribution of different components. The energy spatial distribution on the scalp, each value is measured with mean values in a specific age range (the age ranges are under the arrow), and the topography is colored within the local scale. Whereas the scalp topographies of power in the alpha and beta bands are developmentally stable, the distributions of the aperiodic component and the lower-frequency delta and theta components undergo a subtle shift around age 17. The results can be confirmed with the correlation values above each topography, estimated against the age group of 5-7 years old. The age group is divided on a log scale.

The parameter trends across all electrodes are largely consistent for whole-brain developmental trajectories. Notably, the developmental trajectory of PW displayed greater regional variability than CF (*CF*_*volatilit*_ = 5%, *PW*_*volatility*_ = 67%, *volatility* = (max | *y*_*i*_ − *y*_*j*_ |/max | *y*_*k*_ | × 100%), indicating that amplitude-related features are less spatially uniform and more regionally specific than frequency-based characteristics (check Figure S12 for parameter developmental trajectory across different functional regions). The spatial distribution was remarkably stable across the entire lifespan. Spatial correlation maps computed against the key developmental inflection points (labeled with a red box) remained high (r > 0.5) throughout (Figure 4c-d; Figure S11 for other parameters). Although the magnitude of parameters changed significantly with age, their underlying topographical organization appears to be a core, invariant feature of the brain’s functional architecture.

### 1.3.4 Energy distribution of EEG aperiodic and periodic activity

Although the spatial distribution of the total EEG PSD is well-characterized at the scalp (Nunez and Srinivasan, 2006), the distinct topographies of its aperiodic and periodic components’ energy which involved in both parameters —and how they evolve with age — remain largely unexplored.

To delineate the spatial dynamics of neural activity across the lifespan, we quantified periodic and aperiodic energy, which was estimated by computing the area under the kernel regression curve for discrete age bands and projecting their mean distribution onto scalp topographies. The degree of spatial reorganization over time was then formally assessed by correlating the topography of each age group against a baseline reference established from the youngest cohort (ages 5–7). These correlation coefficients, indicating the stability of the spatial pattern relative to early childhood, are annotated above each corresponding map. As shown in Figure 5, Aperiodic energy, initially concentrated in occipital regions during childhood, undergoes a significant spatial reorganization. It exhibits a decisive shift toward prefrontal dominance around age 20, a localization that subsequently stabilizes and persists into late adulthood.

In contrast, periodic components displayed frequency-specific patterns of spatial maturation. Low-frequency (delta and theta) power demonstrated the most dynamic trajectory. Initially localized to occipital-central areas, it commences a pronounced anterior shift around age 17, eventually diffusing into a more distributed fronto-occipital pattern by age 40 (evident in Figure S13a with global scale). In stark contrast, alpha-band power maintained a highly stable spatial profile, consistent and robust dominance in the occipital cortex throughout the lifespan (spatial correlation > 0.8). Beta activity, while also rooted in occipital regions, showed a gradual expansion into temporal areas with increasing age (Fig. 5b). Notably, these evolving topographies align closely with normative EEG power maps (Fig. S14a) and the age-dependent spatial contributions of the aperiodic component (Fig. S14 b).

### 1.3.5 Multimodal Correlates: Dissociable Trajectories Map onto Distinct Biological Systems

Brain maturation, development, and aging are influenced by various factors, including gene expression, neurotransmitter levels, energy metabolism, and oxidative processes (Frey and Morris, 1997; Flavell and Greenberg, 2008; Mendoza-Halliday et al., 2022; Buzsáki, 2010). These changes are reflected in the synchronized activity of large neuronal populations. For instance, alterations in gene expression can impact neuronal function and synaptic connectivity, manifesting as changes in brain electrical activity patterns (Buzsáki and Wang, 2012; Buzsáki and Mizuseki, 2014). Previous studies have shown that both periodic and aperiodic EEG components contribute to cognitive development and are implicated in neurodegenerative and psychiatric disorders (Colombo et al., 2019; Robertson et al., 2019; Merkin et al., 2023a). However, the physiological basis and potential of these components as clinical biomarkers remain unclear. Specifically, what roles do periodic and aperiodic components play, and how do they contribute to brain development? Addressing these questions requires integrating insights from other brain development models and conducting comparative analyses across multiple modalities.

To elucidate the physiological significance of the developmental trajectories of periodic and aperiodic EEG components, we compared our derived curves with established physiological and phenotypic parameters spanning the human lifespan. Specifically, we analyzed the developmental patterns of higher-level neural system expressions, including neurotransmitter levels, energy metabolism, brain structural imaging, and cognitive performance. We also examined primary regulatory trajectories, such as DNA methylation levels and telomere length, to uncover the mechanisms underpinning the development of periodic and aperiodic neuronal signals in the brain. This integrative approach aims to comprehensively understand the physiological processes that shape these distinct EEG components throughout life.

As shown in Figure 6, we compared EEG spectral features with the developmental trajectories of gamma-aminobutyric acid (GABA) levels, basal energy expenditure (BEE), water turnover (WT), total white matter volume (WMV), total cortical grey matter volume (GMV), functional health of white matter, and cognitive performance. The results reveal that these parameters follow developmental patterns closely resembling those of EEG periodic components, showing an overall increase from childhood until around 20, followed by a gradual decline over the subsequent seven decades. Notably, GMV exhibits a distinct trajectory: it rises steadily from mid-gestation, peaks at approximately 5.9 years of age, and then declines essentially linearly. This pattern diverges from that of periodic components, potentially reflecting the predominant role of gray matter in short-range neural processing. Further comparative analyses by adjusting GMV are warranted to explore the trajectory consistency (Boord et al., 2007).

**Figure 6:**
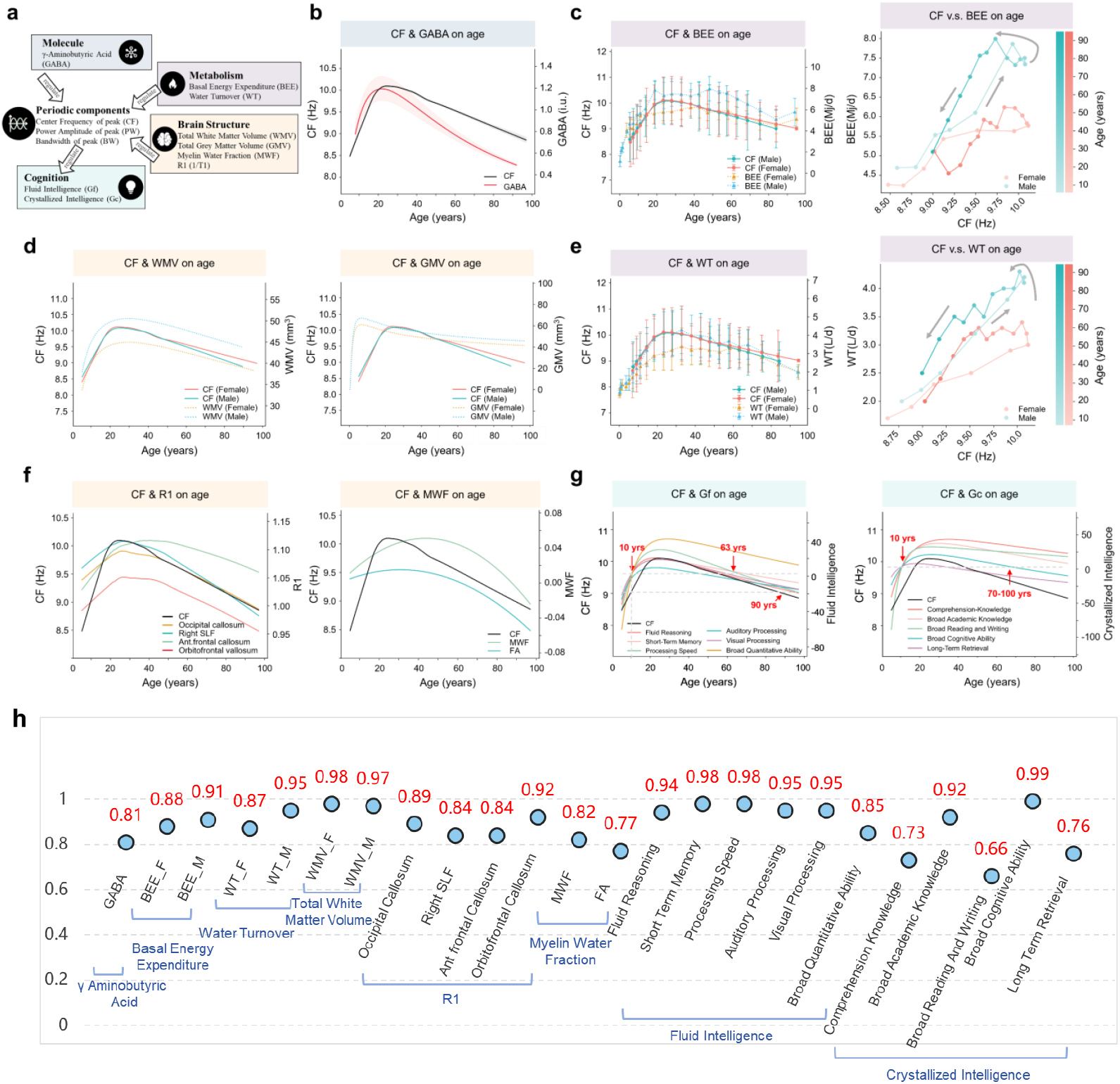
The relation of the developmental trajectory of CF with other modalities. a: Summary of all the modalities used. b-g: The developmental trajectory of CF is consistent with b-molecule feature-γ-Aminobutyric acid (GABA), c and e: metabolism features of basal energy expenditure (BEE), water turnover (WT). The standard deviation is labeled here for BEE and WT, which are directly extracted from the papers of Pontzer et al., 2021 and Yamada et al. (2022. d: brain structure features of total brain white matter volume (WMV), total gray matter volume (GMV), and f: Myelin health feature of R1 and myelin water fraction (MWF)), and intellectual ability including fluid intelligence (Gf), crystallized intelligence (Gc). The right panels of c and e display the progression of CF with BEE and WT over the life course, illustrating how BEE and WT vary with CF (color gradients indicate age, from light to dark; pink represents females and blue represents males). The gray arrow indicates the direction of increasing age. h: The correlation values between CF and other modalities.

Here, we first assessed white matter integrity using the longitudinal relaxation rate R1 (R1=1/T1, Figure 6 f left), which quantifies the rate at which tissue regains longitudinal magnetization in an external magnetic field (the curve obtained here differs from the one derived using a quadratic function in the article; detailed discussion can be found in SI 6.3). However, R1 serves only as an indirect marker of myelin health; reductions in extracellular water content within a voxel can also elevate R1 values (Faulkner et al., 2024; MacKay and Laule, 2016a). Therefore, we further examined the developmental trajectory of myelin water fraction (WMF) (Figure 6f, right). However, the results showed limited alignment with the developmental trends of WMV, CF, and PW. MWF peaked around 40 years of age, a finding likely influenced by several factors: 1) The available MWF dataset spanned ages 21–94 years, leaving early developmental changes (<20 years) uncharacterized; 2) MWF trends varied significantly across different brain regions (here show the MWF value of whole brain), such as the corona radiata (CR) and thalamic radiation (TR), where no significant decline was observed before the age of 40 (check Kiely et al., 2022 (Kiely et al., 2022a) in detail); 3) The curve was fitted using a polynomial model (Pi = β_0_ + β_sex_ × sex_i_ + β_a_age × age_i_ + β_a_ age ^2^ × age_i_^2^) rather than a non-parametric fitting method, which may be sensitive to the choice of basis functions compared to non-parametric fitting approaches. These limitations highlight the need for future studies employing more comprehensive datasets spanning the whole lifespan and utilizing nonlinear, non-parametric modeling techniques to characterize myelin development better.

Regarding cognitive development, we observe the following (Figure 7): 1) different dimensions of crystallized intelligence (Gc) and fluid intelligence (Gf) show equivalent levels at age 10 and both peak around the age of 20, similar to the developmental trajectory of center frequency (CF). 2) Gf declines to the 10-year-old level by approximately 60, whereas CF recovers to childhood levels at around 90 years. This result suggests that the decline in Gf precedes the broader deterioration of brain function associated with aging. 3) In contrast, Gc remains relatively preserved, not reaching the 10-year-old level until well beyond 100 years of age. Indeed, at 95 years, Gc levels remain comparable to those observed in a 20-year-old. These findings indicate that Gc declines more slowly and manifests later than general brain aging, highlighting its resilience to age-related cognitive deterioration (Bugg et al., 2006; Bajpai et al., 2022).

**Figure 7:**
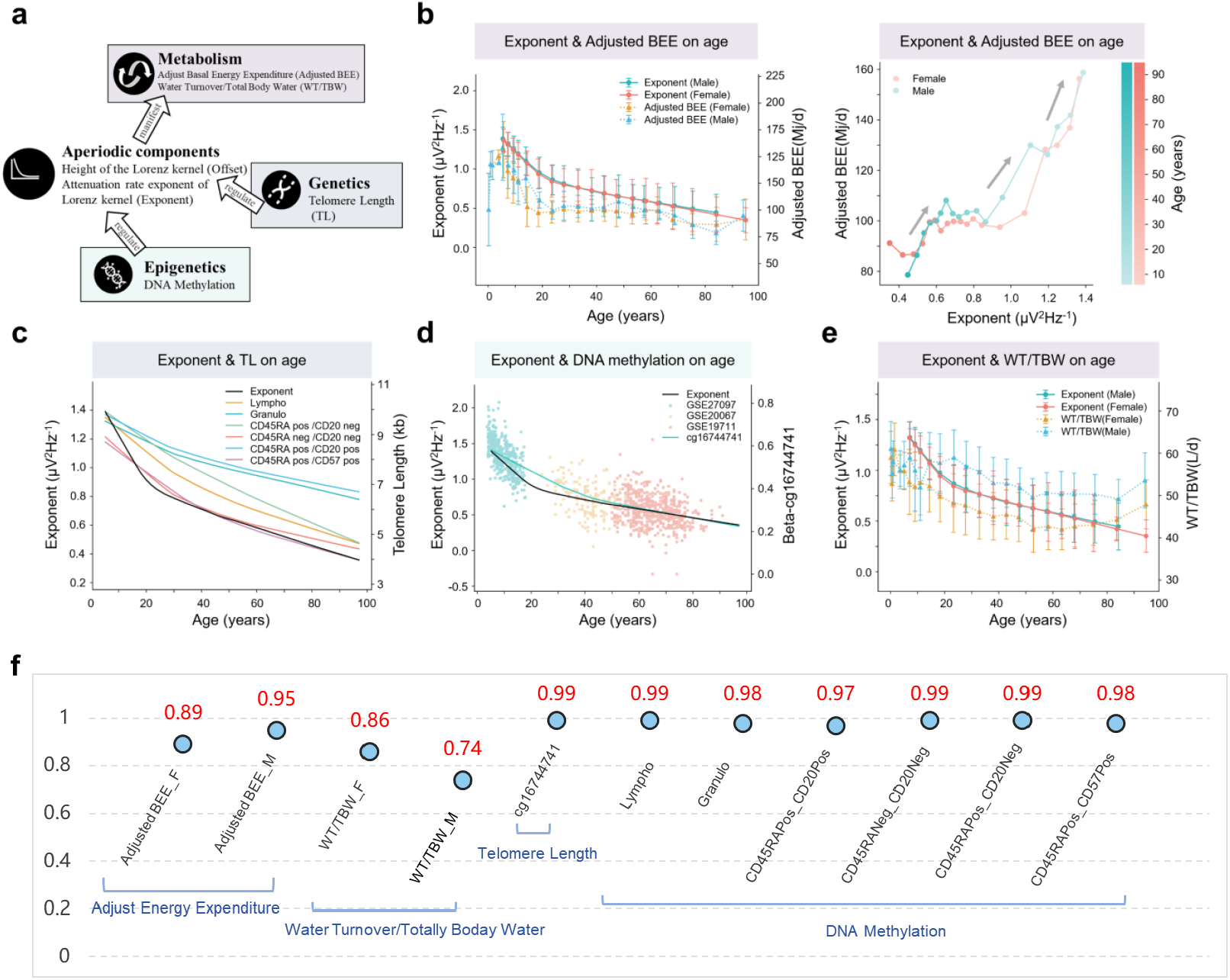
The relation of the developmental trajectory of Exponent with other modalities. a: Summary of all the modalities used. b: Exponent with Adjust BEE. The standard deviation is plotted here, and the values are directly from the paper Pontzer et al., 2021. The right panel displays the progression of Exponent with adjusted BEE over the life course, demonstrating how Exponent changes with adjusted age. c: Exponent with DNA methylation.; d: Exponent with Telomere Length. e: Exponent with WT/TBW. The standard deviation values are marked here, originally from the paper Yamada et al., 2022. f: The correlation values between Exponent and other modalities. The samples are resampled from the mean curve.

The trajectories of BEE and WT align with the developmental patterns of periodic EEG components. However, BEE scales with fat-free mass (FFM) according to a power-law relationship (Pontzer et al., 2021), necessitating adjustment for FFM to track its age-related dynamics. Castrillon et al. show that FFM-adjusted BEE declines sharply between ages 5 and 20, after which it stabilizes—mirroring the developmental trajectory of aperiodic EEG components Exponent and Offset (Figure 7b and Figure S16 b). This result suggests that, at rest, the energy consumed per unit of FFM primarily supports the brain’s intrinsic, non-oscillatory background activity. This observation aligns with findings showing that whole-brain glucose consumption drops to approximately half of normal levels in extreme neural inactivity, such as coma (Raichle and Gusnard, 2002). Similarly, the developmental trajectory of WT normalized by total body water (WT/TBW) parallels that of aperiodic EEG components (Figure 7e, Figure S16e), indicating that WT per unit of body mass reflects the strength of the brain’s background activity rather than its periodic oscillations.

In addition to analyzing data from higher-order regulatory modalities, we also examined primary epigenetic markers. Telomere length (TL) and DNA methylation are long-established biomarkers of aging. We compared their developmental trajectories with EEG components to uncover potential relationships. Figures 7c and S16c show that leukocyte telomere length (LTL) declines monotonically with age, consistent with the developmental trends of aperiodic EEG components. However, LTL shortens more slowly than EEG-derived measures between ages 5 and 20, with convergence in their rates of change observed after age 20. Moreover, TL trajectories vary across leukocyte subpopulations: CD45RA^−^/CD20^+^ and CD45RA^+^/CD57^+^ subsets exhibit steeper declines than others.

Beyond telomere biology, recent attention has focused on epigenetic clocks as robust biomarkers of biological age. Analysis of the Beta Prime value from whole-blood DNA methylation data revealed that methylation at cg16744741 follows a monotonic, nonlinear relationship with age (Alisch et al., 2012), closely mirroring the developmental trajectory of aperiodic EEG components (Figure 7d and S16d). Notably, age-related methylation changes vary across loci, with some sites showing positive correlations, others showing adverse, varying age-specific effects. Current large-scale DNA methylation charts predominantly model linear age associations in adult populations. However, studies such as Alisch et al. (2012) demonstrate that children and adolescents exhibit substantially higher rates of methylation change. These findings highlight the need for larger, more developmentally nuanced datasets and nonlinear modeling approaches to construct accurate, lifespan-spanning epigenetic clocks and enable meaningful cross-modal comparisons with neural activity metrics.

### 1.3.6 The Log-Linear Normative Model for Predicting Clinical Deviations

As shown in Figure 3, when plotted on a log10(age) scale, the developmental trajectories of both aperiodic and periodic components transform to log-linear norms. This simplification facilitates their use in brain age prediction and developmental modeling. Research indicates that the onset of psychiatric disorders occurs at specific age nodes (Kessler et al., 2005; Li et al., 2018) (Figure S18). Using these log-linear models as a normative reference framework, we sought to characterize the unique developmental signatures of deviation in various neurological and psychiatric disorders.

Data were sourced from the TDBrain dataset, publicly available under an open-access license with appropriate ethical approval. (van Dijk et al., 2022). The TDBrain dataset comprises individuals diagnosed with various neurological and psychiatric conditions, including Major Depressive Disorder (MDD), Attention Deficit Hyperactivity Disorder (ADHD), Subjective Memory Complaints (SMC), and Parkinson’s disease, along with age-matched healthy controls, a total of 711 subjects (6-78 years old) (Figure 8c). This broad clinical coverage enables investigation into the shared and distinct spectral biomarkers across diverse brain disorders.

**Figure 8:**
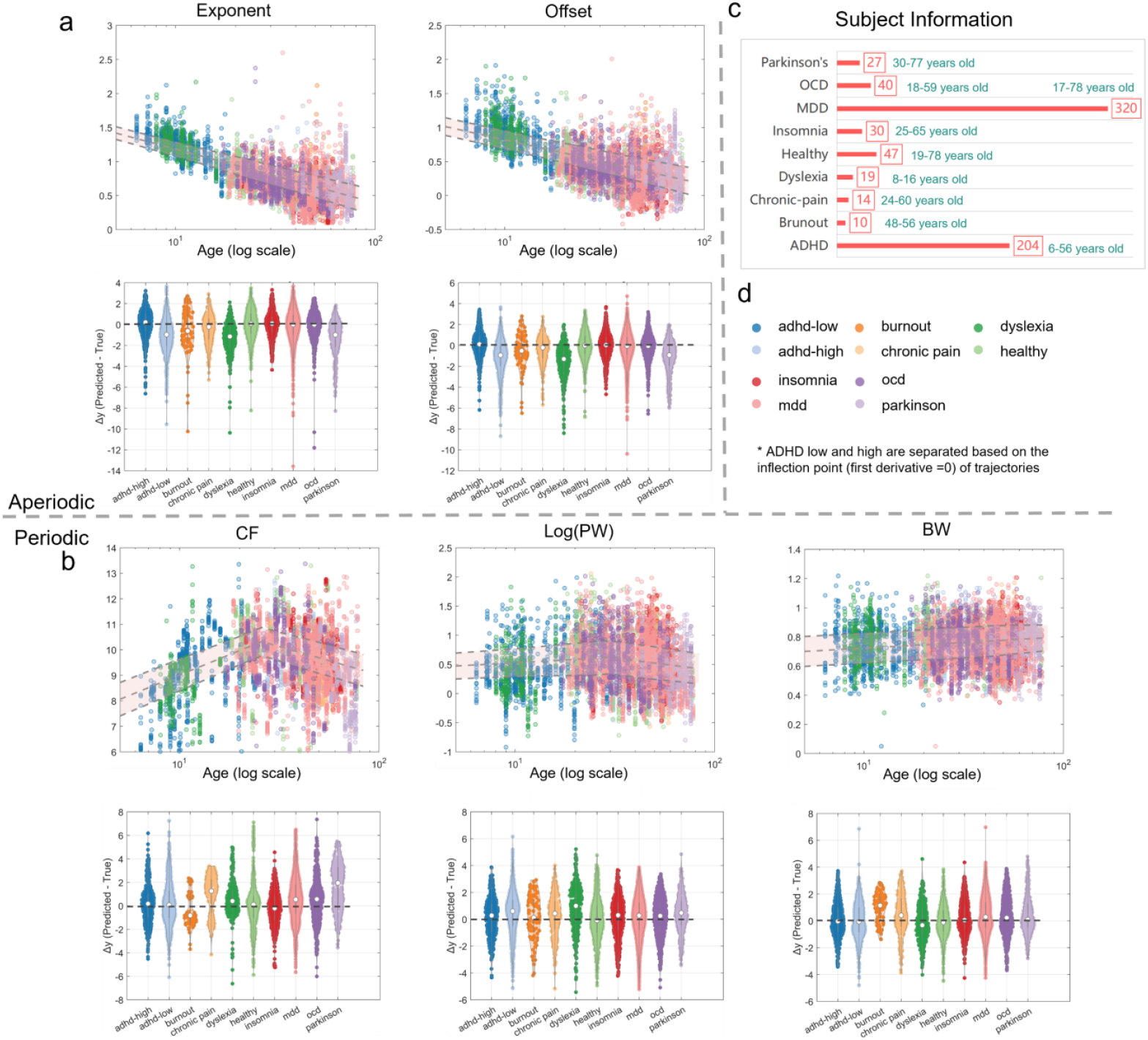
The prediction deviation distribution of the TDBrain dataset. a, b: Predicted deviations from age-adjusted norms for aperiodic (a) and periodic (b) parameters. Aperiodic parameters include the Exponent and Offset; periodic parameters include Central Frequency (CF), log-transformed Power (log(PW)), and Bandwidth (BW). Each panel displays individual data (scatter points) from 711 subjects (aged 6–78 years) overlaid on violin plots summarizing their distribution. The solid gray line indicates the normative mean derived from a multinational cohort, with dashed gray lines marking the confidence interval boundaries. The horizontal dashed black line represents zero deviation from the norm. c: Demographic summary of the subject cohort. d: Color key for experimental conditions.

The TDBrain dataset has undergone the batch effect adjustment with the HarMNqEEG protocol (https://github.com/LMNonlinear/HarMNqEEG). Given the broad age range of the ADHD cohort (6–56 years), we stratified these participants into ADHD-low and ADHD-high subgroups based on the key inflection point (first derivative = 0) of the developmental trajectory for each parameter. As shown in the Figure. 8, when analyzing deviations from the piecewise linear norms, the aperiodic components (Exponent and Offset) for the ADHD-low, burnout, dyslexia, and Parkinson’s disease groups exhibited positive deviations (i.e., observed values were higher than the norm). In contrast, other conditions showed negative deviations (Figure 8b). For the periodic parameters, nearly all conditions exhibited negative deviations, with the minor exceptions of a slight positive deviation in central frequency (CF) for burnout and bandwidth (BW) for dyslexia. This negative deviation was particularly pronounced in Parkinson’s disease (Figure 8a). These findings indicate that, compared to neurotypical controls, individuals with ADHD-low, burnout, dyslexia, and Parkinson’s disease have higher and steeper aperiodic activity. In parallel, in most of these conditions, the alpha peak occurred at a lower frequency with reduced power and narrower bandwidth, resulting in an overall flatter alpha spectral profile. Moreover, given the age-dependent heterogeneity of ADHD, aperiodic parameters serve as effective markers for distinguishing its age-related variations.

## 1.4 Discussion

Neural oscillations are shaped by a range of biological factors—including genetic expression (Frey and Morris, 1997; Flavell and Greenberg, 2008), hormonal regulation (Logothetis, 2003; Raichle, 2003; Mendoza-Halliday et al., 2024), and metabolic demands (Watson and Buzsáki, 2015)—throughout brain development, maturation, and aging (Cole, 2017; Cole and Voytek, 2017; Cole et al., 2018). In turn, these oscillations reflect underlying cognitive function (Fox et al., 2024) and the pathophysiological states of mental disorders (Mushtaq et al., 2024). Therefore, mapping the developmental trajectories of neural oscillations across the lifespan is essential for establishing normative models of brain aging, enabling personalized assessment of brain deviation, and improving brain age prediction.

Recent advances have established EEG-based developmental trajectories that offer direct insights into population-level patterns of neuronal cluster activity across ages (Li et al., 2022a). However, accumulating evidence indicates that EEG PSD consists of both periodic components— reflecting rhythmic neuronal synchronization—and aperiodic components, which capture background neural activity (Donoghue et al., 2020; Leroy et al., 2025; Wilkinson et al., 2024). These two components likely represent distinct neurophysiological mechanisms (Gil Avila et al., 2025; Mendoza-Halliday et al., 2024). Yet, whether each exhibits unique, age-specific developmental patterns remains unclear.

In this study, we harmonized multicenter EEG PSD data to derive, for the first time, comprehensive lifespan developmental curves (ages 5–95) for both periodic and aperiodic EEG components. We explore potential physiological and epigenetic mechanisms underlying their distinct developmental profiles by aligning these trajectories with those derived from other modalities. We found that the developmental trajectory of the periodic component correlates with advanced regulatory systems such as GABAergic neurotransmission, energy metabolism, white matter volume, and cognitive performance, following a Growth-then-Decline curve marked by initial increases followed by decline. In contrast, the aperiodic component shows a Monotonic Decrease with age, aligning with epigenetic markers such as DNA methylation levels and telomere length.

Through fine-grained analysis of age scaling and rate of change, we identified key developmental inflection points: around age 20, a critical transition phase occurs, while age 40 marks the onset of developmental stagnation. Notably, these ages correspond closely with the typical onset windows of major neuropsychiatric disorders across the lifespan (Kessler et al., 2005). While spectral parameters and energy levels exhibit significant age-related changes, their spatial distributions remain remarkably stable across development. This suggests that although global neural dynamics evolve with age, the topographical organization of oscillatory activity is preserved, reflecting a core invariant feature of brain architecture.

In summary, our findings establish a novel framework that dissociates the periodic and aperiodic components of EEG signals to understand brain development. This approach offers valuable insights into the underlying physiological and molecular mechanisms. It provides a crucial benchmark for quantifying developmental trajectories and assessing neurological abnormalities.

### 1.4.1 Physical Modeling of Periodic and Aperiodic Activities

Neurons are not independent entities; neural oscillations communicate through axonal transmission and the propagation of membrane charges, forming “traveling waves” in neural networks composed of neurons to transmit information (Cole and Voytek, 2017). Differences in neuronal firing times create specific spatial patterns, participating in complex and subtle cognitive processes such as attention, perception, and decision-making (Buzsáki, 2006). Based on this physiological foundation, since Hans Berger (Mushtaq et al., 2024) discovered brain waves, research on EEG and human consciousness and states has focused on exploring the periodic characteristics of specific regions and interactions between regions.

In addition to periodic components, the EEG spectrum contains many low-frequency components similar to 1/f noise (aperiodic components). Growing evidence suggests that these broadband aperiodic components can confound analyses of periodic activity (such as cross-frequency coupling) derived from EEG (Monchy et al., 2024). Simultaneously, aperiodic components weaken the correlation between EEG PSD and age; periodic components corrected for aperiodic influences exhibit steeper linear regression slopes (Tröndle et al., 2022). This does not, however, imply that aperiodic components are age-invariant. On the contrary, aperiodic components are essential features of neural data; their presence has been observed in various task scenarios with real data and in physical simulation models, and they depend on exposure factors and regulatory parameters (Brake et al., 2024; Kramer and Chu, 2023). Therefore, we need to separate the two components to explore their physiological and biological mechanisms more clearly.

The division of periodic and aperiodic components in the EEG power spectrum is a statistical and metaphysical method. A widely held view is that aperiodic activity is separable background noise. For example, Zetterberg et al. and Isaksson et al. demonstrated through first-order differences that an AR(1) process can generate 1/f activity, which is linearly additive with periodic activity (Isaksson et al., 1976a; Lopes da Silva et al., 1974; Zetterberg, 1969a). Pascual-Marqui et al. parameterized the EEG logarithmic power spectrum (EEG log-PSD) into xi and alpha processes (xi-alpha model) (Pascual-Marqui et al., 1995) through maximum likelihood estimation. Subsequently, least squares fitting of the logarithmic spectrum to parameterize and divide periodic and aperiodic components is popular now (Donoghue et al., 2020). Now, there have been several spectral parameterization methods, such as SPA (Spectral Parameter Analysis) (Isaksson et al., 1976b; Narasimhan, 1989; Zetterberg, 1969b), BOSC (Better Oscillation Detection), IRASA (Irregular-Resampling Auto-Spectral Analysis) (Hughes et al., 2012; Kosciessa et al., 2020; Seymour et al., 2022), SPRiNT (Spectral Parameterization Resolved in Time)(Wilson et al., 2022), and so on. The summary of spectral parameterization methods can be found in the paper (Wang et al., 2025 (Wang et al., 2025a)

However, the necessity and validity of spectral decomposition depend on the relationship between periodic and aperiodic components (Brake et al., 2024). Analysis of local field potential (LFP) power in the primate cortex reveals that oscillatory activity at specific frequencies is concentrated at different cortical depths, exhibiting a “spectral-laminar profile”; for instance, low-frequency aperiodic activity appears in superficial layers, whereas alpha-beta band electrical activity is found in deep layers (5/6). This segregation of oscillations implies the independence of periodic and aperiodic signals, and thus their additivity (Mendoza-Halliday et al., 2024). Similarly, experiments with biophysical models indicate that when a cortical neuron receives oscillatory inputs from the thalamus or subthreshold inputs from local cortical circuits, the amplitude of the spectral peak does not increase proportionally, implying an additive relationship between the two (Brake et al., 2024). In contrast, when biophysical parameters such as synaptic current properties undergo systematic changes, the spectral peak scales multiplicatively, a conclusion also verified using a multiplicative FOOOF algorithm (Brake et al., 2024). In an additive context, the model is formulated on a natural scale, while the multiplicative model is formulated as log*s*(*ω*) = log*ξ* (*ω*) + log*α*(*ω*). Therefore, Brake et al. (Brake et al., 2024) argue that different mechanisms require distinct spectral decomposition models to extract periodic and aperiodic components for further characterization of their respective features. As the relationship between periodic and aperiodic components is currently unclear—and given that most pharmacological experiments (e.g., with propofol, isoflurane, and ketamine) have found interdependent periodic and aperiodic activity via multiplicative models (Colombo et al., 2019; Gao et al., 2017a; Waschke et al., 2021), while additive models have been constructed from physical models of the resting state (Zetterberg, 1969a) and applied to the analysis of resting-state data (Hu et al., 2024; Pascual-marqui et al., 1988; Wang et al., 2025a)—we venture to hypothesize that periodic and aperiodic activities are additive in the resting state. Therefore, the model constructed and used here is an additive one.

### 1.4.2 Age Effects of Aperiodic Activity

Our previous work presented the EEG PSD variation with age and high-resolution frequency, showing that age differences are mainly reflected in the peak position in the alpha band and the amplitude of the EEG log-PSD (Li et al., 2022a), but lacked individual characterization of periodic and aperiodic components.

Currently, literature has reported the age characteristics of aperiodic components (Merkin et al., 2023b). Tro ndle et al. (Tröndle et al., 2021, Tröndle et al., 2020), based on EEG data from ages 5 to 20, found that the amplitude of the aperiodic component of the EEG PSD is negatively correlated with age. This is consistent with the conclusion in this study that in the alpha band from ages 5 to 20, PW (power) and CF (center frequency) are inversely related to offset and exponent with age. This also explains why periodic component parameters corrected for aperiodic components show a greater correlation with age.

In the aperiodic component, elderly individuals have flatter 1/f activity. Recent studies have also shown that compared to adolescents, the slope of aperiodic activity in the EEG of older people is lower. Complexity is reduced, which is explained by the neural noise hypothesis (Voytek et al., 2015). That is, the background noise of the brain continues to increase with age and work that continuously increases background noise in EEG signal simulation models and finds that the absolute value of the signal exponent decreases also supports the rationality of this hypothesis (Kramer and Chu, 2023; Lipsitz and Goldberger, 1992; Medel et al., 2023).

### 1.4.3 Physiological Mechanisms Explaining Pathological Changes in Periodic and Aperiodic Activities

Research generally believes that 1/f activity may involve changes in the excitation-inhibition (E: I) ratio, synaptic reductions, and decreased efficacy of GABAergic actions (Gao et al., 2017a). Experiments show that aperiodic activity can serve as an idealized indicator reflecting the E: I ratio, often used in detecting and explaining mechanisms of age-related neurodegenerative diseases and interpreting brain aging processes (Albertson et al., 2024; Bai et al., 2024; Gil Avila et al., 2025). However, there are doubts about using features like the slope of aperiodic activity to quantify and explain the physiological mechanisms of brain aging and lesions via the E: I ratio:

#### 1) Reflection of E: I Ratio Changes

Aperiodic activity in local field potentials (LFP) and scalp EEG reflects changes in the neuronal E: I ratio (Gao et al., 2017b). Experiments show that barbiturate anesthesia (Scott et al., 2014), propofol anesthesia (Solovey et al., 2015), or deep sleep (Meisel et al., 2017) mainly enhance GABA-A receptor activity, reduce the E: I ratio, leading to loss of consciousness. Their PSD exhibits a steeper 1/f decay than in the awake state. On the other hand, during xenon anesthesia (Hirota, 2006)and ketamine anesthesia, mechanisms mainly involve binding to NMDA receptors, inhibiting glutamate actions, thereby reducing neuronal excitability. The resulting high E: I may also lead to coma, similar to that during epileptic seizures (Meisel et al., 2012). Currently, the PSD exponent decreases less than low E: I ratios.

Therefore, it is generally believed that neurotransmitter release regulates aperiodic activity, and aperiodic components can serve as effective markers reflecting neurotransmitters like GABA (Zhou et al., 2021; Albertson et al., 2024). However, these studies overlook the relationship between periodic components and neurotransmitters, only emphasizing conclusions about aperiodic components. Periodic components are also regulated by neurotransmitters, and decreases in periodic components have been observed in young and aged groups compared to normal subjects in brain injuries like stroke(Albertson et al., 2024). Therefore, emphasizing the relationship between aperiodic components and GABA suggests that GABA directly regulates aperiodic components. However, this requires further comprehensive evaluation of the relative changes of periodic and aperiodic components.

#### 2) Differences in Brain Aging Processes

During brain aging, the slope of aperiodic activity gradually decreases, and the intercept decreases. The background noise activity of the brain gradually expands to wide frequencies, and periodic coupling activities gradually decrease with age (Podvalny et al., 2015; Voytek et al., 2015). Although this seems similar to the process where neuronal activities in the brain during anesthesia (Colombo et al., 2019; Waschke et al., 2021) and near-death states (Zhou et al., 2021) gradually collapse toward scale-free activities— i.e., periodic activity coupling decreases—in the anesthesia state, the slope of aperiodic activity is steeper compared to the awake state (changes in intercept and total energy are not reported). These may be two different processes.

The total brain energy does not change quickly in anesthesia and near-death states. Still, it rapidly transitions from periodic to aperiodic activity (Zhou et al., 2021), showing decreased periodic and increased aperiodic activity. It then gradually returns to normal, or activities cease. However, natural aging is a long, gradual process. From the results of this study, during brain aging, the total energy of both aperiodic and periodic activities decreases. Although the energy of high-frequency bands like beta increases with age in the periodic component, overall, the decrease in energy with age is greater than the increase. Therefore, in natural aging, there is a phenomenon of reduced periodic coupling activity and flatter aperiodic components. Based on this, explaining brain aging according to neural mechanisms like anesthesia requires further consideration.

#### 3) Correlation with Neurotr(Franks, 2008; Waschke et al., 2021)ansmitters

A large body of research, including animal (Gao et al., 2017a)and human experiments(Donoghue et al., 2020; Mamiya et al., 2021; Ostlund et al., 2022) as well as computational modeling, has investigated the relationship between aperiodic activity and the Excitatory-Inhibitory (E:I) balance. These findings indicate that a flattening of the aperiodic spectrum is associated with an increased E:I ratio. For instance, reduced overall inhibition—caused by blocking excitatory N-methyl-D-aspartate (NMDA) receptors and the associated decrease in GABA release—results in a flattened broadband spectrum (Deane et al., 2020; Waschke et al., 2021). Conversely, a decreased E:I ratio leads to a steeper broadband spectrum, as exemplified by propofol, which enhances inhibition by increasing GABAergic activity (Franks, 2008; Waschke et al., 2021).Concurrently, these anesthetic agents also exert broad effects on the overall level of neural activity (Taub et al., 2013) and on oscillatory activity within the periodic alpha, beta, and gamma bands. The results of the present study show that GABA levels are highly correlated with the developmental trajectories of periodic (but not aperiodic) activity. However, because our study did not include data on excitatory neurotransmitters, the specific contributions of the E:I balance to periodic and aperiodic

### 1.4.4 Multimodal Relationships

Our study shows that, from the developmental trend within the lifespan, aperiodic activity is more consistent with the trajectories of telomere length and DNA methylation levels. Telomere length and DNA methylation are essential markers for predicting biological age, influenced by genetic regulation, inflammation, immunity, etc. They not only reflect an individual’s degree of aging but may also be involved in the occurrence of various age-related diseases.

Telomere length strongly correlates with lifespan; when cells stop dividing and enter a senescent state, they undergo programmed apoptosis. Studies have found that Parkinson’s patients exhibit longer telomere lengths (Fani et al., 2020; Forero et al., 2016), and excessively long or short telomere lengths are associated with cognitive decline. In the brains of Alzheimer’s patients, specific genes related to neuronal plasticity and survival (such as the BDNF gene) also show changes in methylation levels.

These conclusions indicate that the brain’s aperiodic component may, like telomere length and DNA methylation, be an objective clock of the human cell cycle, apoptosis, and proliferation. It is directly regulated by genes and influenced by various external factors like cell damage and inflammation, thereby affecting neuronal plasticity and function, accelerating brain aging.

Ages 20 and 40 mark pivotal transitions in human brain development, reflecting distinct neurobiological, cognitive, and physiological shifts. Around age 20, the brain undergoes a critical transition from adolescence to adulthood, characterized by profound physiological, psychological, and social transformations. This period aligns with the near-complete maturation of the prefrontal cortex, which underpins executive functions such as decision-making and impulse control, as demonstrated by Arain et al. (Sharma et al., 2013). White matter myelination, essential for efficient neural conduction, peaks in the late 20s, as shown by Bethlehem et al. (R. A. I. Bethlehem et al., 2022), representing a neurobiological apex that supports enhanced cognitive control and social role transitions, such as workforce entry or higher education (Johnson et al., 2009). Sex hormones, including estrogen, progesterone, and testosterone, drive myelination, and synaptic pruning, sculpting neural circuits by age 20 (Sharma et al., 2013). Efficient neurotransmitter release, particularly dopamine and acetylcholine, bolsters synaptic communication, while genetic influences, including GABAergic and cholinergic systems, shape oscillatory patterns critical for cognitive processing (Begleiter and Porjesz, 2006).

Brain development peaks around age 20, enters a negative acceleration state, and begins to age at age 40, corroborated by changes in neurotransmitter release and cognitive abilities. By age 40, brain development transitions into accelerated aging, marked by non-linear declines in white matter integrity and cortical thickness, as reported by Dohm et al. (Dohm-Hansen et al., 2024). White matter volume begins to decline around this age, coinciding with early reductions in cognitive functions such as episodic memory and processing speed (R. A. I. Bethlehem et al., 2022). Myelin deterioration and increased white matter lesions, particularly in the frontal lobes, further underscore this turning point (Peters, 2006). At the molecular level, gene expression shifts toward immune-related pathways and away from synaptic maintenance, reflecting a critical transition in brain aging (Dohm-Hansen et al., 2024). Hormonal changes, notably declining estrogen levels in women during perimenopause, alter neural activity, as evidenced by menopause-related changes in oscillatory patterns (Tegeler et al., 2015). Metabolic dysregulation, such as impaired glucose metabolism, also impacts brain function, particularly in aging or diabetic populations, contributing to altered neural dynamics (Yeung et al., 2025). Cognitive declines in processing speed and memory at age 40 are linked to reduced neurotransmitter efficiency and diminished synaptic plasticity (Dohm-Hansen et al., 2024; Peters, 2006).

### 1.4.5 Limitation

Due to data limitations, the study did not include children under five. Previous studies have reported the developmental trends of periodic and aperiodic brain activity from 2 to 44 months of age (Rico-Picó et al., 2023; Wilkinson et al., 2024). Aperiodic parameters show an overall increasing trend but with greater fluctuations. This provides a basis for the future inclusion of pediatric data to conduct cross-age studies and characterize developmental patterns.

Due to inconsistencies in electrode montages across multicenter EEG data, this study standardized all the other montages to the international 19-channel (18 channels after average reference) 10-20 system montage (Seeck et al., 2017) with EEG inverse method (Dong et al., 2021). Although 19-channel EEG offers lower spatial resolution than high-density systems, it was fully adequate for the scope of our investigation, which focused on elucidating the overarching developmental trajectories and spatial trends of aperiodic and periodic components, not on resolving subtle inter-group differences within specific cortical parcels. However, future high-density EEG studies will be essential to refine these norms for clinical applications, particularly for assessing disease progression and subtypes. The high-density EEG is also critical for source-space localization, enabling more profound insights into these neural components’ electrophysiological mechanisms and functional roles.

The interplay between periodic and aperiodic components and their underlying generative mechanisms remains under intensive investigation. In studies employing physiological and biophysical modeling approaches, age represents a critical variable, underscoring the necessity of longitudinal investigation. Furthermore, the present study does not address the developmental trajectories of functional connectivity derived from periodic and aperiodic components—a critical line of inquiry for elucidating pathophysiological states and functional deficits that emerge across development, maturation, and senescence (Friston, 2011; Valdes-Sosa et al., 2011; B. He et al., 2019).

### 1.4.6 Future Directions

In the context of multimodal interactions, our results suggest these neural signals track distinct biological processes. For instance, the aperiodic component of EEG exhibits a developmental trajectory that parallels changes in telomere length and DNA methylation. In contrast, periodic components align with processes such as GABAergic neurotransmission and metabolic energy consumption. Concurrently, periodic activity mirrors the maturation of human cognitive faculties. These convergent findings inspire the pursuit of a unified “biological clock” for human aging and open new avenues for drug development and identifying clinical diagnostic indicators. However, the scarcity of suitable multimodal datasets precluded an analysis of age-related effects on the interplay between modalities during cognitive tasks or in the context of neurological dysfunction. With growing support from global open-data initiatives and policy development, it will become possible to leverage large-scale multimodal data to construct a comprehensive causal knowledge graph of neurodevelopment. Such a resource would enable a multi-level, multi-perspective analysis of brain development, maturation, and senescence processes. Besides the above, there are still the following questions worth considering:

#### 1. Age-Dependent Frequency Band Segmentation

Existing frequency band classifications do not account for age differences. Frequency band classification rules should be age-dependent and based on two key developmental periods combined with brain cognitive development.

#### 2. Impact of Stress and Inflammation on Telomere Length

Telomere length is influenced by stress and inflammation. Do these effects manifest in the aperiodic components of EEG?

#### 3. Heritability of EEG Components

Telomere length exhibits an intergenerational inheritance mechanism. Do the periodic and aperiodic components of EEG possess hereditary characteristics?

#### 4. Causal Relationship Between Periodic and Aperiodic Components

The causal relationship between periodic and aperiodic components remains unclear. Is their interaction disease-specific?

#### 5. Environmental and Cognitive Training Influences on Brain Rhythms

Do environmental factors and cognitive training influence brain rhythm characteristics, and do these changes similarly occur in other modalities?

#### 6. Reversible Developmental Processes in the Brain

Does the brain undergo reversible developmental processes? More extensive datasets and research on age-related brain developmental abnormalities are needed.

#### 7. Detailed Analysis of Female Brain Development During Pregnancy

Female brain development during pregnancy requires more detailed analysis.

#### 8. EEG Characteristics in Infant Brain Development and Their Relationship with Other Modalities

Explore EEG characteristics in infant brain development and their relationship with other modalities.

#### 9. Enhancing Multimodal Cross-Prediction and Evaluation Using Lifecycle-Spanning and Nonlinear Modeling

Improve multimodal cross-prediction and evaluation through lifecycle-spanning and nonlinear modeling. For example, m**odality Substitution Indicators**, are there modality substitution indicators, such as inferring telomere length from aperiodic EEG characteristics?

#### 10. Predictive Ability of Brain Age for Neurological Diseases

Mental Illnesses, and Mortality Risk: Given that different diseases are characterized by critical periods of onset, it is essential to assess the predictive capability of brain age for neurological diseases, mental illnesses, and mortality risk.

Our study provides the first characterization of the developmental trajectories and spatial topographies of both periodic and aperiodic components of neural signals. In doing so, we highlight two distinct modes of neurodevelopmental change, identify critical transitional periods, and establish robust linear and piecewise-linear normative models. These models provide crucial benchmarks for assessing brain developmental status and disease prediction. Furthermore, our correlation analyses with developmental trajectories from other modalities offer a novel framework and empirical basis for the integrated investigation of human brain development, maturation, and aging.

## 1.5 Method

### 1.5.1 Data acquisition

The data used in this study were obtained from the HarMNqEEG Project (https://doi.org/10.7303/syn26712693), consisting of eyes-closed, resting-state recordings from healthy participants. The data were collected from 14 studies across 9 countries, involving 12 different EEG recording devices, specifically intended for the construction of quantitative EEG (qEEG) norms. To enhance privacy protection and facilitate seamless data sharing, this project adopted a decentralized data compilation model through the GBC project (https://globalbrainconsortium.org/about/). This approach involved distributing standardized computational scripts to each collaborating site. In return, the sites submitted only the computed cross-spectra and associated metadata (participant age, sex, country, and acquisition equipment), eliminating the need to transfer raw data. To further safeguard privacy, all submitted data were anonymized using Arng2 (Figure 9a). Upon collecting, the data adhered to the following criteria:

**Figure 9:**
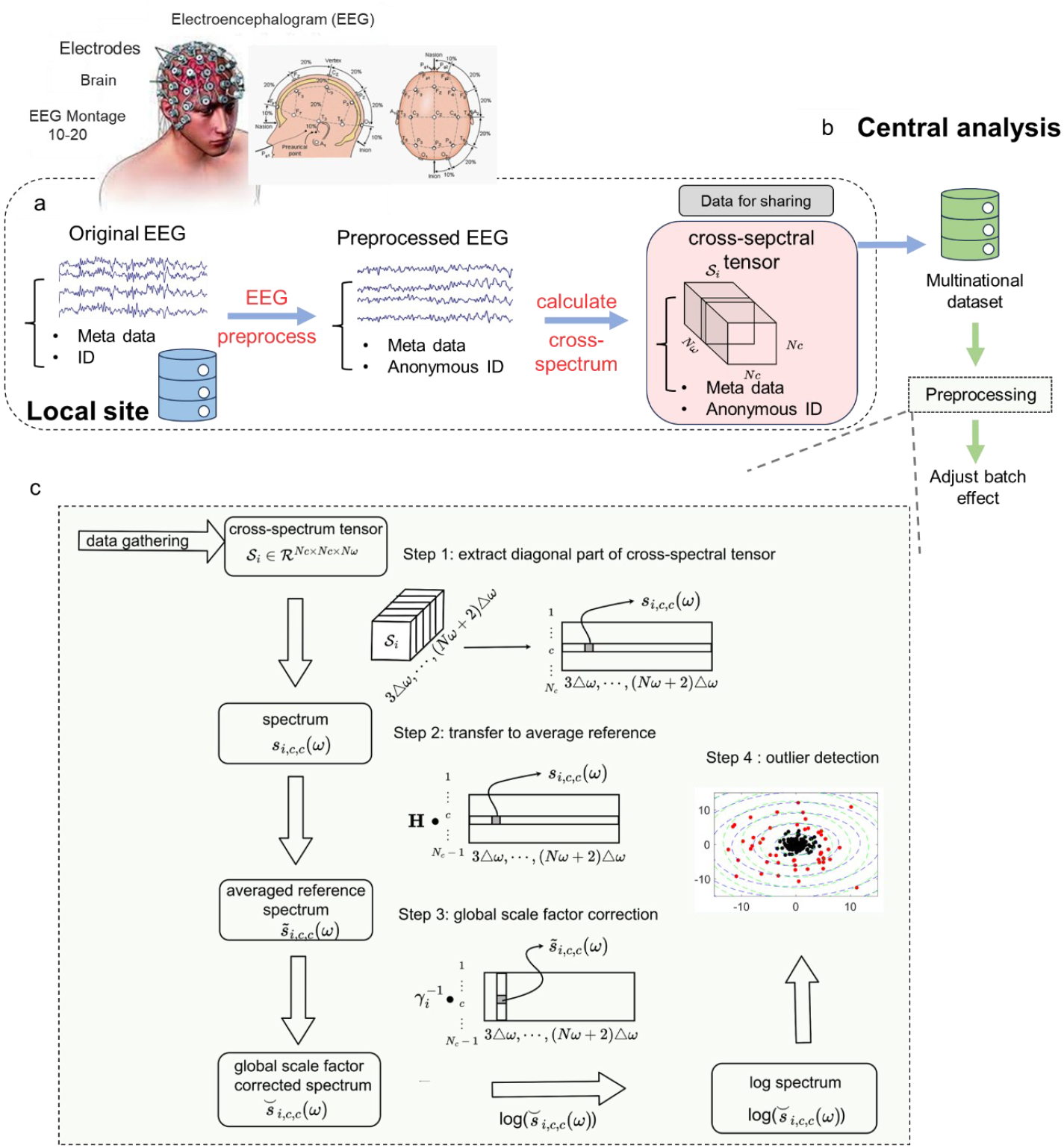
The multinational dataset collection and preprocessing pipeline

1. Standardization for EEG norms: All data were sourced from standard datasets used to establish EEG norms, consisting of at least one minute of eyes-closed, resting-state EEG recordings.
2. Preprocessing procedures: Preprocessing at all sites did not include ICA processes. Each batch has to pass the multinational data quality control protocol (at the following outlier detection section), and the site-specific preprocessing variations and physiological artifacts (e.g., EOG, ECG, and EMG) were removed as batch effects.
3. To ensure uniform data processing, the EEG data were interpolated to the 19 channels of the International 10-20 system, based on the EEG inverse method (Dong et al., 2021) before the spectrum calculation in the shared *data_gather* script.
4. Epoch segmentation: Each local site run the uniform shared *data_gather* script, which segmented the data into artifact-free 2.56-second segments to ensure a uniform frequency resolution *Δω* = 0.3906. The length of each segment is ⌈ (1/0.3906) × *Sampling rate* ⌉. Because of the original data lacking of Cuba-1990 dataset which only preserve EEG-PSD up to 20Hz, thus, to align all the dataset, here the EEG-PSD are up to 20Hz.

Following the completion of data collection, the central site will be responsible for conducting multinational data preprocessing and batch effect harmonization (Figure 9b).

### 1.5.2 Spectral calculation

Bartlett’s (BT) method (Møller, 1986) was employed in the *data_gather* to calculate the EEG cross-spectra. The BT method is a direct power spectral estimation technique that involves dividing the signal into segments, computing the periodogram for each segment (by taking the square of the direct Fourier transform), and then averaging these results across more than 20 non-overlapping segments to reduce variance. The window for BT used here is the classical rectangular window with length ⌈ (1/0.3906) × *Sampling rate* ⌉ for each data.

Given two discrete-time signals *x*[*n*]and *y*[*n*]of length *N*, divide it into *K* non-overlapping segments of equal length *L* = *N/K*. The -th segment is defined as:

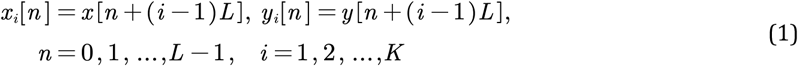

For each segment, the Fourier transformer is,

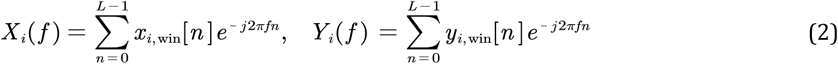

and the *i*-th cross-periodogram is,

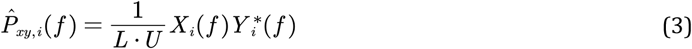

Average the cross-periodogram along all segments, the cross-spectrum is calculated as,

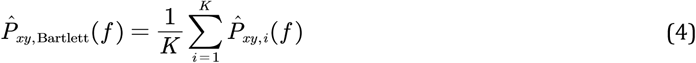

Specifically, each segment of EEG data was automatically divided into an interference-free time series of 2.56 seconds (with EEG data segment length determined as ⌈ (1/2.56) × *Sampling rate* ⌉^1^), and the resulting cross-spectra achieved a frequency resolution of Δ*ω* = 0.39 Hz. Each participant’s cross-spectrum is a 3-D tensor, and together with the participant’s metadata, they were packaged into a structured format and shared using the BIDS (Brain Imaging Data Structure) data format (Pernet et al., 2019).

### 1.5.3 Data preprocessing

In readiness for data sharing, each collaborating site conducted the preprocessing of EEG data before sharing. To maintain consistency in the number of electrodes and reference settings across multicenter data, the central site performed unified preprocessing on the data after the distributed collection of cross-spectrum structures. This preprocessing included re-referencing, batch effect removal, and other steps. The specific procedures are as follows (Figure 9c)

1-Re-referencing: All EEG cross-spectra were re-referenced to the average reference.

The average reference method assumes that the sum of potentials from all electrodes is zero (i.e., the assumption that the head is an electrically neutral, closed system), which in theory makes it applicable to any electrode montage.

The collected cross-spectrum matrices **S**_*i*_ (*ω*) from the original recording montages to the average reference with the following transformation (Bertrand et al., 1985; Hu et al., 2019)

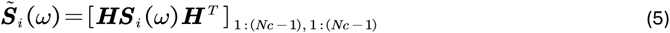

here, the center operator is 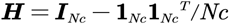. The final electrode (Pz) was eliminated to avoid the trivial singularity of matrices and the number of electrodes *Nc* changes from 19 to 18.

The sparse spatial sampling of a 19-channel EEG montage, which primarily covers superior scalp regions, offers insufficient coverage of inferior areas like the temporal base and inferior occipital lobe. Consequently, an average reference derived from such a montage can be biased.

To mitigate this issue, our study utilized data acquired with the standard international 10-20 system, which minimizes errors arising from inconsistent electrode placement. Furthermore, site-specific preprocessing had already been performed on the data to correct for signal drift and bad channels, thereby substantially reducing potential sources of reference error (Bertrand et al., 1985).

2-Global Scale Factorization (GSF): A fundamental challenge in EEG analysis is the marked inter-individual variability in global signal amplitude, wherein signals may exhibit qualitatively similar waveform morphologies yet differ substantially in overall power. To formally quantify and correct for this discrepancy, J. L. Herna ndez et al (Herna ndez et al., 1994). introduced the concept of a global scaling factor. Application of this factor rescales the EEG data, thereby removing this source of spurious variance from the analysis.?

Suppose for an individual *i, v*_*i,c*_ (*t*) = *γ*_*i*_*p*_*i,c*_ (*t*) is the EEG potential recorded at the electrode *c,γ*_*i*_ is the global scale factor which is dependent with electrode a and time and *p*_*i,c*_(*T*) is the scale free EEG potential. With the logarithm transformation, the EEG records *l*_*i,c*_(*ω*) = *κ*_*i*_ + *σ*_*i,c*_ (*ω*) where *l*_*i,c*_(*ω*) is the log transform of *v*_*i,c*_ (*t*), *κ*_*i*_ = 2*log*(*γ*_*i*_) and *σ*_*i,c*_ is the log-spectrum of EEG records. Write the log form into the vector format, 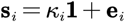,here, **s**_*i*_ and **e**_*i*_ are vectors stacked with all log spectrum and **1** is a vector which norm is 1. The log transform global scale factor *κ*_*i*_ is colinear with vector **1** and *κ*_*i*_, and is the geometric mean of all 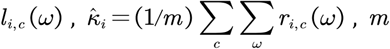, *m* is the sum of the number of frequency points and electrodes. The EEG recording could be rescaled by dividing 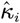.

3-Log Vectorization: The diagonal elements of the cross-spectra were extracted and logarithmically transformed to perform Gaussian vectorization for log-spectra.

The details of the preprocessing can be found in the paper (Li et al., 2022b).

### 1.5.4 Data quality control

Outliers are challenging in EEG research, especially when constructing normative models. The presence of outliers can skew the resulting charts, leading to biased estimates. Outliers violate fundamental assumptions and can disrupt model estimation, comparisons, and inferences. This is a common and unavoidable problem, typically caused by recording artifacts, electrode misplacement, data processing errors, and other factors, and can arise at any stage of the normative study workflow.

In this study, batch effects and outlier biases are intricately intertwined, making conventional methods based on observed variables ineffective. We employed global z-scores for outlier detection instead of traditional observer-based practices to tackle this issue. The global z-scores are derived from a nonlinear normative model that removes the age effect and should adhere to a normal Gaussian distribution for healthy subjects, 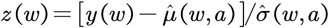. With the fact that the maximum likelihood covariance estimator (MLE) is sensitive to outliers (Leroy and Rousseeuw, 1987) and this releases the relationship between the Mahalanobis distance and the robust distance in a normal Gaussian distribution is represented by the line y=x, which indicates that outliers will be positioned away from this line in a DD (Mahalanobis distance vs robust distance) plot (Olive, 2004). Consequently, we can automatically detect and remove outliers by applying specific criteria 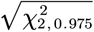, the value of inverse chi-square cumulative distribution with two degrees of freedom for probability equal to 0.975 with the global z-score.

### 1.5.5 Batch harmonization process

Multicenter studies provide a wealth of data resources for constructing normative models, particularly when broad age ranges are required and a globally inclusive norm is desired (R. a. I. Bethlehem et al., 2022). This approach aligns with the current trends and directions in global open science research (Valdes-Sosa et al., 2022). However, a significant challenge in multicenter data research is the presence of batch effects. In neuroimaging studies, batch effects refer to confounding factors introduced by non-physiological variables, typically including differences in acquisition equipment, preprocessing preferences, ethnic variations, and other factors (Johnson et al., 2007). A substantial body of literature has documented significant batch effects in multicenter studies involving EEG (Li et al., 2022b) and MRI (Hu et al., 2023), which severely impact the reproducibility and reliability of results (Gouttard et al., 2008).

The demand for large sample sizes for EEG normative models introduces variability from different vendor recording systems, lack of standardized amplifier parameters, electrode placement systems, and preprocessing protocols exacerbates the issue of batch effects. This problem is particularly pronounced and can significantly compromise the accuracy and applicability of the constructed normative models.

In this study, building on our laboratory’s previous work in developing multicenter multivariate normative models (Li et al., 2022b), we performed batch effect harmonization on the multicenter log-spectrum. This harmonization is crucial for ensuring the accuracy and reliability of the normative models across diverse data sources, thereby enhancing their applicability and robustness in global research contexts.

Assuming the log-spectrum for the subject *i* of the electrode *c* is *y*_*i,c*_(*ω*), for any frequency point *ω* and age *a*, the developmental surface could be written as,

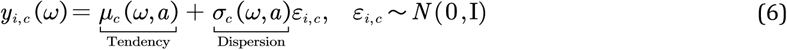

Here, the developmental tendency 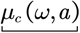, also known as the EEG log-spectrum norm at each electrode. Standard deviation *σ*_*c*_(*ω,a*) describes the dispersion of the developmental trajectory, which is sampled from the Gaussian distribution. It should be emphasized here that the trend and dispersion are nonlinear functions of both age and frequency, rather than functions of only age or frequency. The nonlinear function factor selection has been verified in the paper (Li et al., 2022b) through comparison with the model’s general cross-validation values (GCVs).

For multicenter data analysis, considering the gender and batch effect as random effect, equation (5) could be written as follows,

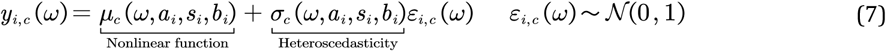

As shown in (Li et al., 2022b), with the hierarchical strategy, the optimal model for the log-spectrum is

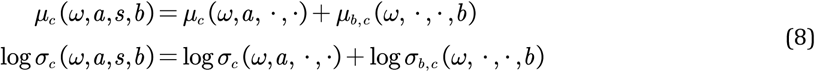

The equation (7) means that there is no sex effect, but the batch effect, and for the log-spectrum, the batch effect is related to frequency instead of age. Following harmonization, the results were assessed both graphically and statistically, with details of the batch validation provided in Supplementary Information 2.2.

### 1.5.2 Robust EEG spectral parametrization

Aperiodic and periodic signals, arising from distinct cortical regions (Mendoza-Halliday et al., 2022), are treated as independently generated processes (Kramer and Chu, 2023; Brake et al., 2024). This premise aligns with Brillinger’s spectral and Volterra theory, where a signal’s moments of all orders adhere to the principle of additivity for independent oscillating sources (Brillinger, 2001; Billings, 2013). Thus, as presented in the paper (Pascual-marqui et al., 1988; Valdés et al., 1992; Wang et al., 2025b), the single-channel EEG power spectrum could be divided into two type of dependent components in nature scale,

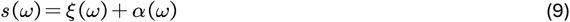

One type is process like 1/f activity *ξ*(*ω*), we called it aperiodic component, which has been discussed extensively from the view of statistical (He, 2014), biophysical (Gao et al., 2017a; Lindén et al., 2010), and neural models (Kramer and Chu, 2023). Another type is composed with a set of harmonic activities *α*(*ω*), we call them periodic components. The *ξ*(*ω*) activity could be parameterized with the Lorentzian model,

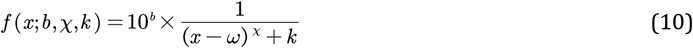

where *b* is the height or offset of the board band kernel, which is the gain factor in control system, and with format of power of 10, typically represents a gain adjustment on a decibel (dB) scale. A higher value of *b* means an increased amplification across all frequency ranges. *χ* is the attenuation rate exponent, and it determines how quickly the system response decays as the frequency deviates from the center point. A larger *χ* results in a sharper roll-off, similar to a high-Q filter, while a smaller *χ* leads to a more gradual transition, like that of a low-Q filter of the kernel. *ω* is the center position of kernel. is the baseline offset, a small offset term to prevent the denominator from being zero, and it can also be understood as a background value or saturation term. It should be noted that the *k* affects the height of the bottom part of the function which describes the ‘knee’ of the function (Donoghue et al., 2020).

The harmonic activity could be expressed with Gaussian function,

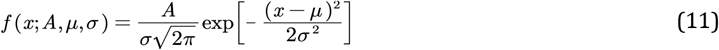

here, *μ* represents the peak position of the function, and *σ* is the scale parameter, it is also known as half of the Full Width at Half Maximum (FWHM), determines the width of the distribution, i.e., how “sharp” the peak is. A larger *σ* value indicates a wider distribution, while a smaller *σ* value suggests a sharper and more concentrated peak. *A* is the amplitude factor, controls the height of the kernel. The corresponding parameters used in paper show in table 1.

**Table 1:**
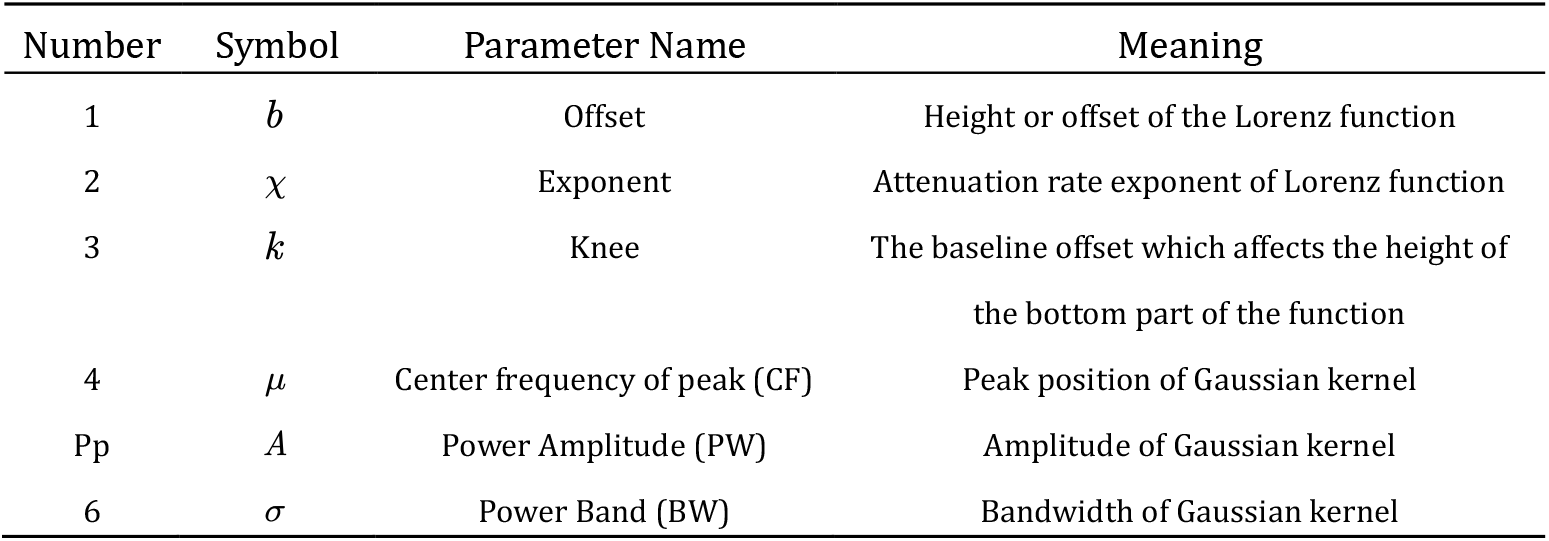
Aperiodic and periodic parameters symbol.

| Number | Symbol | Parameter Name | Meaning |
| --- | --- | --- | --- |
| 1 | $b$ | Offset | Height or offset of the Lorenz function |
| 2 | $\chi$ | Exponent | Attenuation rate exponent of Lorenz function |
| 3 | $k$ | Knee | The baseline offset which affects the height of the bottom part of the function |
| 4 | $\mu$ | Center frequency of peak (CF) | Peak position of Gaussian kernel |
| Pp | $A$ | Power Amplitude (PW) | Amplitude of Gaussian kernel |
| 6 | $\sigma$ | Power Band (BW) | Bandwidth of Gaussian kernel |

It should be noted that Equation (9) is additive on a nature scale, not the logarithmic scale used in FOOOF. The issue of additivity versus multiplicativity will be discussed in detail in the Discussion section.

The solution to Equation (9) is formulated as an optimization problem and can be effectively addressed using non-linear Least-Squares Minimization (NLSM). There are two main strategies for fitting separate kernels:

1. **Divide and Conquer**, employed by FOOOF, iteratively estimates the aperiodic and periodic components in an alternating fashion. This approach allows for faster convergence, but it comes with a trade-off: the absolute magnitudes of the periodic and aperiodic components mutually influence each other, especially in the presence of a ‘knee’. Consequently, the initial step of removing the aperiodic component to fit a flattened Power Spectral Density (PSD) may not yield a globally optimal solution. This is precisely why the FOOOF algorithm first needs to determine whether a knee should be included in the model. Although this can be assessed beforehand by plotting the power spectrum on a log-log scale and observing if the overall trend is linear, this method undoubtedly adds to the workload.
2. **Simultaneous Optimization**, as implemented in the lmfit toolkit (https://lmfit.github.io/lmfit-py/), fits all component sources concurrently. This method generally provides higher accuracy by considering the interactions between components during the optimization process (check the results in the model comparison section).

The Levenberg-Marquardt (L-M) algorithm is an iterative optimization technique widely used for solving non-linear least squares problems. It effectively combines the strengths of both the Gauss-Newton method and the Gradient Descent method. When the current parameter estimates are close to the optimal solution, the L-M algorithm achieves fast convergence by leveraging the inner product of the Jacobian matrix. Conversely, in situations where convergence is slow or oscillatory, it enhances robustness through the introduction of a damping factor.

The solution to Equation (8) can be obtained using the L-M algorithm; however, a critical challenge lies in the appropriate selection of initial parameter values of kernel fitting. In the context of EEG spectra, the parameterization of Gaussian and Lorentzian kernels is inherently interdependent, making the choice of initial values crucial for accurately determining kernel localization and related characteristics.

Given that EEG spectra typically contain a substantial amount of low-frequency power, selecting initial parameters for the Lorentzian kernel is relatively straightforward, often centered at *ω* = 0. In contrast, initializing the parameters of the Gaussian kernel—which models periodic activity—is significantly more complex. The behavior of the Gaussian kernel is primarily governed by its bandwidth: if the bandwidth is too large, multiple spectral peaks may be captured by a single kernel, leading to underfitting; conversely, if the bandwidth is too small, it may result in overfitting due to excessive kernel proliferation.

However, initializing the bandwidth (BW) is a challenging task. In contrast, the location (CF) and height (pw) of peaks can adaptively control extreme bandwidth values during the optimization process, and these parameters (CF and PW) can be effectively initialized through certain strategies. Therefore, initializing the position and amplitude of the Gaussian kernel is a key step in the parameterization process.

Currently, the primary method for initializing the peak locations relies on the “findpeaks” function, which selects peaks based on the number of peaks or their minimum hight. Like FOOOF, which controls the number of peaks using two criteria: a relative threshold calculated from the standard deviation (SD) of the spectrum (defaulting to thresh_relative_ = 2\**SD*), and an absolute threshold for peak height (default to thresh_absolute_ = 0). However, this approach involves numerous parameters that are difficult to tune effectively. Improper selection may lead to the identification of peaks with high amplitude but very wide widths. Such peaks do not align with the characteristics of EEG signals and may be caused by noise or oscillations introduced by trends. The FOOOF method initializes Gaussian kernel peaks by removing the fitted aperiodic component ξ from the original signal. However, this approach is heavily dependent on the quality of the ξ component fitting. If the signal’s trend is not adequately removed, it can lead to false peaks, resulting in errors in the location and height of the identified peaks.

Here, we propose a dual residual initialization strategy for Gaussian kernel initialization and apply the simultaneous optimization method. The procedures are as follows (Figure 10):

**Figure 10:**
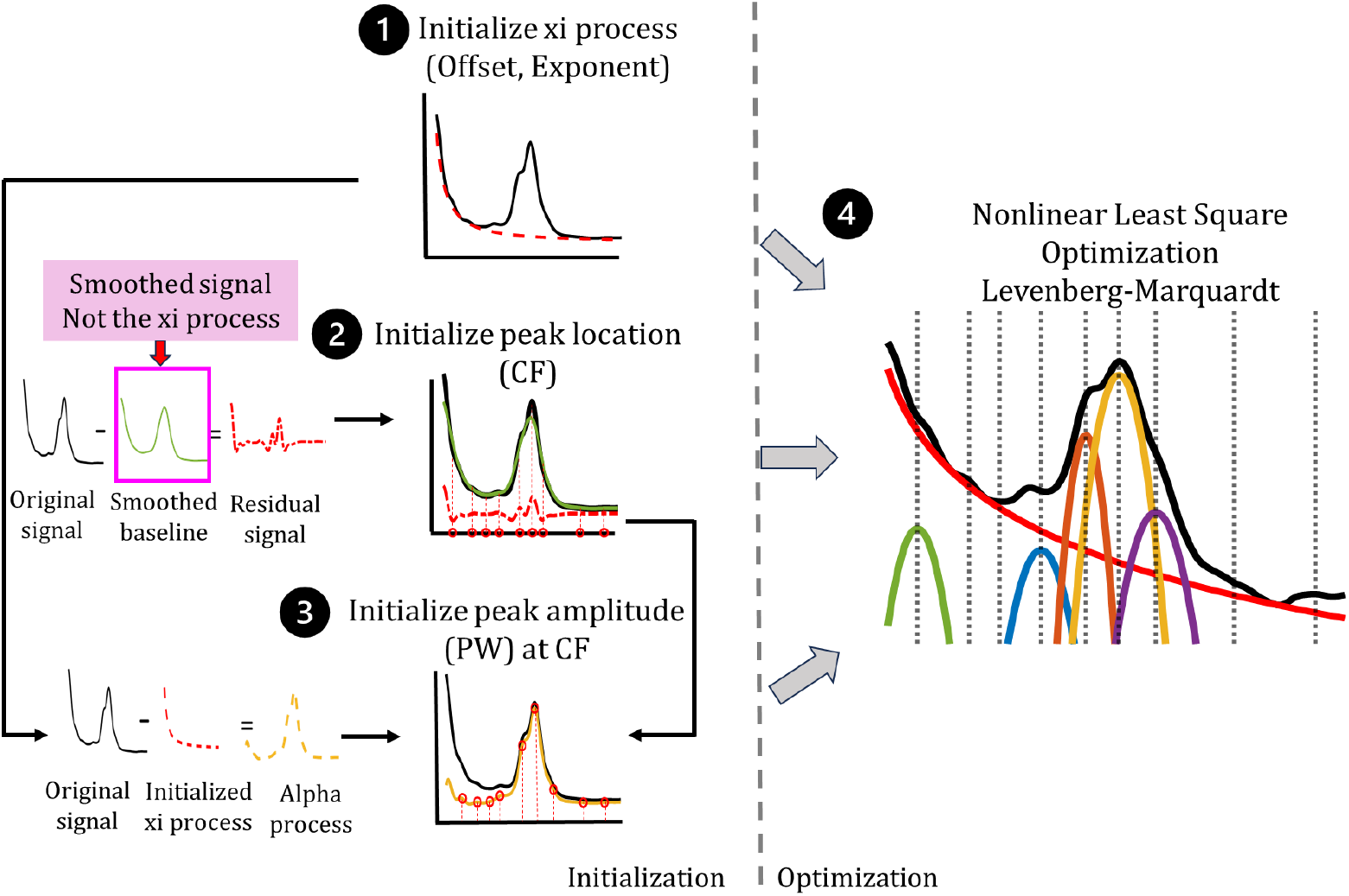
The workflow of robust EEG spectral parametrization

1. **Lorenz curve initialization:** First, perform Lorentzian function parameter inference on the original signal ***s*** using given initialization parameters to obtain curve ***s***_*Lorenz*_;
2. **Gaussian peak location (CF) initialization**: Generate a smoothed signal ***s***_smooth_ from the original signal ***s***, and then **detect peak locations** using the residual signal ***s***_*r*_ = ***s*** − ***s***_smooth_. (first residual). This approach enables peak detection by simply adjusting the smoothing parameters—achieved through non-parametric locally weighted regression. By increasing or decreasing the smoothing bandwidth, one can control the sensitivity of peak detection, capturing more or fewer peaks, respectively. **Importantly, the residual signal *S***_*r*_ **inherently removes underlying trends without introducing high-amplitude, wideband artifacts, making it more robust for identifying true spectral peaks**.
3. **Gaussian peak height (PW) initialization:** for better initialization of PW, we calculate the residual signal with ***s***_*alpha*_ = ***s*** − ***s***_*Lorenz*_(second residual), that the comes ***s***_*Lorenz*_ from step 1. Here, the Gaussian peak location is decided by CF comes from step 2. **Parameters optimization:** Optimize all the parameters with the initial values under the nonlinear least square problem using strategy L-M algorithm. Here, the initialize Gaussian bandwidth parameters-BW are within the 0.8-1 range to account for potential temporal drift and tailing effects of low-amplitude peaks in EEG spectra.

**This Gaussian peak selection method enhances robustness and simplicity by requiring only baseline smoothness half-width adjustment, while residual-based detrending circumvents fitting errors inherent in prior trend estimation approaches**.

Here, we compare our robust parametrization method with FOOOF. The results (check in SI 2.3) show that our robust method achieves a better goodness-of-fit and a higher R^2^ value. Furthermore, applying the FOOOF algorithm did not alter the developmental trajectories of the periodic and aperiodic components found in this study. This may suggest that under resting-state conditions (i.e., without neurotransmitter modulation), the periodic and aperiodic components manifest independently.(Brake et al., 2024).

### 1.5.7 Developmental trajectory function generation

To characterize the non-linear relationship between age and five periodic and aperiodic parameters, we used Locally Weighted Scatterplot Smoothing (LOWESS, statsmodels library, https://www.statsmodels.org/stable/index.html). The analysis was based on data from all electrophysiological leads across 1,563 participants (*N*_*points*_ =1563*18 = 28134). To prevent overfitting and identify the optimal degree of smoothing, the kernel bandwidth was selected by minimizing the Generalized Cross-Validation (GCV) value. A single, unified bandwidth of 2/3 ≈ 0.67 was applied to all models. We estimated the uncertainty of the smoothed trend by deriving 95% point-wise confidence intervals through a bootstrap resampling procedure. Specifically, we generated 1,000 bootstrap datasets by resampling the original data points with replacement. A LOWESS curve was fitted to each bootstrap sample (Davison and Hinkley, 1997), and the final 95% confidence interval was defined by the 2.5th and 97.5th percentiles of the distribution of these 1,000 curves at each point along the age axis.

### 1.5.8 Multimodalities data analysis

We aim to explore the physiological mechanisms underlying the developmental patterns of neuronal periodic and aperiodic signals in the brain by examining the advanced developmental trajectories of neurotransmitters, energy consumption, structural imaging, and cognitive performance, as well as the primary regulatory trajectories of gene methylation levels and telomere length.

#### 1.5.8.1 Periodic components with GABA

Gamma-Aminobutyric acid (GABA) is the main inhibitory neurotransmitter in the human nervous system, playing a crucial role in the functioning of the central nervous system by regulating neuronal firing rhythms(Buzsa ki et al., 2007). EEG studies have shown that enhanced GABAergic inhibition is typically associated with synchronizing brain electrical activity, manifested as increased alpha and theta waves(Buzsaki, 2004). Extensive research has investigated age-related variations in cortical GABA level concentrations measured by magnetic resonance spectroscopy (MRS), revealing marked associations between GABA levels and chronological age throughout maturation, development, and aging processes (Ghisleni et al., 2015; Marenco et al., 2018; Porges et al., 2017; Rowland et al., 2016). However, the age-related trends observed show considerable variability. Building on these findings, Porges et al. (2021) proposed a hypothesis suggesting a nonlinear relationship between GABA levels and age. To investigate this, a lifespan cortical GABA developmental curve was derived using the Penalized Basis Spline model, based on all available MRS data. It was also hypothesized that the resulting optimal curve would align with other established neurodevelopmental markers, such as white volume and EEG power spectra. Thus, we compare the curve from Porges et al. (2021) with the developmental charts of the EEG periodic component parameters including mean and stand deviation. The standard deviations of the developmental trajectories for both GABA and the periodic components show considerable fluctuation around the peak, near the age of 20, indicating substantial individual differences in neuronal maturation at this stage.

#### 1.5.8.2 Periodic components with metabolism

In addition to regulating neurotransmitters, neurons require substantial energy during activity (such as firing and signal transmission), primarily provided by adenosine triphosphate (ATP). The EEG measures the postsynaptic potential activity of pyramidal cells that are synchronously activated in the cortex, and its power spectrum is directly dependent on metabolic processes. Existing data show significant differences in the mass-specific metabolic rate of organs at rest. Although the brain accounts for only 2% of body weight, its metabolic demand is relatively high, with energy consumption at rest accounting for about 20% to 25% of total body energy expenditure, approximately 1000 to 1200 kJ (or 240 to 290 kcal) per day (Castrillon et al., 2023). Pontzer et al. (2021), using a large (n = 6421 subjects; 64% female, ages 8 days to 95 years) and diverse (n = 29 countries) dual-label water measurement database, studied the relationship between total energy expenditure and age in humans (Pontzer et al., 2021). Since the heart, liver, brain, and kidneys together account for 60% of the total basal energy expenditure (BEE) in adults, and the BEE of these organs aligns with the overall BEE changes with age, we use the total body BEE as an indirect measure of brain energy consumption for analysis.

Brain activity is not only associated with energy changes but is also closely linked to the total body water (TBW) and water turnover rate (WT). Water is one of the most important components of the brain, accounting for approximately 2% of the body weight, yet its content in the brain is as high as 75%-80%. The role of water in the brain extends beyond providing physical support (e.g., maintaining cell shape and structure) to include maintaining electrolyte balance between neurons and glial cells, buffering heat generated during neural activity, and participating in metabolic processes (such as the synthesis and transmission of neurotransmitters)(Go, 1997; MacAulay, 2021; Pham et al., 2024). Water turnover, the rate at which water is recycled within the body, is closely associated with cerebral blood flow, brain cell activity, neurotransmitter activity, and metabolic rate. Yamada et al. (2022) used the isotope-tracking (^2^H) method to investigate fluid turnover rates in 5,604 individuals from 23 countries, ranging in age from 8 days to 96 years. They constructed functions relating total body water (TBW), water turnover (WT) (non-athletes), and other related metrics to age.

Here, we used the data of BEE and WT from paper Pontzer et al., 2021 and Yamada et al., 2022, with local weight kernel regression build the developmental trajectory of BEE and WT with age, and compare these curves with aperiodic and periodic developmental trajectory. Notably, since basal energy expenditure (BEE) and fat-free mass (FFM) exhibit an allometric (power-law) relationship, and water turnover (WT) is influenced by total body water (TBW), we further examined the association between the developmental trajectories of FFM-adjusted BEE and WT/TBW and those of EEG-derived metrics.

#### 1.5.8.3 Periodic components with MRI

In addition to physiological regulation, brain activity is closely linked to brain structure. The brain’s white matter is primarily composed of myelinated nerve fibers, which are responsible for connecting different brain regions to facilitate information exchange and communication(Chorghay et al., 2018).. White matter’s integrity and functional state directly affect the cortical electrical activity patterns. White matter contains myelinated and unmyelinated axons and glial cells, including oligodendrocytes that produce myelin, microglia, astrocytes, and oligodendrocyte precursor cells. Myelin serves as an electrical insulator for axons, enabling the rapid saltatory conduction of action potentials and increasing the speed of neural signal transmission (de Faria et al., 2021). Due to neural plasticity, the white matter (WMV) volume undergoes continuous changes throughout a person’s life. Smaller WMV is considered to be associated with attention, declarative memory, executive function, and intellectual impairments, reflecting the overall morphological integrity of the brain’s white matter (Chorghay et al., 2018; Kiely et al., 2022b) as obtained through MRI T1 imaging.

In this study, we compare the developmental trajectory of human brain WMV, derived from a multicenter study published by Bethlehem et al (Bethlehem et al., 2022), with the developmental patterns of both periodic and non-periodic components identified in our analysis. At the same time, considering the functional metrics of white matter, it is important to further investigate the health and integrity of myelin. In this context, we examine two key indicators R1 and myelin water fraction (MWF). For white matter, R1 (1/T1) is primarily influenced by changes in myelin content and reflects the longitudinal relaxation rate of water proton signals in a magnetic field, providing complementary information for assessing fiber diffusion properties. Here, we fitted R1 data from Jason D. Yeatman and colleagues, collected across the lifespan from 7 to 85 years, and plotted the developmental trajectory of R1 across stages of maturation, development, and aging (Yeatman et al., 2014). The R1 developmental trajectory is built based on the model *y* = *w*_1_ × *age*^2^ + *w*_2_ × *age* + *w*_3_. Here we extractsoem example trajectories of Occipital callosum, Right SLF, Ant. frontal callosum and Orbitofrontal callosum. The details of the parameters of trajectory model and other trajectories can check the site https://github.com/jyeatman/lifespan.

R1 reflects the overall relaxation properties of tissue and is influenced by various non-myelin factors, including total tissue water content, protein concentration, iron deposition, and inflammation (MacKay and Laule, 2016b). Additionally, confounding factors such as edema or gliosis may artificially elevate R1, thereby obscuring genuine myelin-related changes. Experimental validation has demonstrated that MWF exhibits a strong correlation with histological myelin density (e.g., Luxol Fast Blue staining; (Laule et al., 2006)). Consequently, we further compared the developmental trajectories of MWF.

Advanced MRI techniques based on multi-component relaxation (MCR) provide sensitive and specific measurements of changes in myelin water content, offering a valuable measure of demyelination in brain tissue. Based on the work of Matthew Kiely and colleagues(Kiely et al., 2022a), we further compared the CF and PW curves with the developmental trajectory of myelin water fraction (MWF). Here, the function of whole brain MWF with age *y* = 4.33 × 10^−10^ × *age* + 2.73 × 10^−5^ × *age*^2^ and FA trajectory model *y* = 1 × 10^−15^ × *age* + 8.65 × 10^−5^ × *age*^2^ are used in this study. For other modality trajectories can check paper (Kiely et al., 2022a) in detail.

#### 1.5.8.4 Periodic components with Cognitive

Neuroscience data ultimately reflects the cognitive abilities of the human brain, which are central to individual consciousness. In recent years, with the development of neuroscience, research on age-related cognitive changes and their cognitive and neural mechanisms has transitioned from being independent to a more mutually validating approach. Researchers have sought to link physiological and anatomical abnormalities with cognitive behavior measurements, aiming to uncover more direct and robust evidence of cognitive changes (Baltes et al., 2007; Bartzokis, 2004; Klimesch, 2012; Lindenberger, 2014; Qiu et al., 2024). Studying the age-related patterns of cognitive abilities based on neuroscience data enables a better understanding of cognitive function changes, helps address aging issues, and informs related educational strategies (Germine et al., 2011; Hertzog et al., 2008). However, the relationship between cognitive abilities and age-related changes with neuroscience findings remains largely unknown (Gallego-Rudolf et al., 2024; Wischnewski et al., 2023). Cognitive abilities include crystallized intelligence, which evaluates the knowledge and skills accumulated through experience, learning, and education, and fluid intelligence, which assesses an individual’s ability to quickly process information, solve problems, and engage in abstract reasoning in novel situations. These are typically measured through vocabulary tests, reasoning tasks, and pattern recognition tests.

In this study, we build upon the dual exponential model from McArdle et al. (2002a), which is based on data collected in 1988. According to the Gc-Gf theory, this research includes 1,193 subjects, aged 2 to 95 years, and constructs a model involving 11 key factors for broad cognitive abilities from the WJ-R test. The dual exponential model used here is as follows, and the detail value of the parameters can check the paper McArdle et al. (2002a).

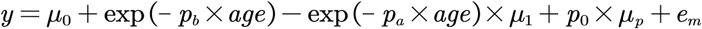

#### 1.5.8.5 Aperiodic components with Telomere

Telomeres are DNA repeat sequences located at the ends of chromosomes, playing a crucial role in protecting chromosome stability and preventing wear and tear at the chromosome ends (Vaiserman and Krasnienkov, 2020). Since the replication mechanism cannot fully replicate the very ends of chromosomes, each time a cell divides, the telomere region shortens until its length is no longer sufficient to protect the chromosomes. At this point, the cell may enter a senescent state, stop dividing, or die. Therefore, telomere length (TL) is considered to limit the number of cell divisions and acts as a “mitotic clock” in cells (Olovnikov, 1996). TL is also associated with various age-related diseases, such as Alzheimer’s disease, cancer, and heart disease (Fani et al., 2020; Forero et al., 2016; Notterman and Schneper, 2020). Numerous studies have shown that short leukocyte telomere length (LTL) reflects the overall body TL in other tissues and is a potentially useful biomarker for healthy aging. However, it remains unclear whether LTL in leukocytes correlates with TL in tissues with low proliferative activity, such as neurons (Mather et al., 2011; Vaiserman and Krasnienkov, 2020). Blood leukocytes are a highly heterogeneous cell population, including various cell types, and their composition is highly variable, easily influenced by stress exposure and other factors (Koliada et al., 2015). Therefore, to explore the relationship between telomere length and neural cells, we compared the developmental trajectories of telomere length in different leukocyte subpopulations with the developmental trajectories of cyclic and non-cyclic EEG parameters, based on the study by Aubert et al., 2012. LTL data were obtained through flow cytometry-based FISH measurement, comprising 835 healthy individuals aged from birth to 102 years. The leukocyte population includes lymphocytes, granulocytes, CD20+, CD45RA+, CD20, CD45RA2, and CD57+ and the LTL developmental trajectory is constructed based on Local weighted kernel regression (LOWESS).

#### 1.5.8.6 Aperiodic components with DNA methylation

In reliable biological age biomarker studies, in addition to telomere length, epigenetic clocks have become a recent research hotspot. By measuring the changes in DNA methylation levels at specific loci with age, several biological age prediction calculators have been developed, such as the Horvath and Hannum clocks (Hannum et al., 2013; Horvath and Raj, 2018) DNA methylation refers to the process of adding a methyl (-CH3) group to specific positions in the DNA molecule. It is an important modification in epigenetics, involving gene expression regulation without altering the DNA sequence. In mammals, DNA methylation primarily occurs at CpG islands, which are typically located in the promoter regions of genes and control gene initiation and expression. DNA methylation can suppress gene expression, while demethylation is usually associated with gene activation (Horvath and Raj, 2018). However, the large-scale epigenetic age studies currently available primarily focus on adults, and there is limited knowledge about studies involving children (Horvath, 2013; Jones et al., 2015; Yang et al., 2023).

To explore the relationship between brain rhythms and epigenetic factors, we analyzed the age-related correlations of the Beta Prime values from peripheral blood DNA of 398 boys aged 3–17 years (GSE27097) collected by Barwick Lab, Emory University, and 192 Irish individuals (GSE19711) and 540 adult females (GSE20067) collected by the UCL Cancer Institute/Institute for Women’s Health, University College London. The curve of DNA methylation is built based on LOWESS.

## 1.6 Data available

EEG cross-spectrum: https://doi.org/10.7303/syn26712693;

The other modality dataset will be available after formal publication.

## 1.7 Code available

1. Harmonized z-scores for external dataset (https://github.com/LMNonlinear/HarMNqEEG).
2. The spectrum parametrization toolbox in this study will be made publicly available in a GitHub repository upon formal publication of this article, and the link will be provided.

## 1.8 Support Information

Will be available after formal publication

## 1.9 Acknowledgements

Thanks for Eric C Porges, Greg Jensen, Britton Mark for providing the gaba dataset and helping on the code for gaba trajectory generation.

## Chapter 1 : Tables

**Table S1:**
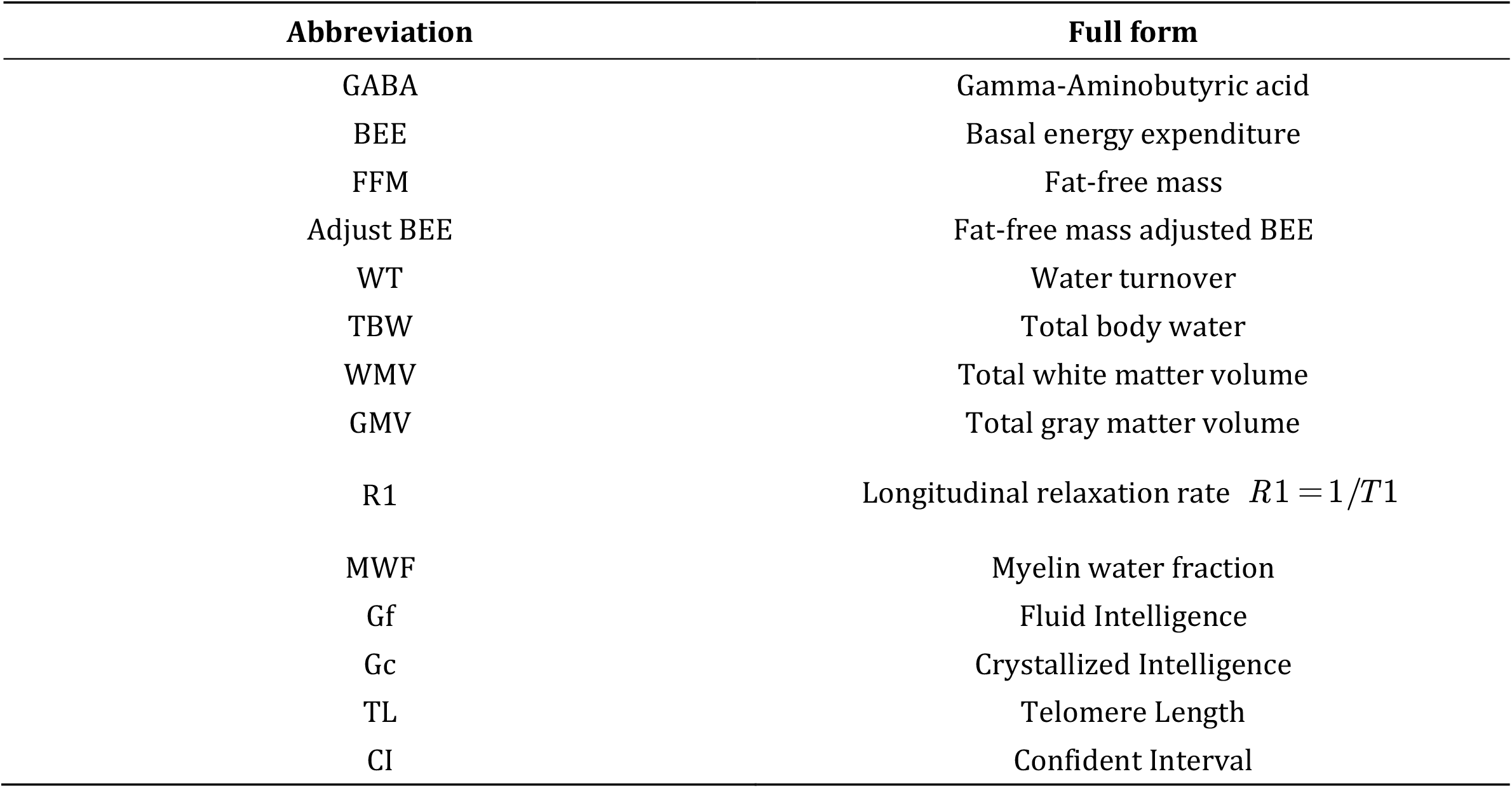
Abbreviation table.

| Abbreviation | Full form |
| --- | --- |
| GABA | Gamma-Aminobutyric acid |
| BEE | Basal energy expenditure |
| FFM | Fat-free mass |
| Adjust BEE | Fat-free mass adjusted BEE |
| WT | Water turnover |
| TBW | Total body water |
| WMV | Total white matter volume |
| GMV | Total gray matter volume |
| R1 | Longitudinal relaxation rate $R1 = 1/T1$ |
| MWF | Myelin water fraction |
| Gf | Fluid Intelligence |
| Gc | Crystallized Intelligence |
| TL | Telomere Length |
| CI | Confident Interval |

**Table S2:** Comparison of R^2^ Values for Linear, Nonlinear, and Simplified Models in Characterizing Parameter Developmental Trajectories.

|  | Exponent |  |  | Offset |  |  | CF |  |  | PW |  |  | BW |  |  |
| --- | --- | --- | --- | --- | --- | --- | --- | --- | --- | --- | --- | --- | --- | --- | --- |
| R<sup>2</sup> | 0.41 | 0.52 | 0.51 | 0.26 | 0.33 | 0.33 | 9.5-5e | 0.22 | 0.22 | 0.005 | 0.01 | 0.02 | 0.01 | 0.05 | 0.02 |

linear| nonlinear | piecewise linear

## Chapter 2 EEG log-spectrum distribution and batch harmonization

### 2.1 The Age-Related Distribution Characteristics of EEG Power Spectrum along all Frequency Points

**Figure S1:**
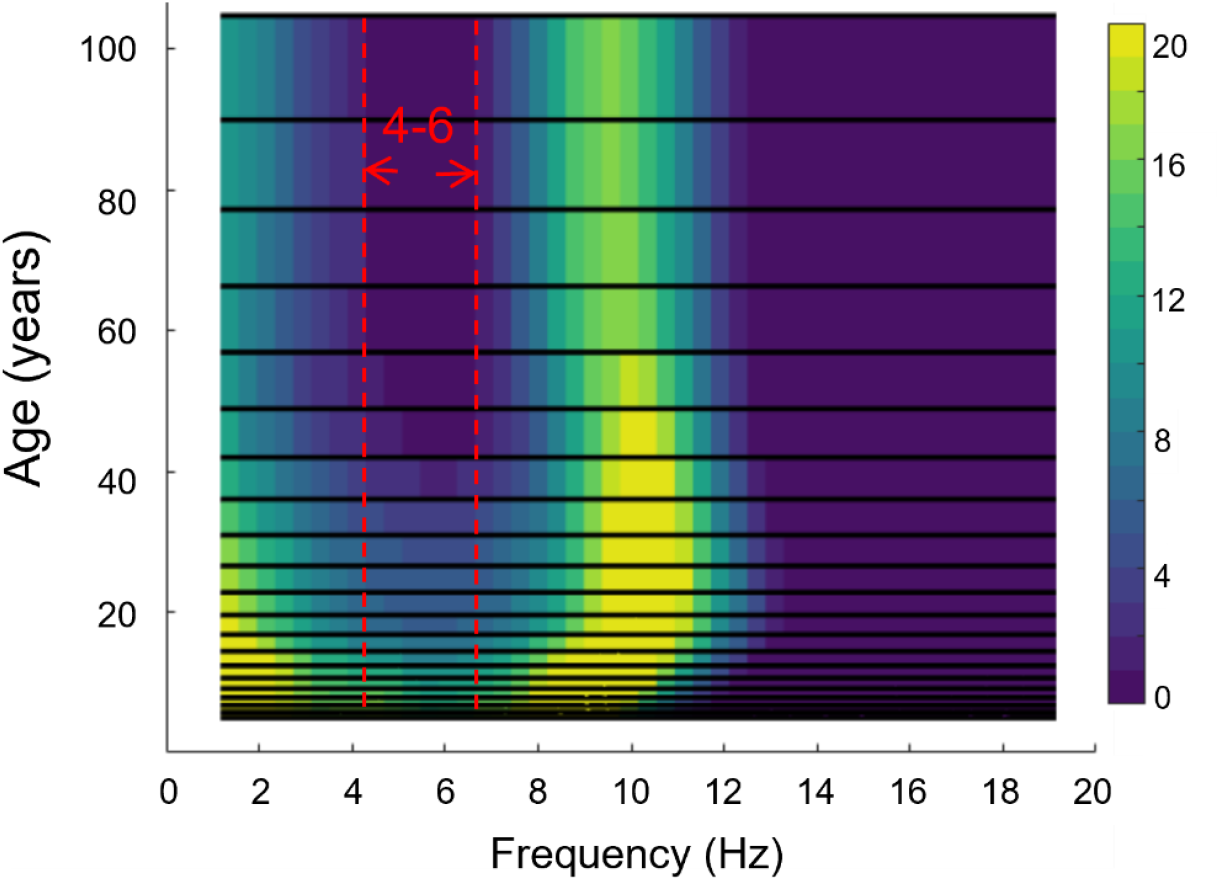
Age Distribution of EEG log-spectrum in different frequency bands. The figure is adapted from the paper Li et al., 2022, presented in a planform view. To better delineate the boundaries, the limitation of the colorbar was narrowed. It can be observed that under resting state with eyes closed: 1) The log-spectrum across all age groups exhibited the highest power within the 6-14 Hz range; 2)A gradual decline in log-spectrum power with aging was observed in the 1-4 Hz (delta band), with a notable transition around 20 years of age; 3) Across all age groups, the log-spectrum exhibited the lowest power in the 4-6 Hz and 14-20 Hz frequency ranges. Compared to the conventional 4-8 Hz (theta band), the 4-6 Hz sub-band demonstrated significantly greater age-related differentiation in spectral characteristics. Considering the attenuated power patterns in both 4-6 Hz and 14-20 Hz (beta), this study operationalizes 1-4 Hz (delta), 4-6 Hz (theta), 6-14 Hz (alpha), and 14-20 Hz (beta) as the revised frequency band classification criteria.

According to the spectral characteristics of EEG in different typical states, frequencies are usually divided from low to high into six frequency bands: delta (1–4 Hz), theta (4–8 Hz), alpha (8–13 Hz), beta (13–30 Hz), low-gamma (30–70 Hz), and high-frequency oscillations high-gamma (>70–150 Hz) (Tatum, 2014), and each frequency band reflects different neural oscillation modes. In the resting-state eyes-closed condition, the amplitude of alpha waves significantly increases, which is related to reduced visual input and a relaxed state. In very few individuals, theta and beta waves may also appear when eyes are closed, where beta is associated with mild thinking, meditation, and light sleep, and theta may be in those who remain highly cognitive and alert even with eyes closed (Cahn and Polich, 2006; Schmidig et al., 2024). However, whether this phenomenon is also related to age is still unknown. According to our previous research on the EEG log spectral brain developmental surface, children have higher delta components compared to adults, and beta energy is almost absent in all age groups. To better observe the distribution of theta and beta with age, we appropriately lowered the EEG power spectrum threshold (Figure S1) and found that both theta and beta decrease with increasing age, and theta is generally lower than delta in energy, that is, on the spectral-age-frequency figure, there is a 4–6 Hz gap, meaning that middle-aged and elderly people have much less low-theta components compared to children, which aligns with the conclusion that beta waves are suppressed in the resting-state eyes-closed condition and that children have more light sleep components (He et al., 2019; Lomas et al., 2015). This inspires the basis for frequency band division in different age groups. However, since delta and theta are in low frequencies, it is still unclear whether the distribution of delta and theta energy with age is affected by aperiodic components. Therefore, to more comprehensively explore the age dependency of different frequency bands, we need to separate periodic and aperiodic components.

### 2.2 Batch harmonization result

**Figure S2:**
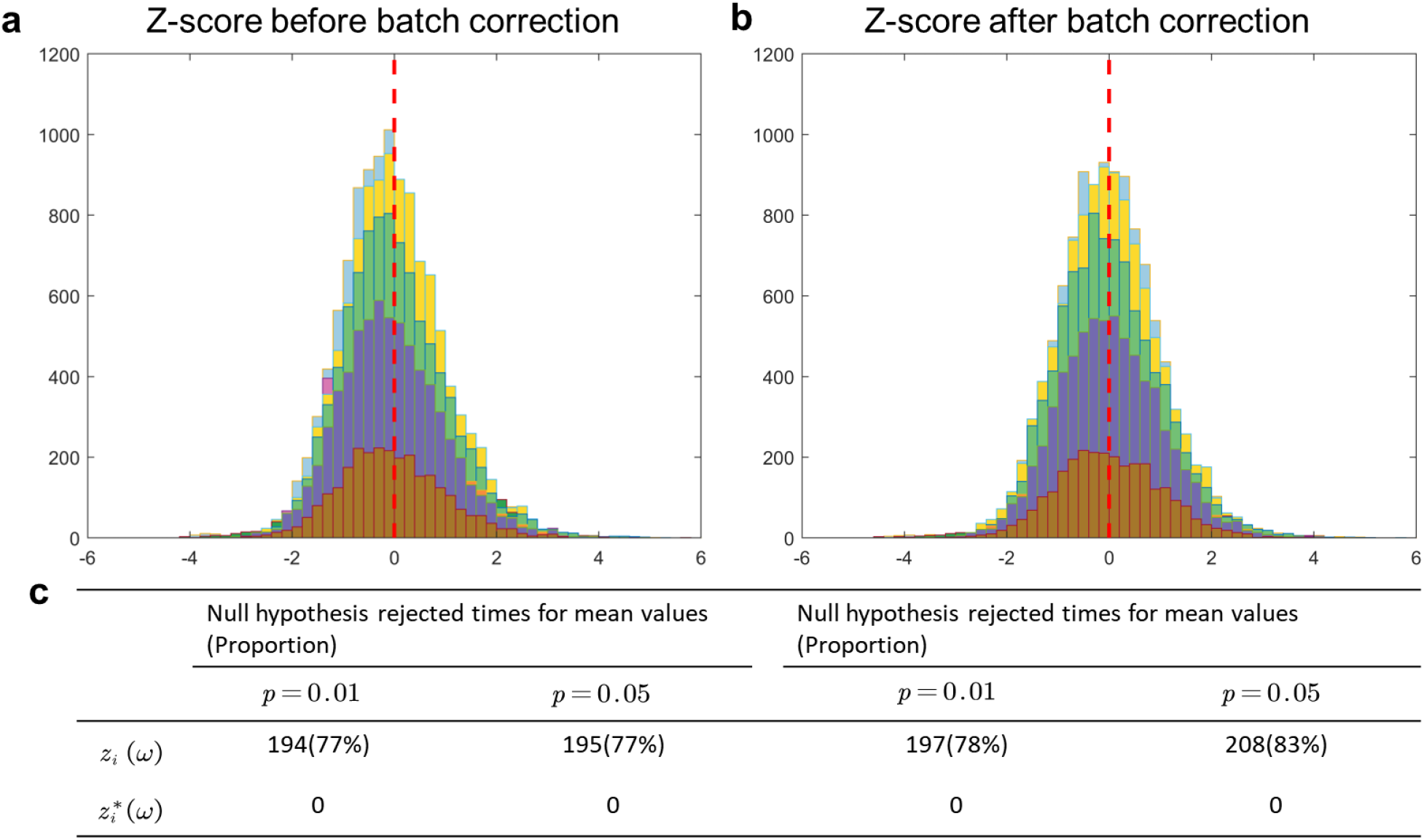
Distribution of EEG log-spectral z scores of batches before and after batch harmonization. a: The distribution of batch z-scores before harmonization, the mean value of z-scores is not equal to 0; b: The distribution of batch z-scores after harmonization, the mean value of z-scores is equal to 0; c: Summarization of the zero mean and standard variance test results of z-scores before and after harmonization. By applying the MATLAB function ttest, there are 77% and 78% variables that do not have a zero mean and standard deviation, after harmonization, all the hypotheses are met.

### 2.3 Method comparison with FOOOF of EEG spectral parameterization

To validate the robustness of the trajectories, we parameterized the EEG spectra into periodic and aperiodic components using the FOOOF algorithm (“peak_width_limits”: [0.5, 12.0], “max_n_peaks”: Infinity, “min_peak_height”: 1e-05, “peak_threshold”: 2.0, “aperiodic_mode”: “fixed”) and subsequently plotted the developmental curves for each component. As shown in Fig S3, the developmental trends of the periodic and aperiodic components were not significantly altered, with the only subtle difference appearing in the bandwidth (BW) around 20 years of age.

**Figure S3:**
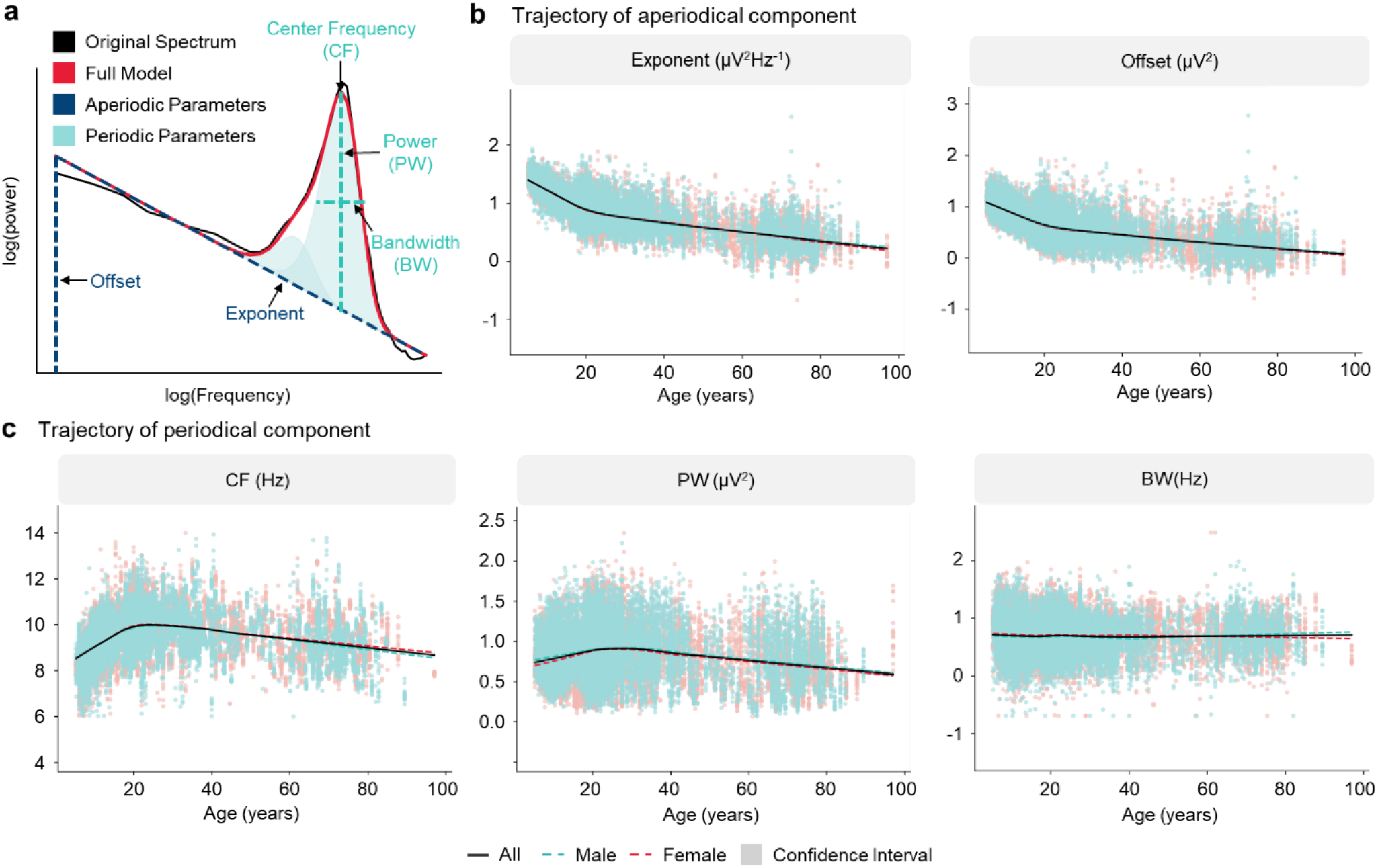
Developmental Trajectories of EEG Aperiodic and Periodic Components Based on FOOOF. a: The FOOOF spectral decomposition algorithm, which operates on the log-power spectrum. Here, the frequency axis is plotted on a logarithmic scale to visualize the aperiodic component without a “knee”. b: Developmental trajectories of the aperiodic parameters: Exponent and Offset. c: Developmental trajectories of the periodic parameters: Center Frequency (CF), Power (PW), and log-transformed Bandwidth (log(BW)).

To compare the robust spectral parametrization and FOOOF algorithms, we compared the goodness-of-fit, AIC (Akaike Information Criterion), and BIC (Bayesian Information Criterion) of the two methods. As shown in the Cumulative Distribution Function of R^2^ values, for RSP 95% of R^2^ values exceeded 0.91 (Figure S4 a), but for FOOOF, 95% of R^2^ values exceeded 0.87 (Figure S4 b. To balance model complexity, we also compared the AIC and BIC values of the two models. Compared to FOOOF, RSP has lower AIC and BIC values (Figure S4 c-d), indicating that the RSP fit is more robust (avoiding overfitting and underfitting) and has the minimum information loss.

**Figure S4:**
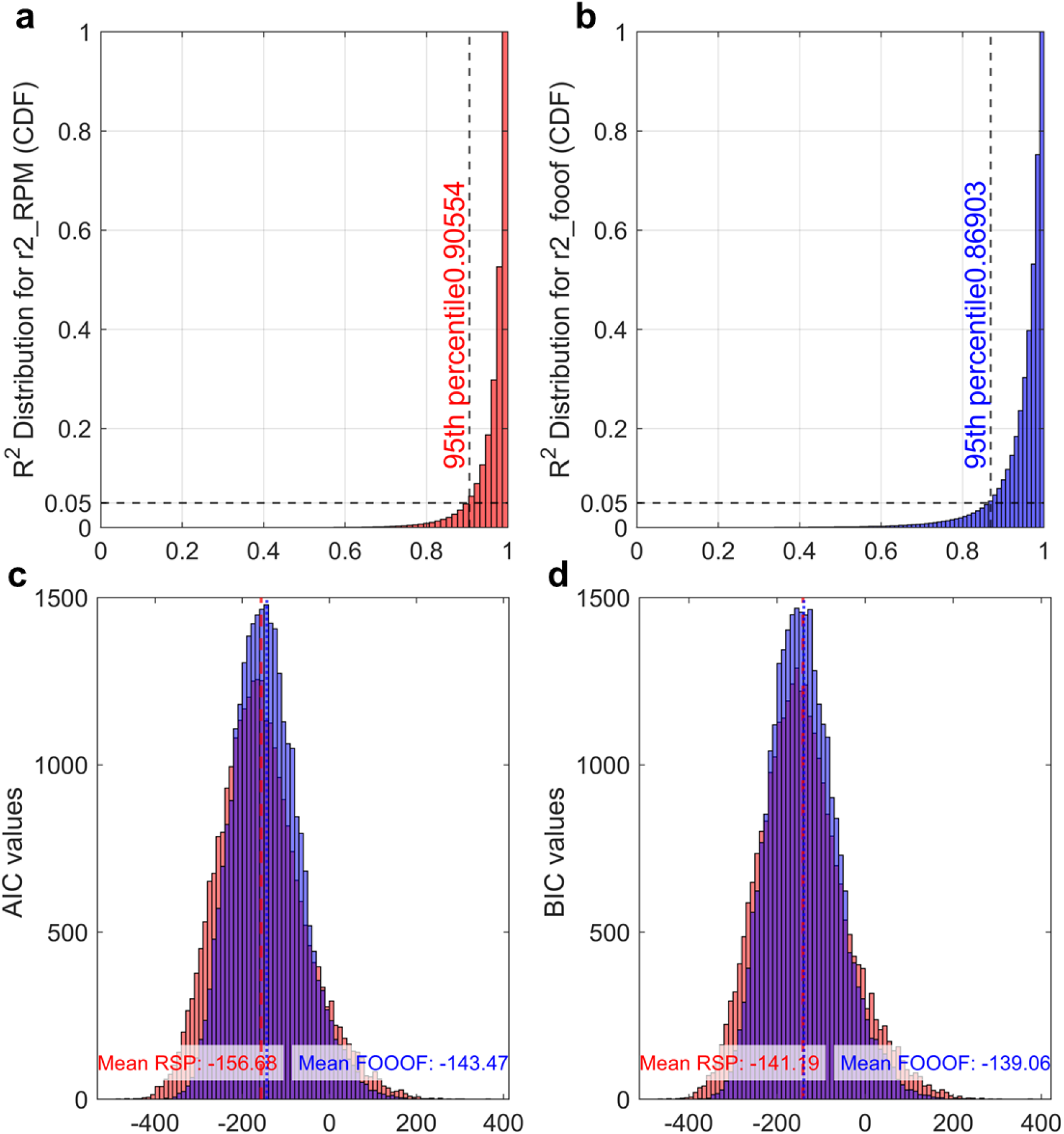
Cumulative distribution function (CDF) of R^2^ values and ACI, BIC values for RSP and FOOOF. a-b: CDF of R^2^ of RSP and FOOOF, Horizontal black dashed line is the 5th percentile of the distribution, and it interest at *R*^2^ = 0.9 (Vertical black dashed line). This means that only 5% of observations have for RSP and FOOOF. c: AIC values for *R*^2^ < 0.91 RSP (red) and FOOOF (blue). The AIC mean value for RSP is −156.68 and −143.47 for FOOOF. d: BIC values for RSP (red) and FOOOF (blue). The BIC mean value for RSP is −1441.19 and −139.06 for FOOOF.

## Chapter 3 Developmental trajectory of EEG spectral components

### 3.1 Linear Developmental Trajectory of EEG Aperiodic and Periodic Parameters

**Figure S5:**
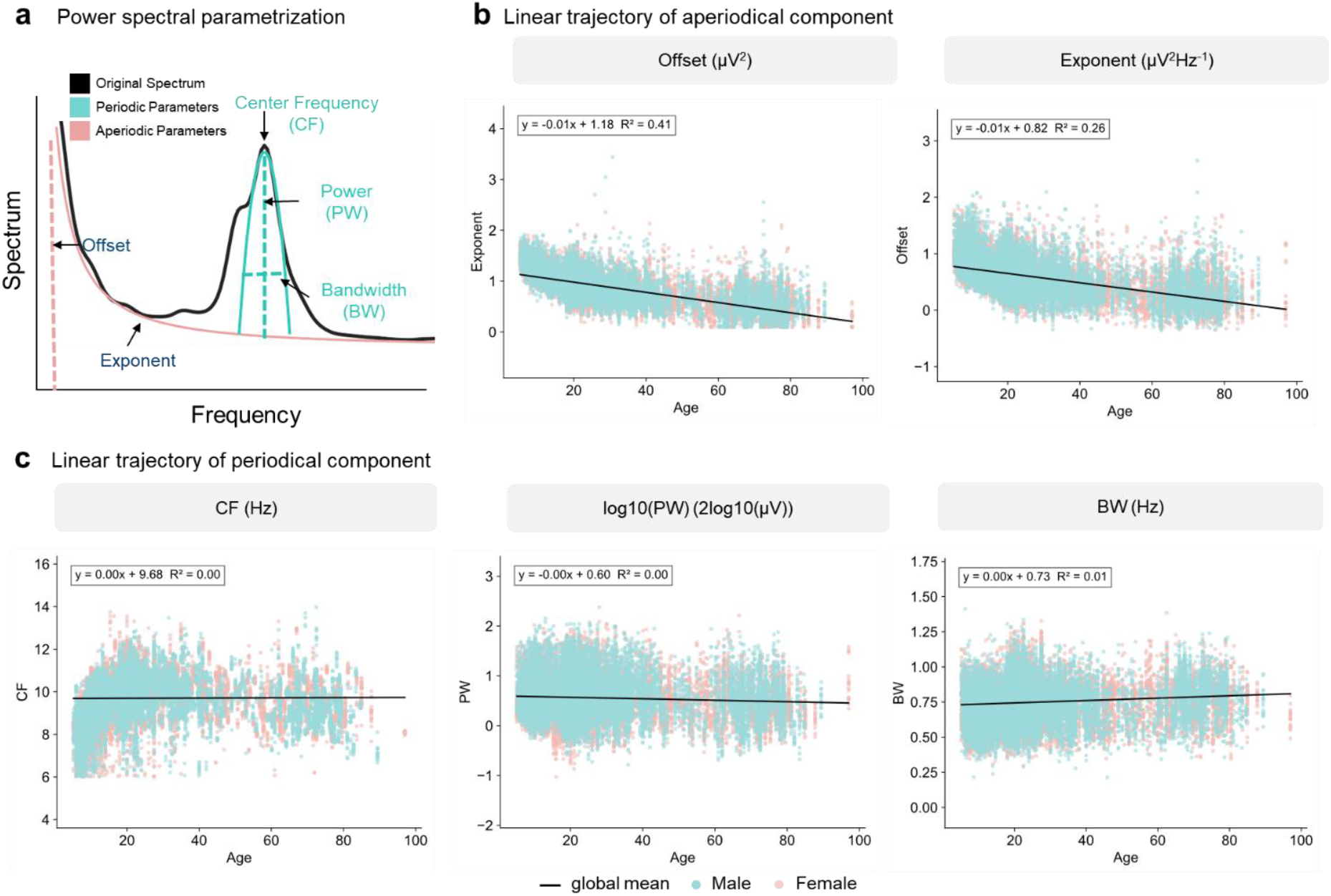
Linear Developmental trajectory of spectral parameters. a: With the same parameterization method, obtain Exponent, Offset, CF, PW and BW. b: Linear developmental trajectory of aperiodic parameters. The R^2 is 0.41 and 0.26 for Exponent and Offset separately; c: Linear developmental trajectory of periodic parameters. The R^2 is 0, 0.005 and 0.05 for CF, PW and BW. Blue solid scatter represents the parameter of males, and pink solid scatter represents the parameter value of females, the black solid line is the global mean value, and the gray area is the confidence interval.

### 3.2 EEG spectral periodic components of delta, theta and beta bands

In this study, beyond the widely researched alpha band, we also examined the parameter distribution of other periodic components across different frequency bands. Figure S6 shows that: 1): In adults, periodic components become increasingly sparse (lacking peaks) outside the alpha range. Center frequency (CF) detections decline markedly after approximately 30 years of age, with fewer prominent peaks observed in higher frequencies (Figure S6 a, d, g). Children exhibit a greater prevalence of low-frequency periodic activity. Between 1–4 Hz, children show stronger periodic power (PW), which declines rapidly before age 20 and stabilizes thereafter (Figure S6 b, e). Additionally, bandwidth (BW) values in this range are elevated compared to adolescents (Figure S6 c, f), suggesting broader spectral peaks during early development. Adults over 20 years of age demonstrate increased high-frequency components relative to younger individuals. Specifically, PW in the 14–20 Hz range increases after age 20 (Figure S6 h), accompanied by a gradual widening of peak bandwidth (BW) (Figure S6 i), indicating enhanced spectral power and dispersion in this frequency range with maturation.

**Figure S6:**
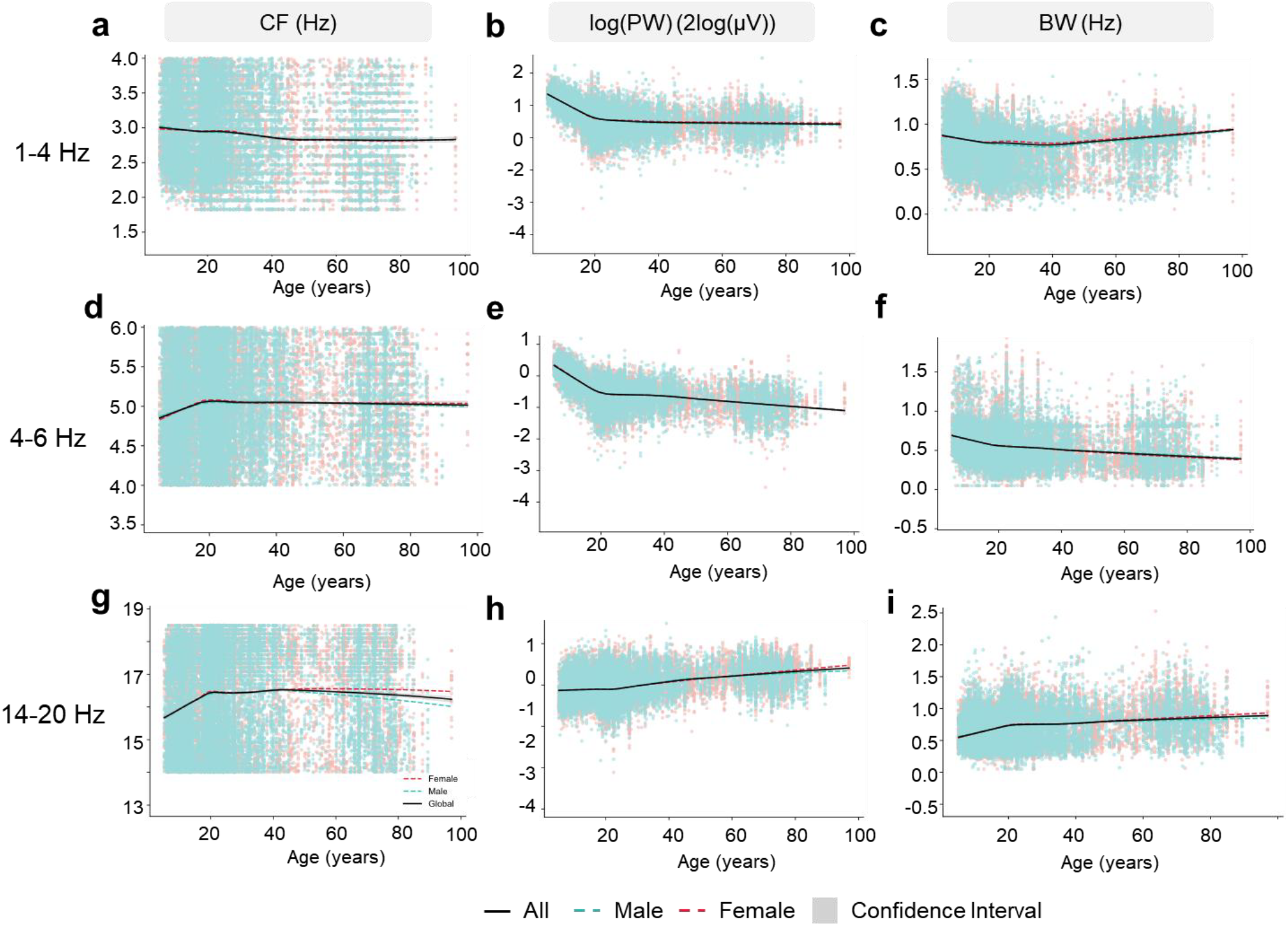
The developmental trajectory of periodic components on delta, theta and beta bands. Panels a-c show the parameter changes in the 1-4Hz range, panels d-f show the parameter changes in the 3-6Hz range, and panels g-i show the parameter changes in the 4-20Hz range.

### 3.3 The development trajectory of log-variance of aperiodic and alpha periodic components

**Figure S7:**
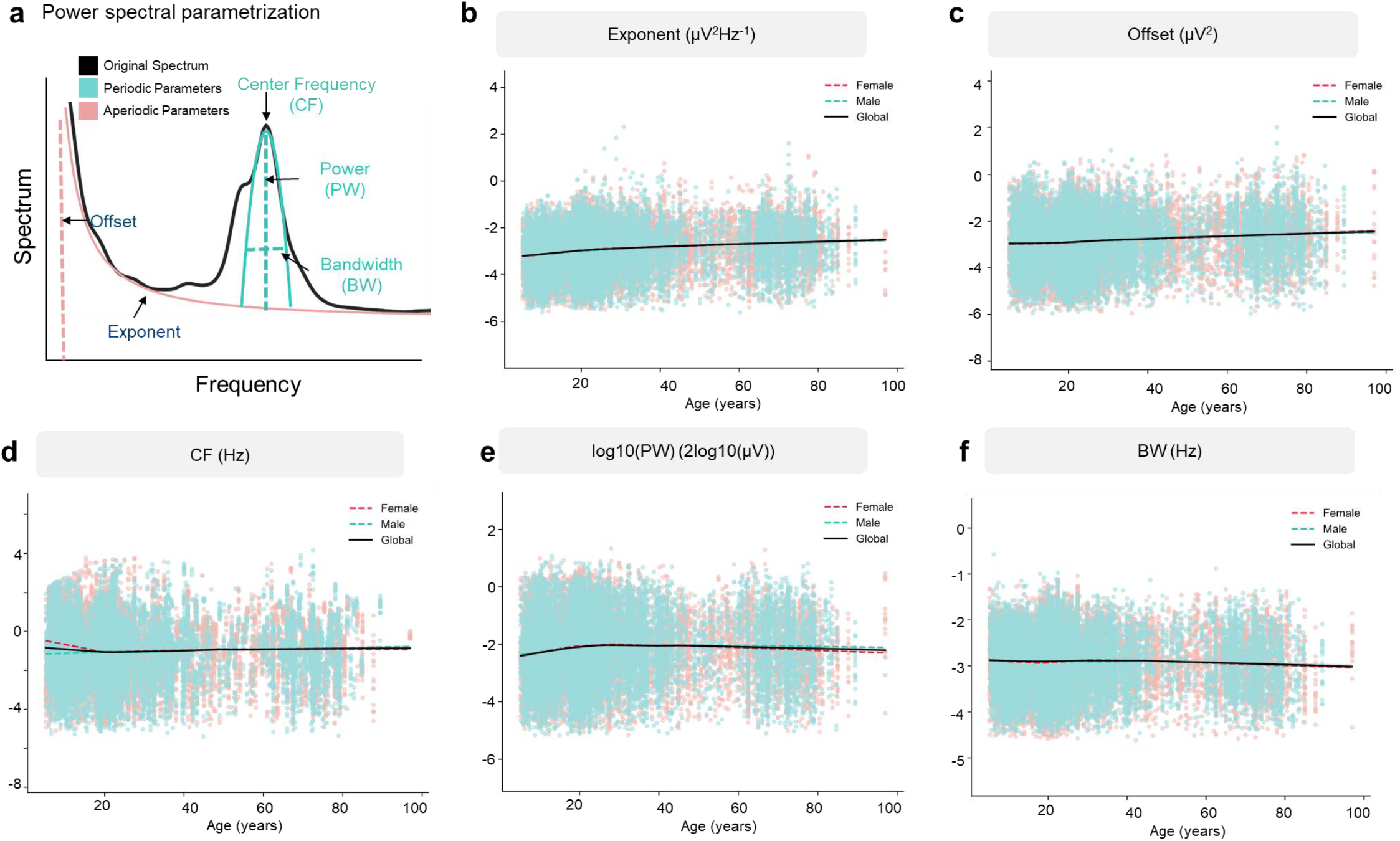
The development trajectory of log variance of aperiodic and alpha periodic components. Unlike the fitted curves for the mean values of the parameters, the variance of all components undergoes slight changes before the age of 20 and remains relatively stable thereafter.

### 3.4 The physical space explanation of developmental trajectory

**Figure S8:**
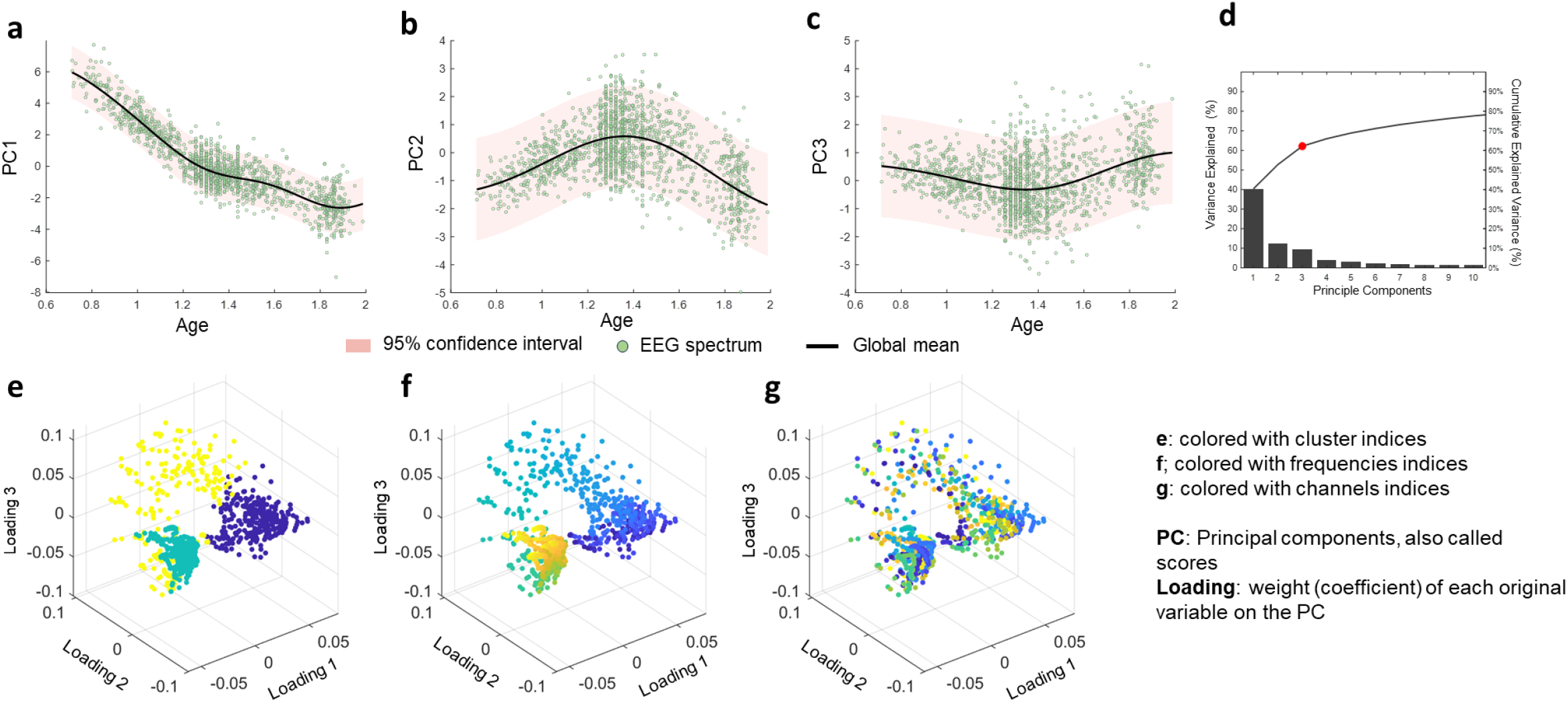
The developmental trajectory of the first 3 main components on a log scale of age. The main components are calculated with principal component analysis (PCA) of the EEG log-spectrum, which embeds all electrodes and frequencies into a low-dimensional space. d: The first three principal components (PCs), which capture the dominant patterns of variance (68%), show distinct developmental trajectories when plotted as functions of log(age). a: PC1 aligns with the developmental pattern of the aperiodic component; b: PC2 mirrors the trajectory of the alpha-band (periodic) activity; and c: PC3 reflects a composite of the remaining periodic components across other frequency bands. These components summarize the joint spatial–spectral organization of brain rhythms across the lifespan. The loadings reveal that the primary distinctions among these three patterns are based more on frequency than on electrode location. Specifically, the clustering results (e) are more consistent with frequency modes (f) rather than the arrangement of electrodes (g), indicating that the observed developmental trajectories are predominantly driven by changes in spectral composition rather than spatial distribution across the scalp.

### 3.5 The turning points and simplified developmental trajectory of PW and Offset

**Figure S9:**
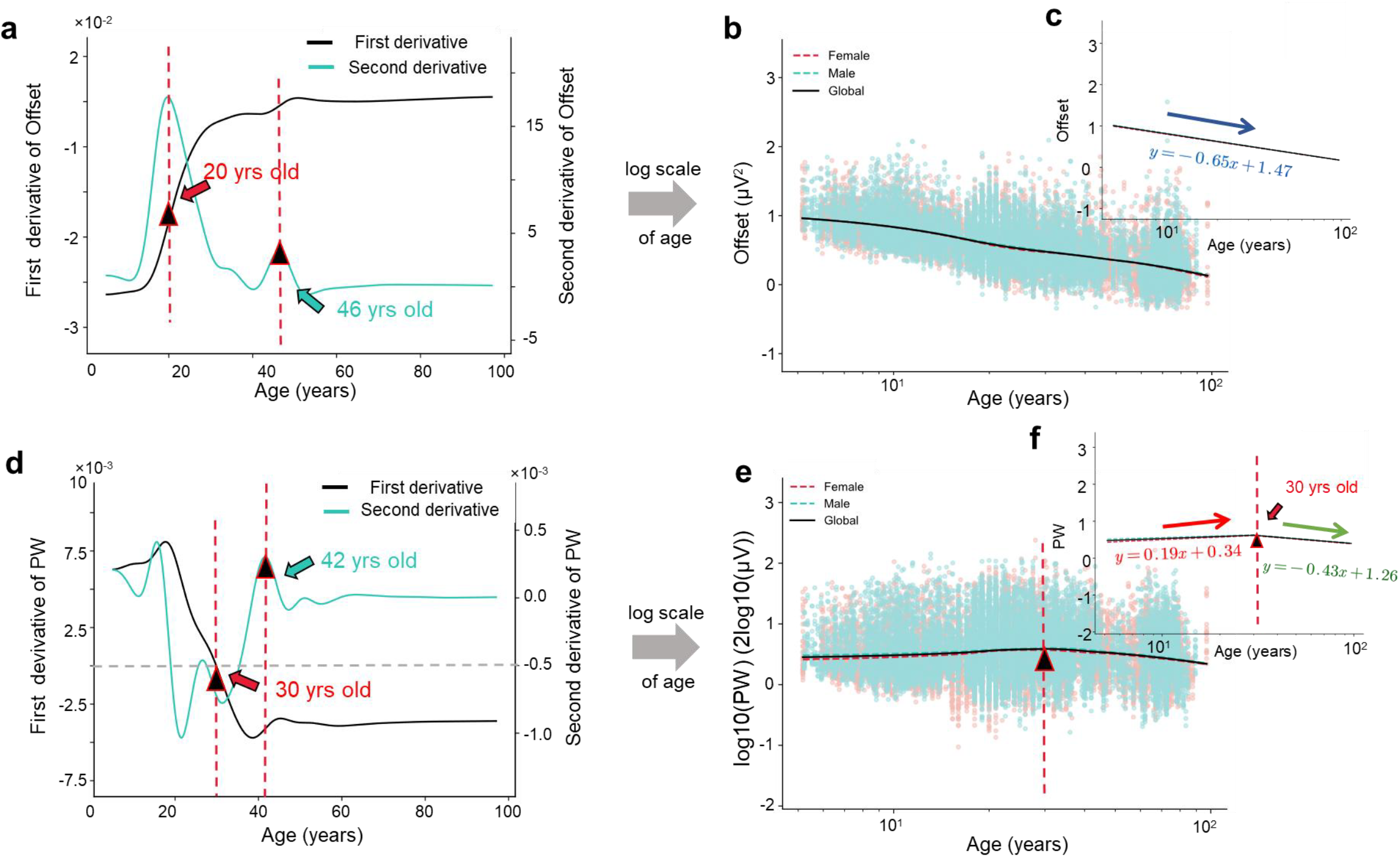
Developmental Trajectories of Offset and log(PW) on the Log-age scale. b: When plotted against log-transformed age, the Offset parameter shows a nearly perfect linear decline over log(age) (d, R^2 is 0.33 at both natural and log age scale). c: Peaking at around 30 years of age (first derivative is 0) and development stabilizes at around 42 years of age (second derivative equal to 0).d: In contrast, the developmental trajectory of log(PW) exhibits an approximately symmetric, inverted U-shape, This pattern is markedly more linear (R^2 is 0.02) and structured compared to the trajectory observed on the natural age scale (R^2 is 0.01)., d: With a notable inflection point occurring around 20 years of age when viewed on the chronological (natural) scale (first derivative is 0) and a stable period of development begins at approximately 46 years old (second derivative equal to 0).

## Chapter 4 The scalp topography of EEG spectral parameters and energy

### 4.1 The scalp topography of parameters on turning points of delta, theta and gamma bands across all channels

**Figure S10:**
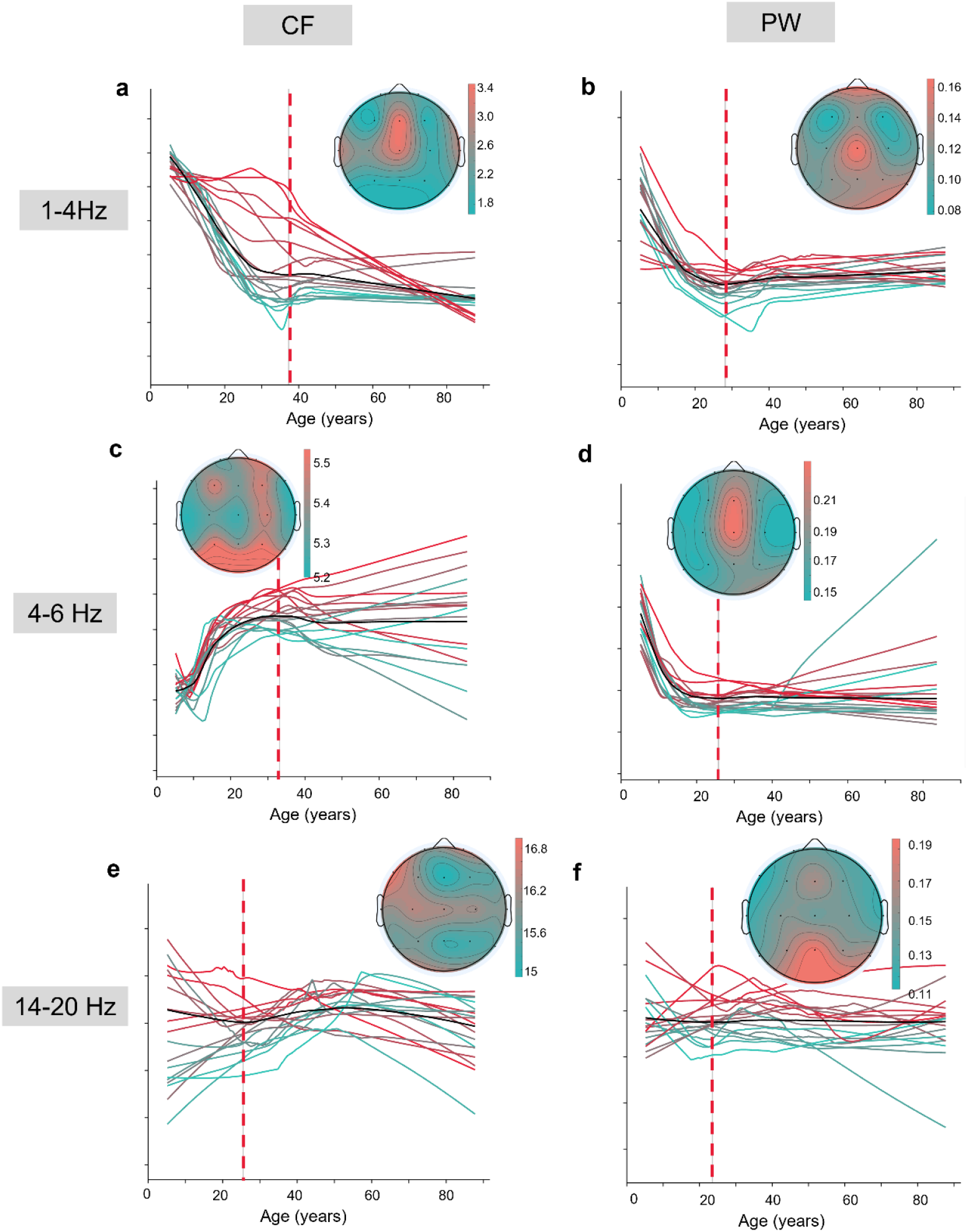
The developmental trajectory of CF and PW on delta, theta and gamma bands across all channels. The topography shows spatial distribution at the age of the turning point of the parameter developmental trajectory. The trajectory are colored based on the age of turning points.

### 4.2 The scalp topography developmental trajectory of Offset, PW and BW

**Figure S11:**
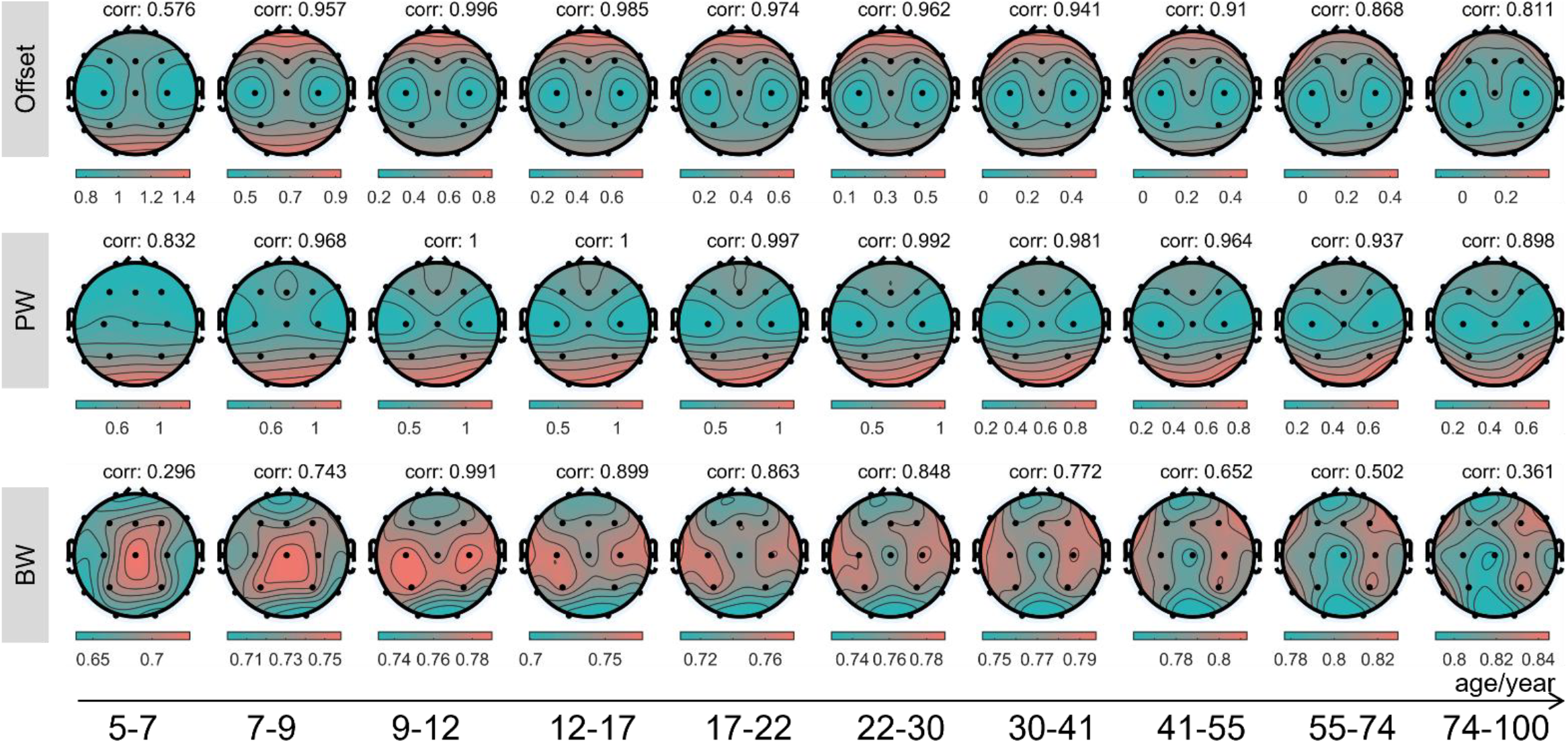
The scalp topography developmental trajectory of Offset, PW and BW. The value of PW and BW are the means of all frequency bands. The ages are divided with log scale and shows mean age of each segment here.

### 4.3 Parameter scalp topography developmental trajectory across different functional regions at alpha band

**Figure S12:**
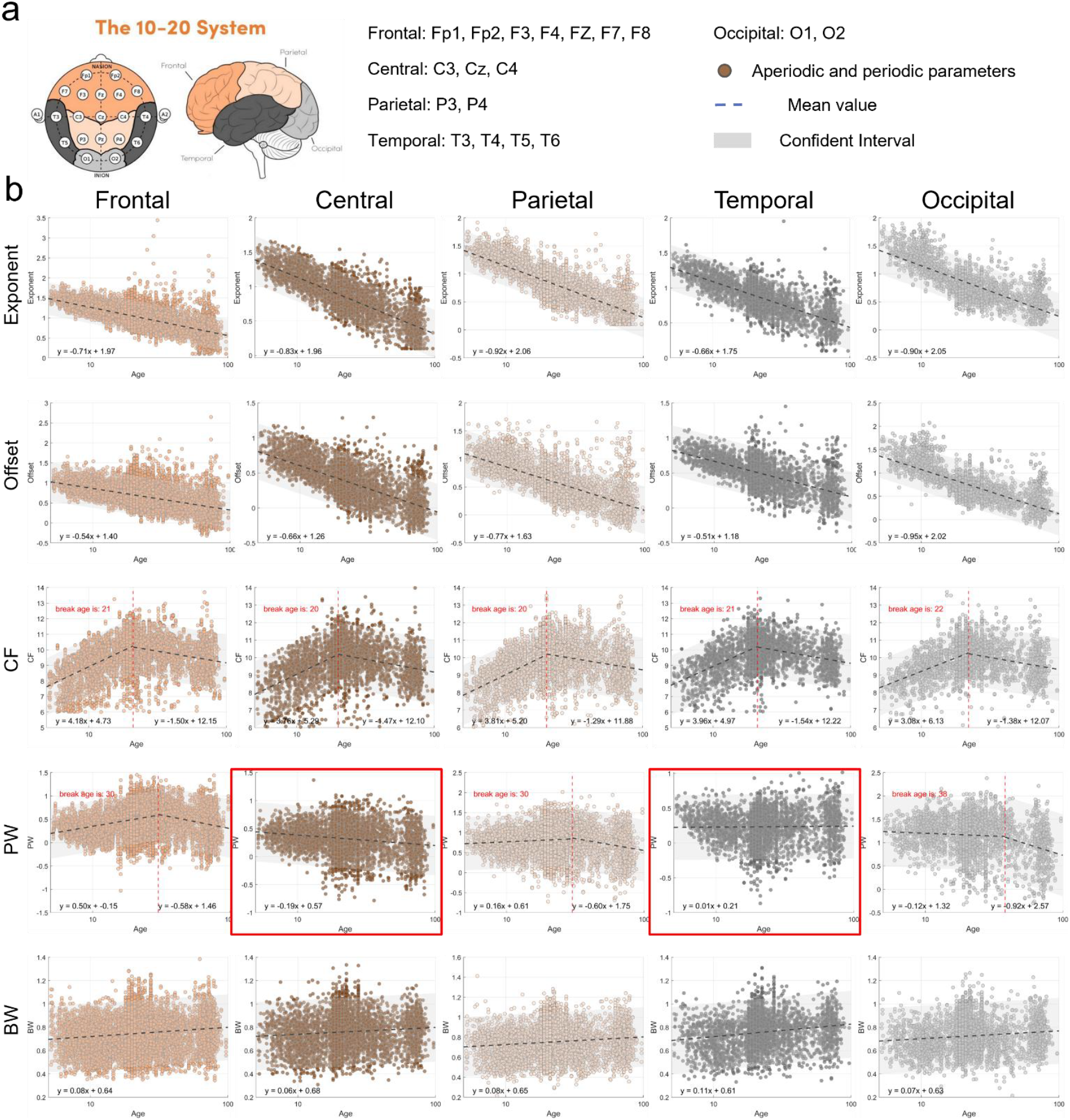
Aperiodic and periodic developmental chart of scalp functional zoning. a: Schematic of functional parcellation of the scalp, divided into five non-overlapping regions: Frontal (includes Fp1, Fp2, F3, F4, FZ, F7, F8), colored in light orange; Central (includes C3, Cz, C4) no color labeled here but in b are colored with dark orange; Parietal (includes P3, P4), colored in light orange; Temporal (includes T3, T4, T5, T6), colored in dark gray; and Occipital (includes O1, O2), colored in light gray. b, Developmental trajectories of aperiodic and periodic parameters across the five functional zones. The dashed gray line represents the piecewise linear fit, with the exact equation noted at the bottom left of each subpanel. Red numbers indicate the estimated breakpoints for two-segment developmental curves. Notably, for power (PW), the developmental trajectory is monotonic in the Central and Temporal regions, potentially due to strong lead field effects in these areas.

## Chapter 5 Spatial distribution of aperiodic and periodic component energy

### 5.1 Spatial distribution of aperiodic and periodic component energy

**Figure S13:**
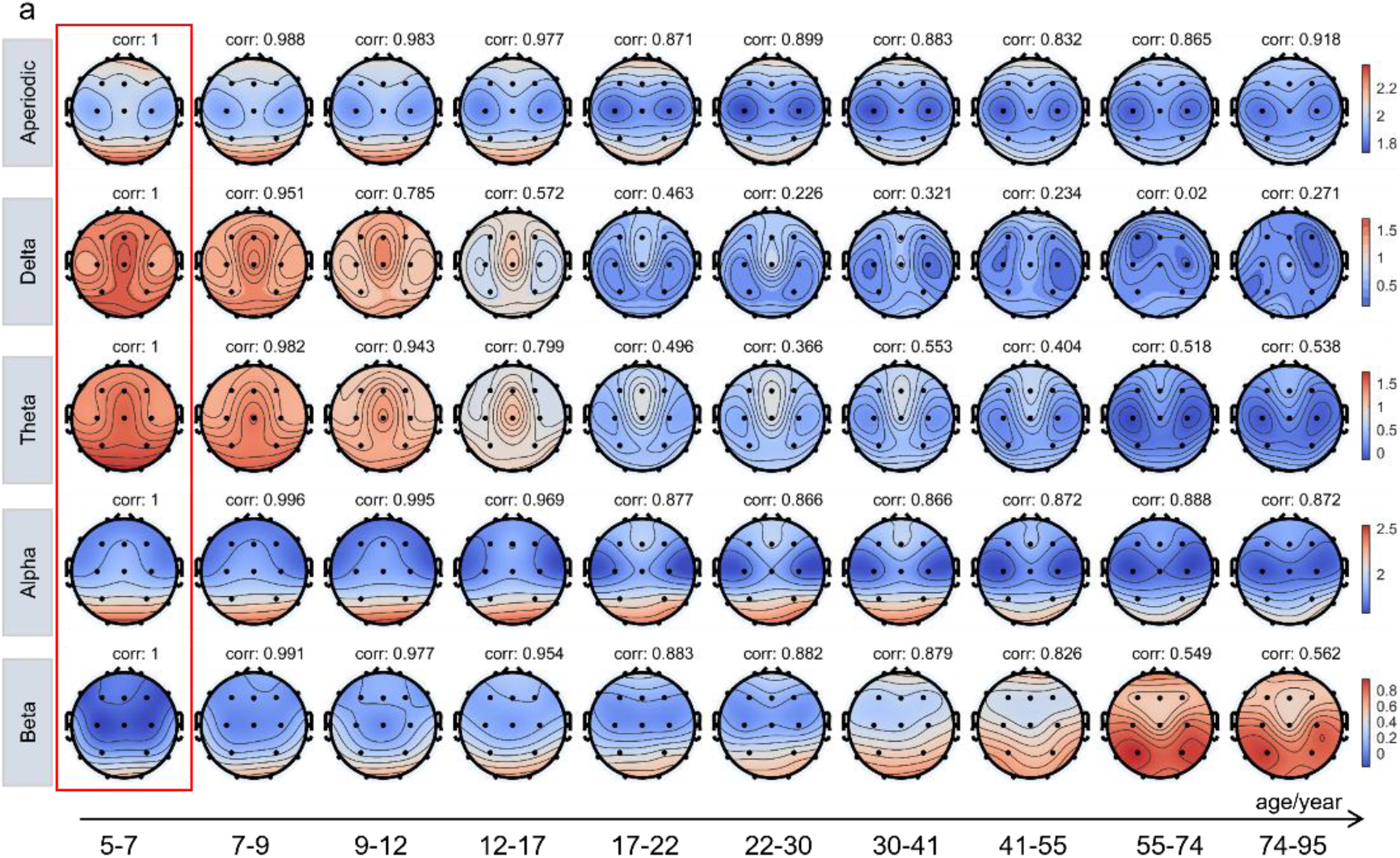
Energy Spatial Distribution on Scalp and Cortex Across Different Components. a: Energy of the aperiodic and periodic components on the scalp. The energy and spatial distribution remain stable in the alpha band, both at the scalp and cortical levels (see correlation values above each subplot, with each value calculated relative to the baseline ages of 5–7 years). For low-frequency bands, energy significantly decreases around age 17, and the spatial similarity on the scalp (relative to the baseline ages of 5–7 years) also drops below 0.5.

### 5.2 Attribution of aperiodic components in different periodic bands

**Figure S14:**
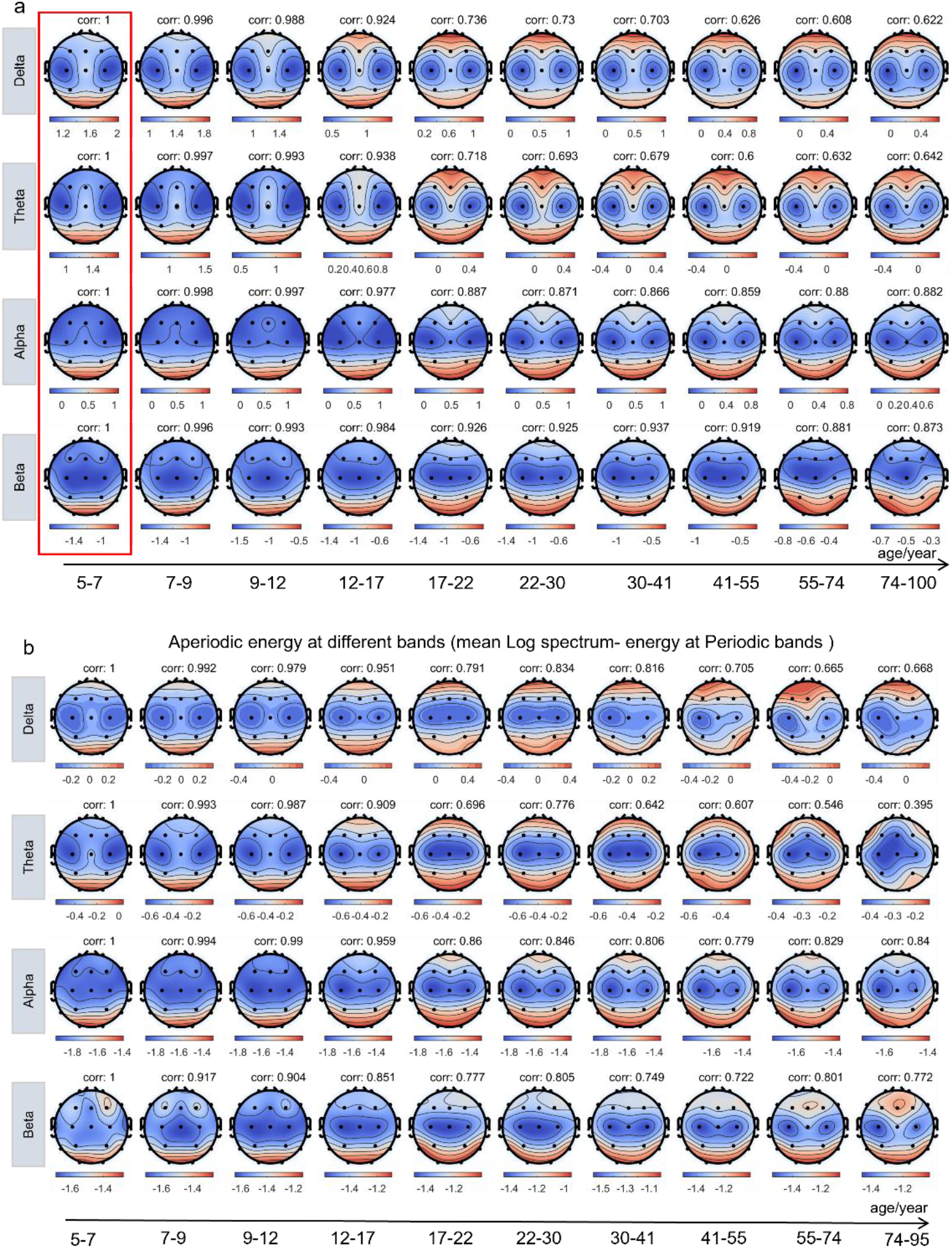
Scalp Topography of the Normative EEG Log-Spectrum Across the Lifespan. Age is segmented on a logarithmic scale to reflect the non-linear trajectory of brain development. The spatial distribution of the EEG log-spectrum remains largely consistent within each frequency band across age groups. However, distinct developmental shifts are observed in the delta and theta bands, where spatial patterns begin to diverge around age 17, showing correlations below 0.8 when compared to the baseline reference group aged 5–7 years.

## Chapter 6 Aperiodic and Periodic component trajectory with other modality

### 6.1 The relations of PW trajectory with other modalities

**Figure S15:**
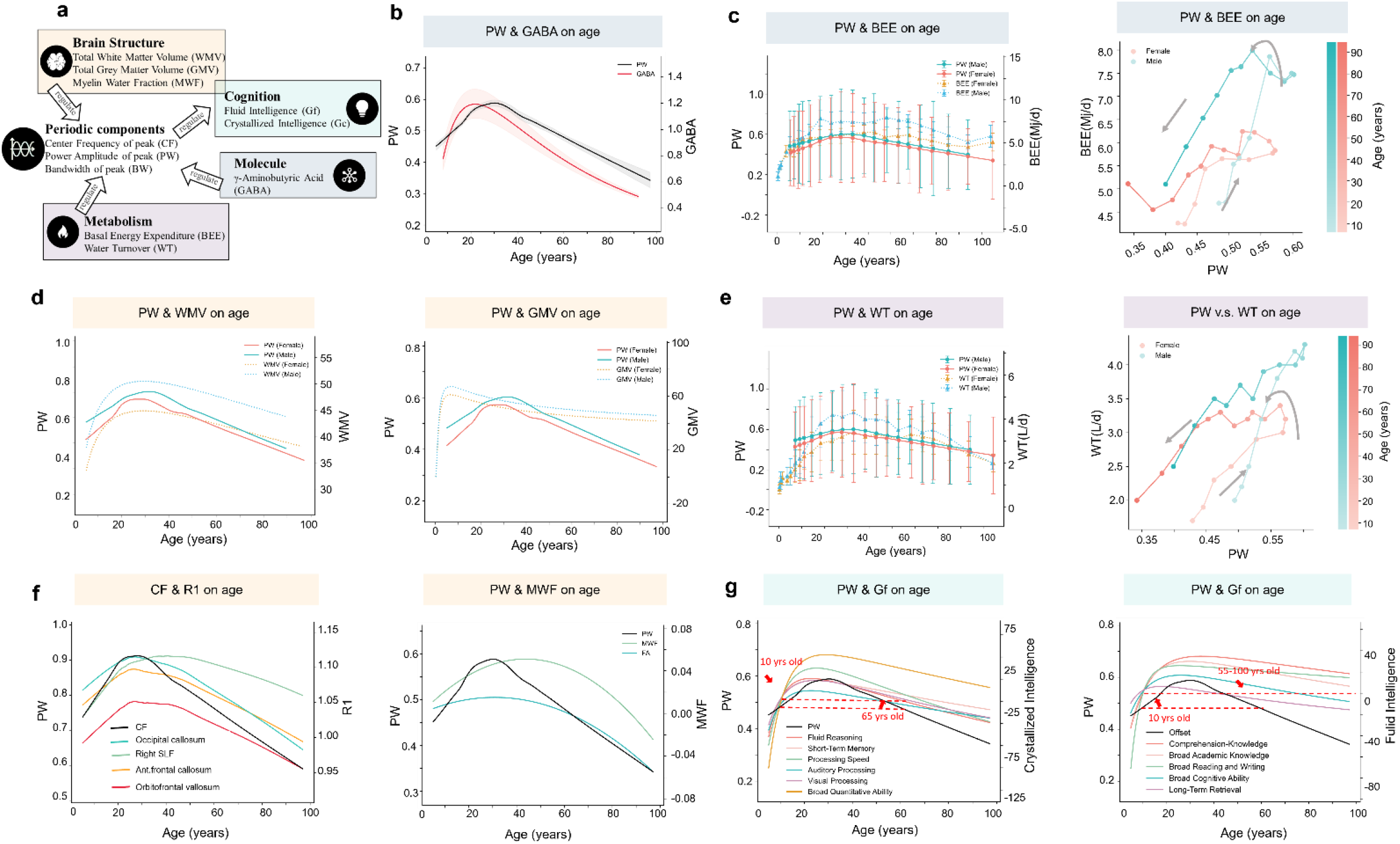
The relations of the developmental trajectory of PW with other modalities.

### 6.2 The relations between Offset with other modalities

**Figure S16:**
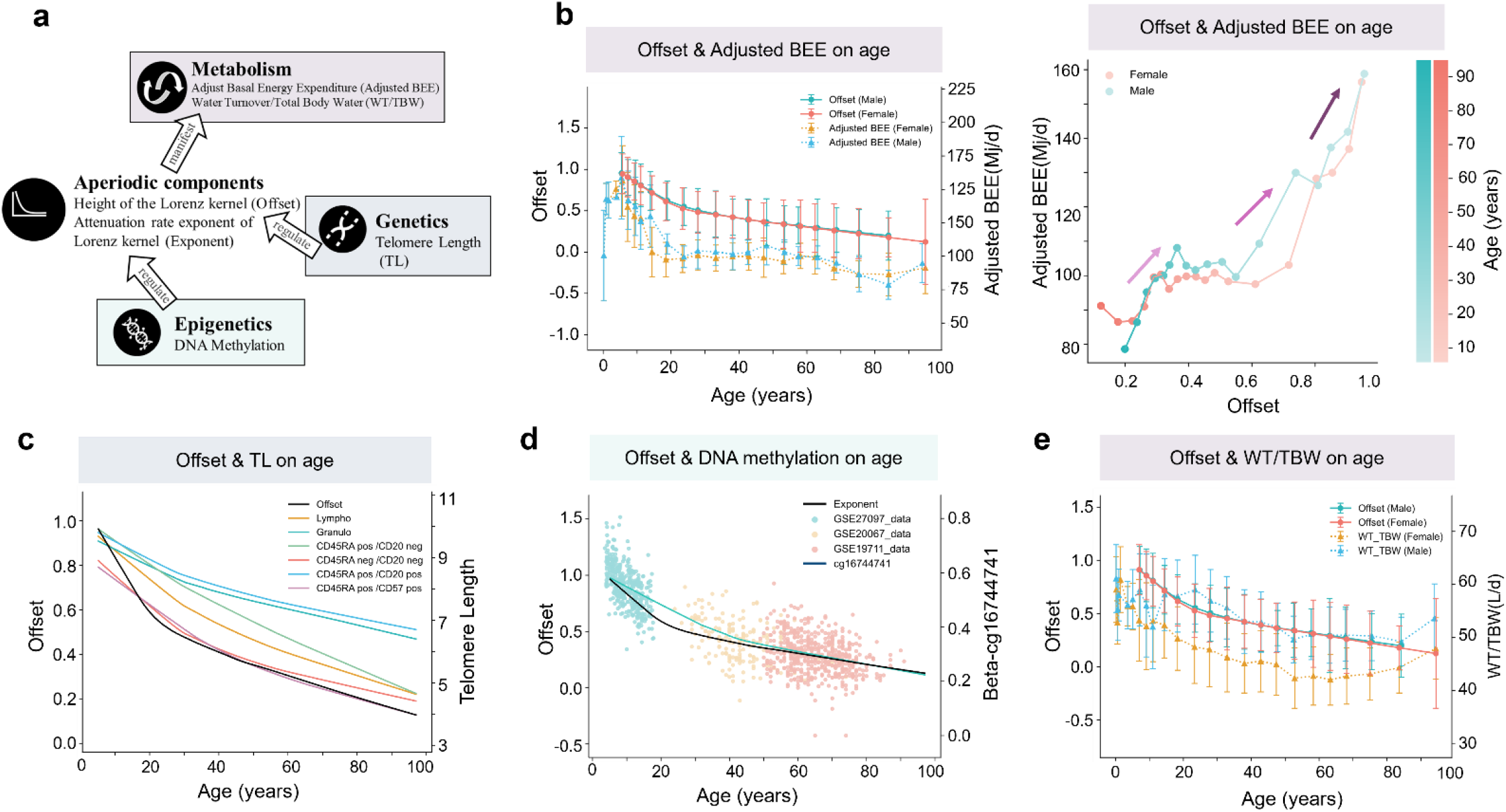
The developmental trajectory relations between Offset and Adjust BEE, WT/TBW, Telomere length and DNA methylation.

### 6.3 The comparison of R1 trajectory between non-parametric regression with parametric

**Figure S17:**
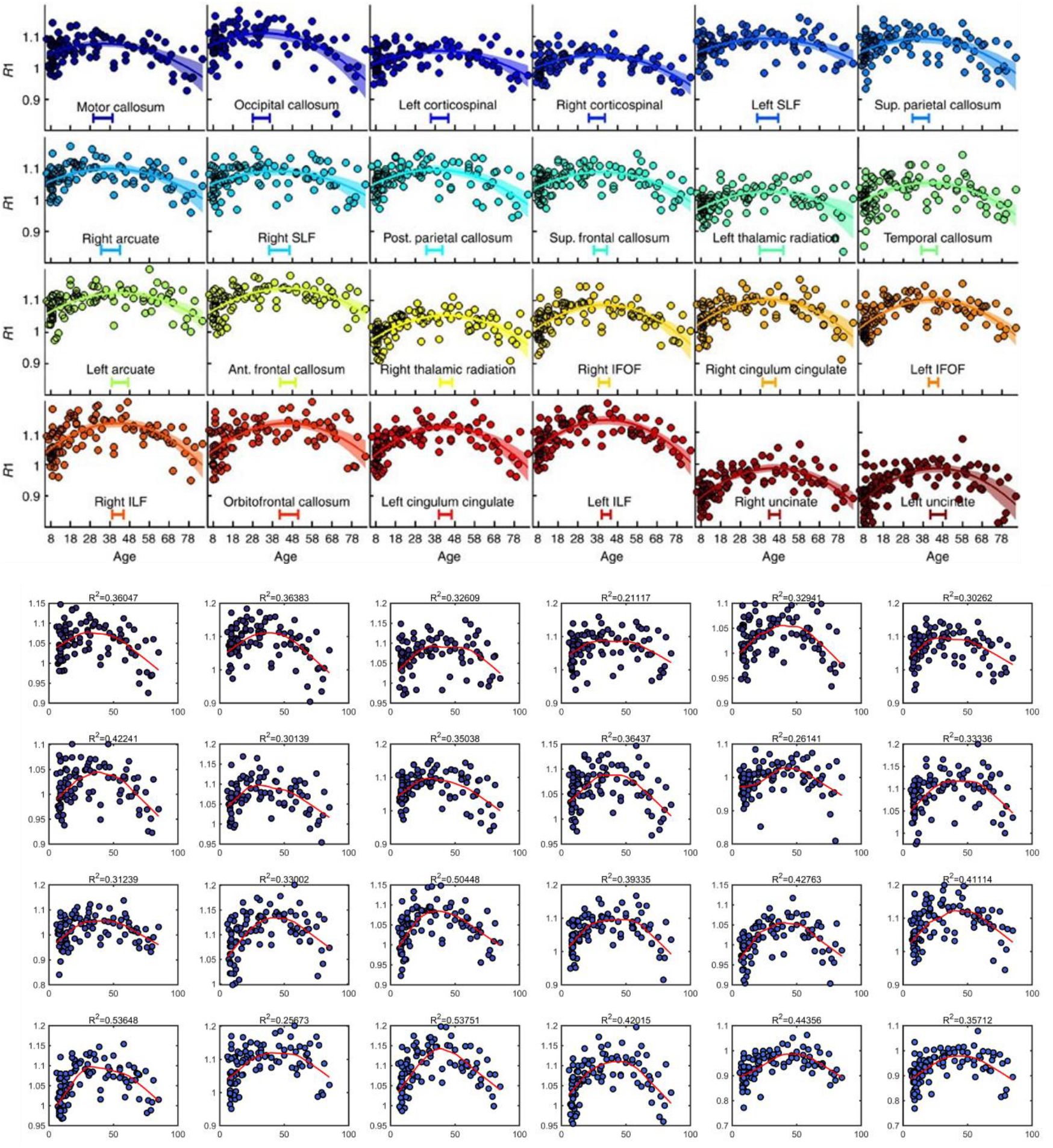
The regression functions of R1 metrics with age. The upper are obtained from the paper (Yeatman et al., 2014) which uses a quadratic function; Lower is a regression curve with Local weighted kernel regression. The results show that not all the region trajectories are symmetric based on the middle period of 38-48 in natural age scale, the non-parametric regression method gives more realistic results than the parametric function here.

## Chapter 7: Standard Age-of-Onset Distributions of DSM-IV/WMH-CIDI Disorders

**Figure S18:**
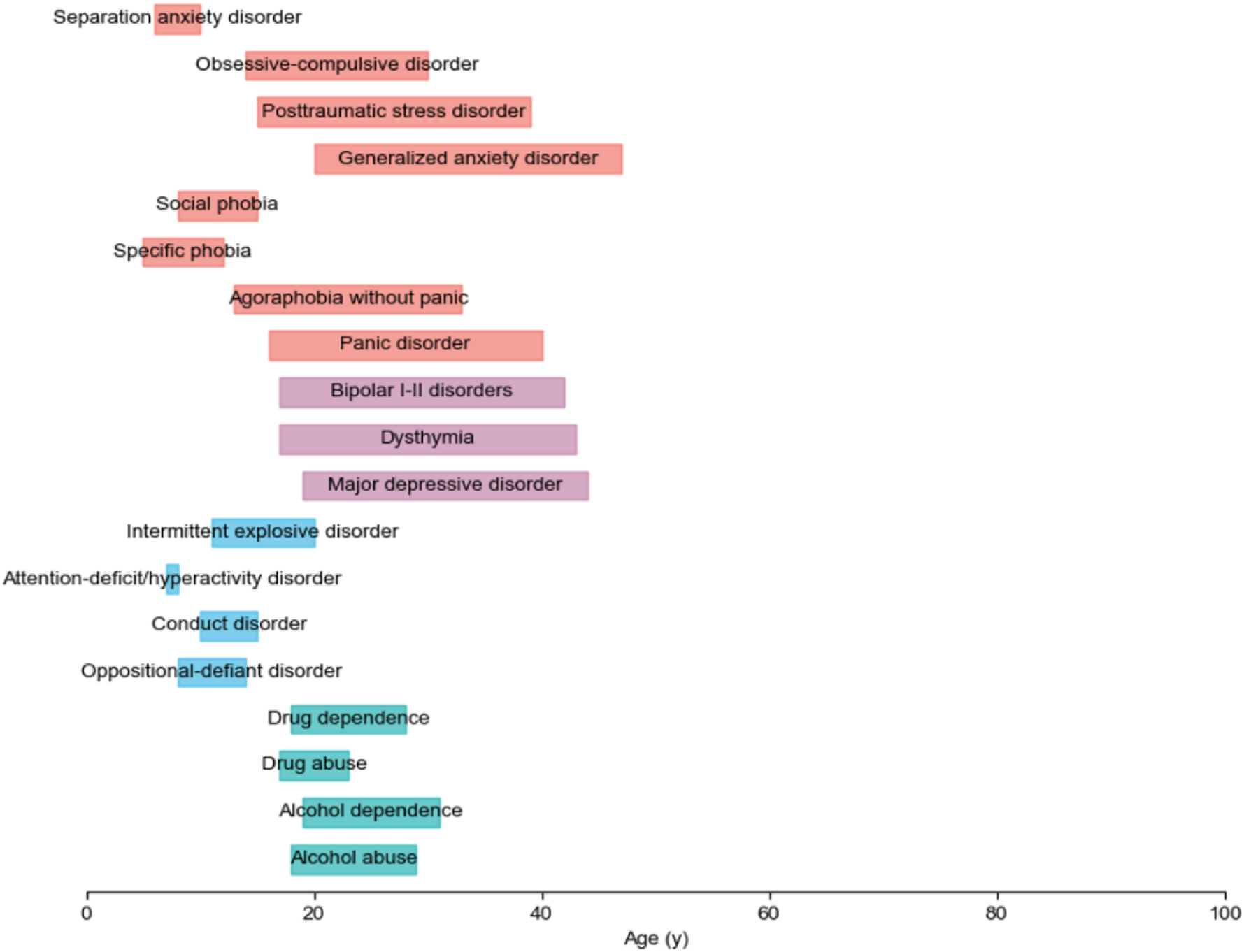
Standardized Age-of-Onset Distributions of DSM-IV/WMH-CIDI Disorders. Data are derived from studies based on the Diagnostic and Statistical Manual of Mental Disorders, Fourth Edition (DSM-IV) and the World Mental Health Survey Composite International Diagnostic Interview (WMH-CIDI). The shaded background indicates the age corresponding to 25% to 75 probability of onset age. The source data used comes from Table 3 of the paper (Kessler et al., 2005).

## Footnotes

1 ⌈·⌉: round up

